# Differential expression and novel activity of FaCCR1 regulate medium-chain ester content and strawberry fruit aroma

**DOI:** 10.1101/2025.09.30.679181

**Authors:** F. Javier Roldán-Guerra, Vítor Amorim-Silva, Joan Jiménez, Ángel Marí-Albert, Rocío Torreblanca, Jorge Ruiz del Río, Miguel A. Botella, Antonio Granell, José F. Sánchez-Sevilla, Cristina Castillejo, Iraida Amaya

**Affiliations:** Centro IFAPA de Málaga, Instituto Andaluz de Investigación y Formación Agraria y Pesquera (IFAPA), 29140 Málaga, Spain; Instituto de Hortofruticultura Subtropical y Mediterránea “La Mayora”, Universidad de Málaga, Consejo Superior de Investigaciones Científicas (IHSM-UMA-CSIC), Campus de Teatinos, 29010 Málaga, Spain; Instituto de Biología Molecular y Celular de Plantas (IBMCP), Consejo Superior de Investigaciones Científicas (CSIC) - Universitat Politècnica de València (UPV), 46022 Valencia, Spain; Unidad Asociada de I+D+i IFAPA-CSIC Biotecnología y Mejora en Fresa, Málaga, Spain

## Abstract

Fruit flavor and aroma are determined by the sugars-to-acid ratio and, most importantly, by the specific profile and abundance of volatile organic compounds (VOCs). While over 550 VOCs have been identified in strawberry fruit, fewer than 20 are believed to be key for aroma, with esters representing the predominant class. A major and stable quantitative trait *locus* (QTL) explaining up to 40% of the phenotypic variation in several medium-chain esters (MCE) was mapped at about 19 Mb in chromosome 6A, through both QTL mapping in a biparental population and a genome-wide association study (GWAS) in a diverse collection. Integration of transcriptomic analyses from bulks contrasting in MCE content and multiple tissues of ‘Camarosa’ identified the cinnamoyl-CoA reductase gene *FaCCR1(6A)* as the most likely candidate underlying this QTL. Subcellular localization experiments indicated that FaCCR1(6A) is localized at the cytosol, and its transient overexpression in fruits significantly increased MCE content, confirming its pivotal role shaping fruit aroma. Although CCR enzymes are known for their involvement in the biosynthesis of monolignols, molecular docking analyses supported the interaction of FaCCR1(6A) with aliphatic acyl-CoA substrates*. In vitro* enzymatic assays further demonstrated that FaCCR1(6A) can catalyze the conversion of aliphatic acyl-CoA substrates to the corresponding medium-chain aldehydes. These findings uncover a previously unrecognized function of FaCCR1(6A) in volatile biosynthesis while providing a mechanistic explanation for the phenotypic differences associated with differential *FaCCR1(6A)* expression. Furthermore, we have developed a predictive KASP assay to facilitate the selection of superior cultivars with enhanced fruit aroma in strawberry breeding programs.

## Introduction

Fruit aroma is a key trait in strawberry (*Fragaria* × *ananassa*) breeding, playing a pivotal role in consumer acceptance and market value. The complex aroma profile of strawberry is determined by the specific composition and relative concentrations of volatile organic compounds (VOCs). While more than 550 VOCs have been identified in cultivated strawberry (Cannon et al., 2015; Ulrich et al., 2018), less than 20 are considered major contributors to perceived flavor and aroma (Jetti et al., 2007; Schieberle & Hofmann, 1997; Ulrich et al., 1997). These compounds belong to diverse chemical classes including esters, alcohols, terpenoids, furans, and lactones, each produced via distinct biosynthetic pathways (Dudareva et al., 2013; Schwab et al., 2008). Esters represent the most abundant and largest family of VOCs, conferring a wide range of fruity notes (Song et al., 2008; Ulrich et al., 2018). While studies often highlight ethyl and methyl esters of butanoic and hexanoic acids as the most abundant compounds (Schwieterman et al., 2014; Ulrich et al., 2018), recent sensory and metabolomic analyses have emphasized the importance of a few medium-chain esters (MCEs) in enhancing consumer preference and increased liking (Fan et al., 2021a; Porter et al., 2023; Schwieterman et al., 2014). These MCEs include butyl esters of acetate, butanoate, and hexanoate, as well as hexyl acetate and octyl esters of butanoate and hexanoate.

The biosynthesis of esters has been increasingly studied in model fruit species such as apple (*Malus domestica*) and strawberry. Esters are produced through the esterification between alcohols and acyl-CoAs, catalyzed by alcohol acyltransferases (AATs), which can utilise a wide range of substrates and contribute to the formation of diverse esters (Aharoni et al., 2000; Beekwilder et al., 2004). Studies in apple, banana, apricot and strawberry have identified specific AAT enzymes that exhibit different preferences for specific acyl-CoA donors and alcohols (Cao et al., 2021; Contreras et al., 2017; Dudareva et al., 2013; Klee & Tieman, 2018; Li et al., 2014; Zhou et al., 2021). In strawberry, the genes *SAAT* and *FaAAT2* were shown to be upregulated during ripening, encoding enzymes with acyl-coA transferase activity and direct involvement in ester biosynthesis (Aharoni et al., 2000; Cumplido-Laso et al., 2012). Both enzymes accept a broad range of alcohol substrates. However, substrate availability, particularly medium-chain acyl-CoAs like hexanoyl-CoA, and alcohol precursors, such as ethanol, methanol, butanol, hexanol or octanol, plays a crucial role in the final ester composition (Aharoni et al., 2000; Beekwilder et al., 2004; Schwab et al., 2008; Souleyre et al., 2005).

The pool of precursor metabolites, including short- to medium-chain fatty acids and their corresponding alcohols, originates from amino acid catabolism and fatty acid metabolism, via either α-oxidation, β-oxidation or the lipoxygenase (LOX) pathway, depending on the fruit and developmental stage (Schwab et al., 2008). Despite advances in understanding ester formation, the regulatory mechanisms and specific enzymes supplying ester precursors remain insufficiently characterized. Genetic analyses have begun to uncover quantitative trait *loci* (QTLs) and candidate genes influencing ester accumulation in apple and strawberry, offering potential tools for targeted breeding (Z. Fan et al., 2022; Rey-Serra et al., 2022; Urrutia et al., 2017; Yang et al., 2023; Zorrilla-Fontanesi et al., 2012). Nonetheless, comprehensive molecular insights into ester biosynthesis and precursor availability are still lacking.

A major and stable QTL controlling the content of several esters was first identified on linkage group 6A more than a decade ago using the strawberry ‘232’ × ‘1392’ biparental population (Zorrilla-Fontanesi et al., 2012). The QTL affected the content of several VOCs, including ten MCEs and two alcohols, whose concentrations were positively correlated in the population. These positively correlated VOCs were butyl, hexyl, octyl, nonyl, decyl and cinnamyl acetates, butyl and octyl hexanoates and butanoates, and two alcohols: 1-octanol and 1-decanol. Advances in sequencing technologies have provided new insights into the allo-octoploid genome of *Fragaria* × *ananassa* (2n = 8x = 56), leading to the obtention of transcriptomes (Fan et al., 2022; Sánchez-Sevilla et al., 2017) and reference genomes with high sub-genome resolution (Hardigan et al., 2021a; Liu et al., 2021). Using these novel tools, QTL for subsets of these MCEs were recently detected at equivalent positions of chromosome 6A by GWAS in germplasm from the University of Florida (Fan et al., 2022) and in independent biparental populations (Rey-Serra et al., 2022).

Given the impact of these MCEs on consumer preference and the stability, as well as the major effect of the 6A QTL, the objective of this study was to identify the underlying gene controlling their variation and to develop marker assays that can be used in marker-assisted selection (MAS). We have combined QTL mapping, association analysis and RNA-Seq to identify candidate genes controlling their variation. Functional validation through transient overexpression in strawberry fruit and *Nicotiana benthamiana* leaves revealed that one key gene, *FaCCR1(6A),* and possibly others, such as *FaOCT4(6A)*, modulate MCE content. Biochemical assays further demonstrated that FaCCR1 can accept butanoyl-, hexanoyl- and octanoyl-CoA to produce biologically relevant concentrations of aldehydes that can be further reduced to alcohols and incorporated into ester biosynthesis, thereby contributing to strawberry aroma. These findings provide mechanistic insight into MCE production in strawberry and establish a foundation for improving fruit aroma through molecular breeding.

## Results

### Narrowing down the 6A QTL controlling fruit MCE content in strawberry

Two previous QTL analyses for MCEs resulted in the detection of equivalent QTL intervals in chromosome 6A that spanned 5-20 cM (Rey-Serra et al., 2022; Zorrilla-Fontanesi et al., 2012), comprising several hundred annotated genes. Similarly, GWAS identified a large region (14 Mb) in the same 6A chromosome sharing significant Single Nucleotide Polymorphism (SNPs) for six of these esters (Fan et al., 2022). To increase the resolution of QTL mapping, we re-analyzed VOC data for the 10 esters (9 MCEs and cinnamyl acetate) and the two alcohols from Zorrilla-Fontanesi et al., 2012 using a higher resolution ‘232’ × ‘1392’ genetic map (Sánchez-Sevilla et al., 2015). As previously described, mean values in the three seasons for the 12 VOCs (Supplementary Table S1) displayed significant (*p-value* < 0.05) correlations among them, with several strong positive correlation coefficients (*r* = 0.46-0.93; Fig. 1A). This population exhibited extensive phenotypic diversity, with some F_1 l_ines displaying extremely low MCE levels (<3 relative concentration, rc) and others showing considerably high levels (>25 rc), while the two parental lines were closer to the mean population (Supplementary Table S1). Principal Component Analysis (PCA) further corroborated these observations, organizing VOC vectors and high MCE lines to the right of principal component 1, which accounted for nearly 70% of variance. In contrast, the parental lines were positioned near the origin of the plot (Fig. 1B). New QTL analysis in the ‘232’ × ‘1392’ population detected the same QTL in LG 6A for all VOCs, which remained stable across different years except for hexyl acetate, which was detected below the threshold during one season (Fig. 1C; Supplementary Table S2). For MCEs, the phenotypic variance explained (PVE) ranged from 17.10 to 57.90% depending on the volatile and the season. For the two alcohols, the PVE ranged from 13.80 to 31.70%. The average PVE value over the three seasons for butyl acetate, butyl butanoate, octyl butanoate and octyl hexanoate ranged from 27 to 38%, which are important contributors to strawberry flavor, consumer liking and increased sweetness perception (Fan et al., 2021a; Porter et al., 2023; Schwieterman et al., 2014). In all cases, the allelic effects indicated that the ‘232’ and ‘1392’ parental lines were heterozygous, in agreement with their overall average MCE content. The confidence intervals for the analyzed QTL span a large chromosomal region of approximately 30 cM. A common region overlapping most QTL at least one season spanned an interval of 10 cM, from 12 to 22 cM (Fig. 1C; Supplementary Table S3). The equivalent region in the *Fragaria* × *ananassa* ‘Royal Royce’ v1.0 reference genome spanned the interval 18.57-21.49 Mb and enclosed 373 genes (Supplementary Table S4).

**Figure 1.**
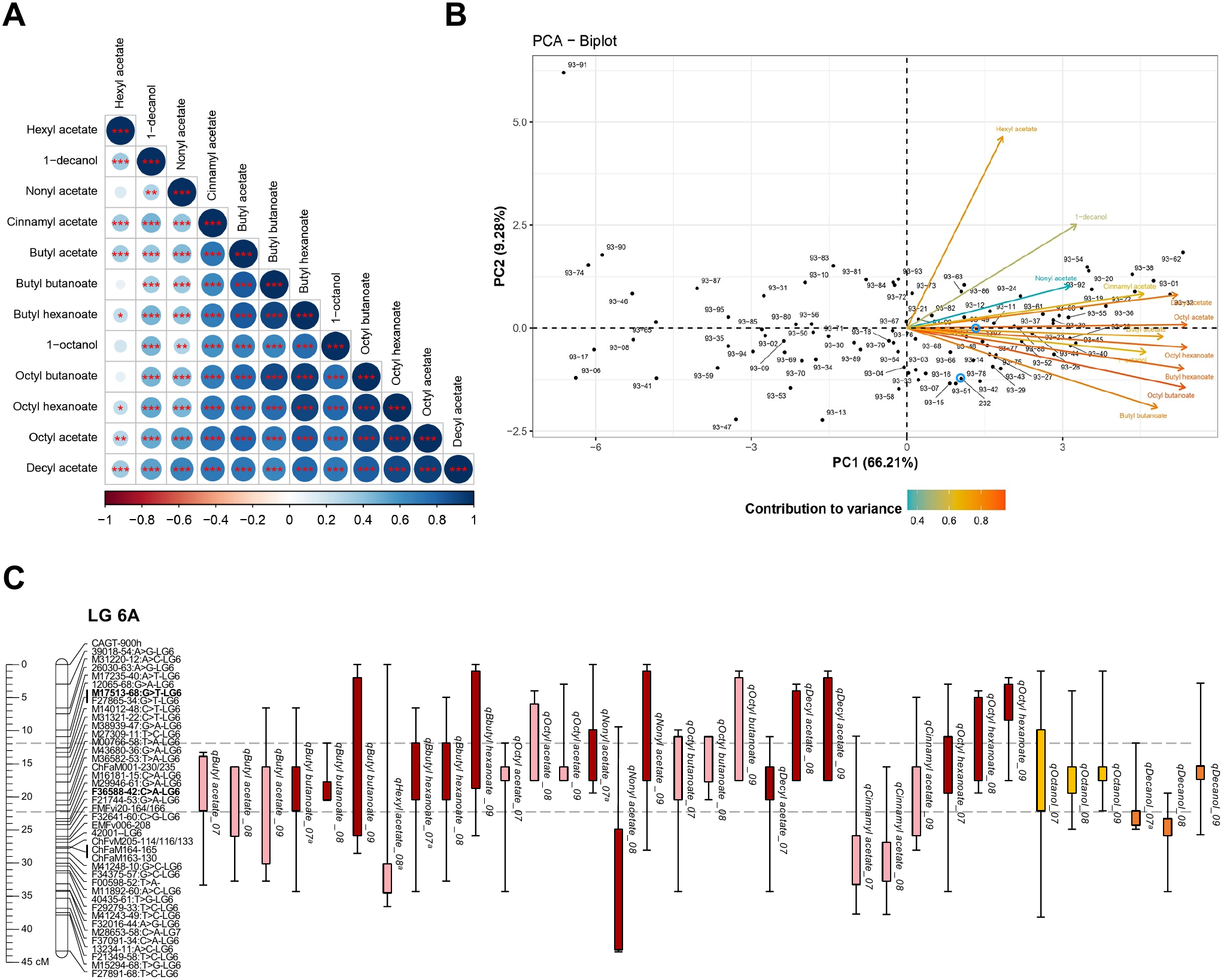
QTL analysis of medium-chain esters (MCE) and two alcohols in the ‘232’ ξ ‘1392’ population. **A)** Correlation matrix among the twelve volatile organic compounds (VOC) quantified in the population. Each circle represents Pearson correlation coefficient (*r*) in a color scale from red (negative correlation) to blue (positive correlation). * *p*-value ≤ 0.05, ** *p*-value ≤ 0.01, *** *p*-value ≤ 0.001. **B)** Principal Component Analysis (PCA) biplot. Each F_1_ line is shown by a dot (parental lines highlighted in blue), while arrows represent VOC effects. The variance explained by each principal component (PC) is displayed in the axis and VOCs are color-coded according to their contribution to this variance. **C)** Position of QTL controlling MCE at linkage group (LG) 6A in three consecutive years (2007–2009) using interval mapping (IM). QTL are shown as colored boxes (1-LOD interval) to the right of the LG, with error bars showing the 2-LOD interval. Dashed horizontal lines define a common region overlapping the 1-LOD QTL intervals at least one season.

As QTL mapping did not further refine this interval, we conducted a genome-wide association study (GWAS) for MCE content using 124 accessions from the genetically diverse IFAPA GWAS collection (Muñoz et al., 2024). Fruit MCE content in this set displayed a broad phenotypic variation, ranging from 1.80 to 55.88 rc, with a population mean of 11.50 rc (Supplementary Table S5). PCA revealed a clustering of MCEs towards the right side of PC1, similar to the pattern observed in the ‘232’ × ‘1392’ mapping population, although the proportion of variance explained (31.11%) was lower (Supplementary Fig. S1). Genetic diversity analyses of this population identified 6 subpopulations based on their genetic background, with a low pairwise fixation index (*F_ST_* _=_ 0.13; Muñoz et al., 2024). Notably, no differentiation in MCE content was observed between older accessions (group 3, blue color) and recent Mediterranean and Californian accessions (group 1, red color), indicating substantial phenotypic diversity regardless of geographic origin or breeding history (Supplementary Fig. S2). We have analyzed linkage disequilibrium (LD) decay using 40,808 SNPs, revealing a genome-wide moderate decline over a 0.8 Mb distance in this population (Supplementary Fig. S3). GWAS using the BLINK multi-*loci* model only identified a total of 5 significant marker-trait associations (*p*-value < 0.05/no. SNPs) corresponding to three MCEs, the aromatic ester cinnamyl acetate, which was also detected in the ‘232’ × ‘1392’ population, and to the total MCE content (Supplementary Fig. S4;

Supplementary Table S6). As observed in the ‘232’ × ‘1392’ population, each SNP detected by GWAS explained ∼35% of the phenotypic variance. A regional association plot showing the lead SNPs for the four esters and for the total MCE content is shown in Fig. 2A. LD analysis revealed a haploblock in strong LD (*D’* > 0.8) spanning a rather large region of approximately 2 Mb (‘Royal Royce’ genome v1.0 Chr_6A: 18,609,279-20,573,409 bp), delimited by the 50K FanaSNP markers AX-184595454 and AX-166507478 (Fig. 2B). This region encloses 245 annotated genes (Supplementary Table S7) and, importantly, it is enclosed within the QTL interval previously detected in the ‘232’ × ‘1392’ population (Supplementary Table S4). Taken together, these results define a specific region on chromosome 6A, hereafter designated *qMCE-6A*, as a major genetic determinant of MCE fruit content.

**Figure 2.**
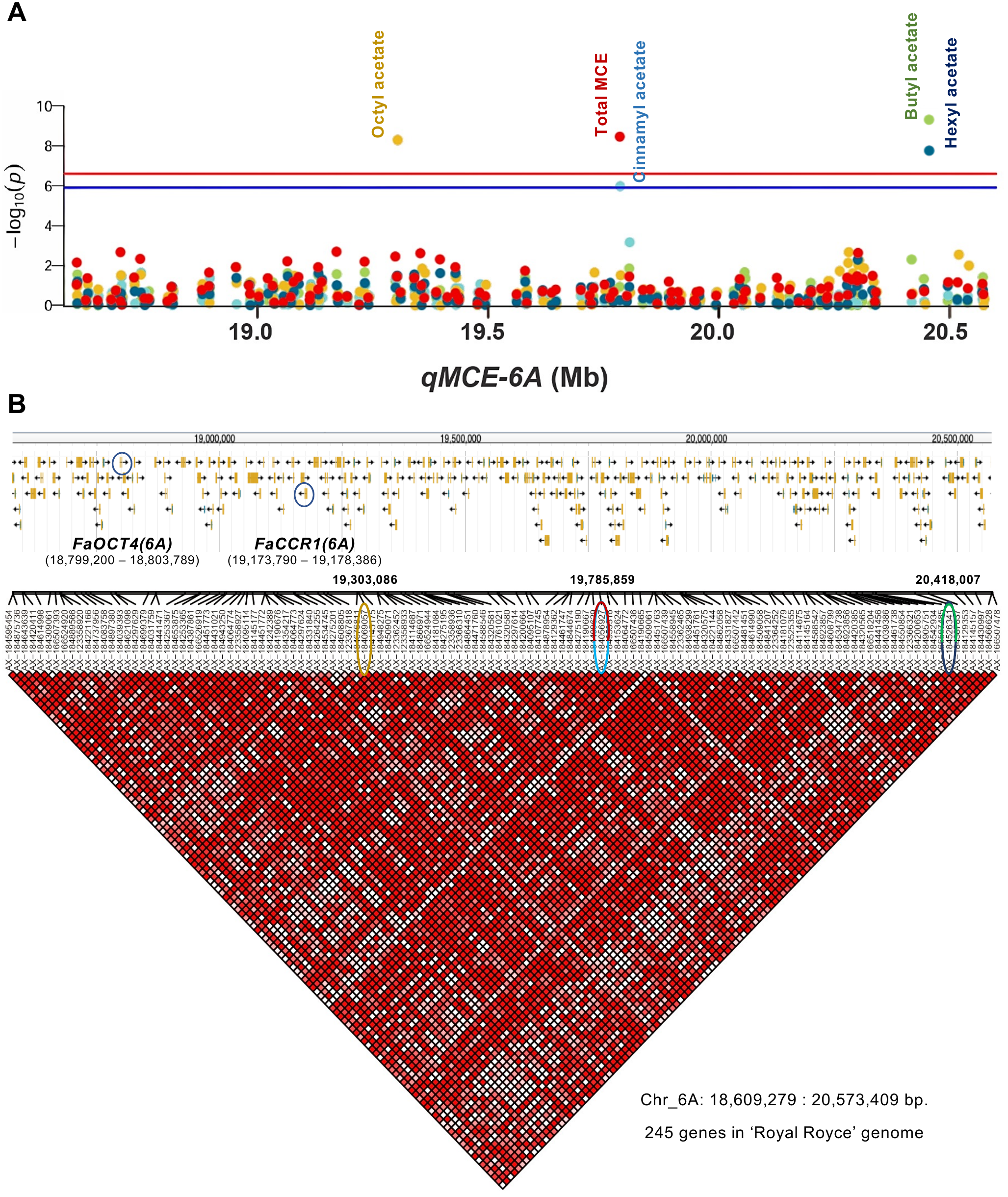
Significant SNP-trait associations in chromosome 6A for MCE content in fruit detected by GWAS. **A)** Regional Manhattan plot within the *qMCE-6A* QTL showing the 5 significant SNP-trait associations detected using the 124 strawberry accessions. SNPs are color coded for each compound. Red and blue horizontal bars denote the significance threshold (−log_10_ (0.01/total SNPs) and −log_10_ (0.05/total SNPs), respectively). **B)** Haplotype block structure of *qMCE-6A*. The relative physical position of each SNP is given above the plot and significant SNPs detected by GWAS are highlighted with circles colored according to the corresponding compound. The *FaCCR1(6A)* and *FaOCT4(6A)* candidate genes are also indicated with a blue circle. The pairwise linkage disequilibrium (LD) value (*D’*) is represented by a scale from white (0, no LD) to dark-red (1, complete LD).

### Transcriptomic analyses unveiled *FaCCR1(6A)* and *FaOCT4(6A)* as candidate genes for MCE biosynthesis

To narrow down the number of candidate genes and to identify molecular determinants controlling these important esters, we performed RNA-Seq analysis to identify differentially expressed genes (DEGs) between bulked ripe fruit pools from F_1 l_ines in the ‘232’ × ‘1392’ population exhibiting contrasting MCE content. The F_1 l_ines used for pool generation are detailed in Supplementary Table S1 and the difference in MCE mean content is shown in Supplementary Fig. S5A. Transcriptomes obtained through Illumina sequencing generated an average of 22 million high-quality reads per sample, with 96.23% of them successfully mapped to the *F*. × *ananassa* ‘Royal Royce’ genome. PCA revealed a notable distinction between high and low MCE samples, suggesting appreciable transcriptomic differences in sample pools (Supplementary Fig. S5B). We further confirmed these findings identifying a total of 823 differentially expressed genes (|log_2(_fold change)| ≥ 1; *p*-adjusted < 0.05), of which 312 were upregulated in the high MCE pool and 511 were downregulated (Fig. 3A; Supplementary Table S8). The top ten most up- and downregulated genes in the pool of fruit with high MCE content are shown in Table 1.

**Figure 3.**
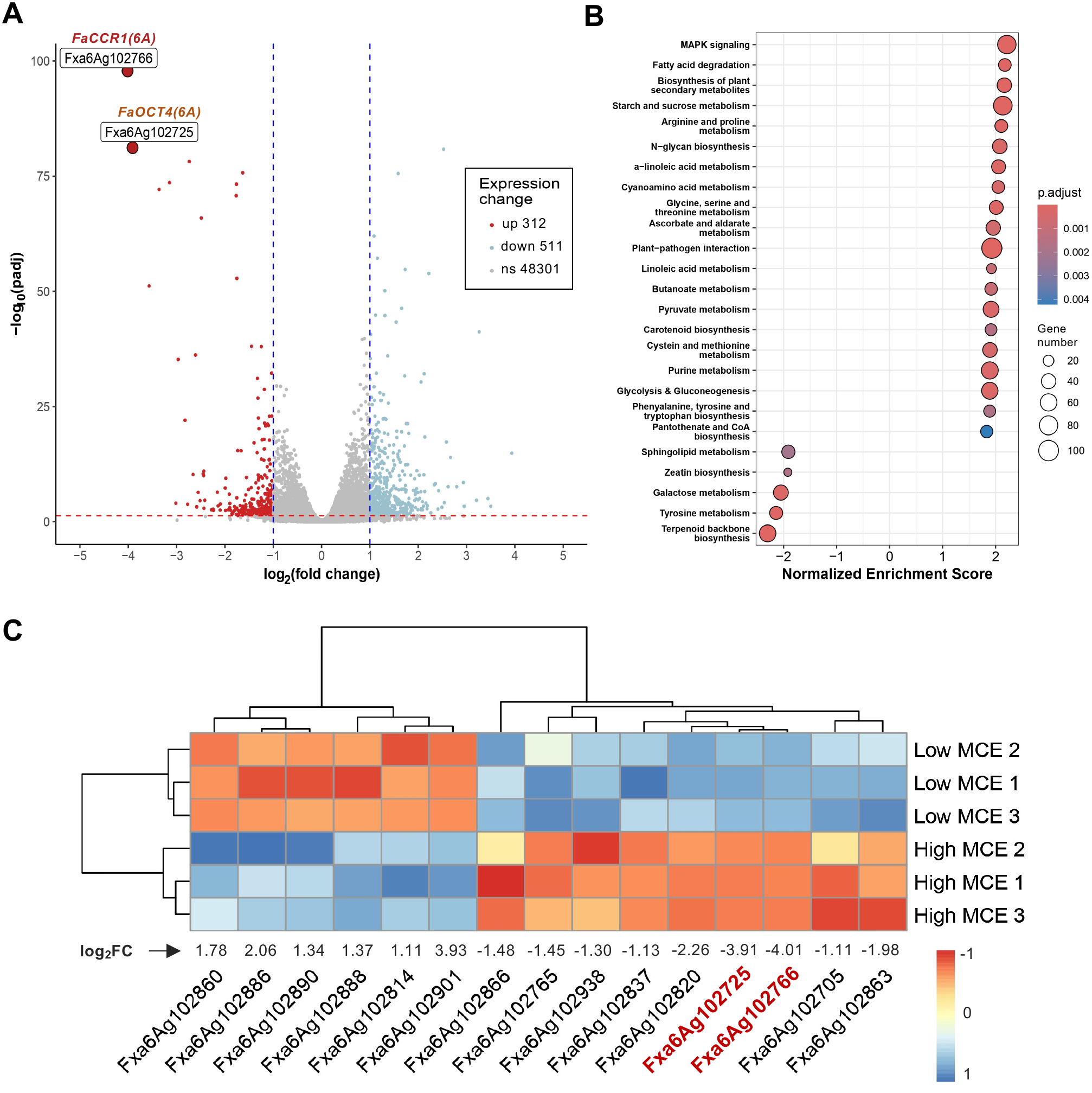
Transcriptomic analysis by RNA-Seq in fruits of F_1_ lines from the ‘232’ × ‘1392’ population contrasting in medium-chain ester content. **A)** Volcano plot of differentially expressed genes (DEGs), showing the –log_10_ adjusted *p*-value (y-axis) versus the log_2_ fold change between pools (x-axis). Upregulated genes (*p*-adj ≤ 0.05, log_2_FC ≤ −1) are shown in red, while blue color denotes down-regulation (*p*-adj ≤ 0.05, log_2_FC ≥ 1), and non-significant genes in grey. The numbers of up-, down-, and non-significant (ns) genes are indicated. *FaCCR1(6A)* (*Fxa6Ag102766*) and *FaOCT4(6A)* (*Fxa6Ag102725*) are highlighted as the most strongly up-regulated DEGs. **B)** KEGG enrichment analysis showing the 25 significantly enriched pathways *(p*-adj *<* 0.05), ranked by Normalized Enrichment Score (NES). Enriched pathways are sorted by their NES, which reflects the overrepresentation degree of a gene set normalized for gene set size and multiple testing. Negative NES values indicates downregulation, while positive values indicate upregulation. **C)** Gene expression (Z-Score scaled) heatmap of DEGs within the *qMCE-6A* QTL across samples. Relative expression values range from blue (down-regulation) to red (up-regulation). Samples from contrasting MCE content pools and genes were grouped by hierarchical clustering. Log_2_ fold change between pools is displayed above each gene ID. *FaCCR1(6A)* (*Fxa6Ag102766*) and *FaOCT4(6A)* (*Fxa6Ag102725*) are highlighted in red.

**Table 1.** Top ten up-regulated and down-regulated DEGs in the high ester pool of F1 lines from the ‘232’ × ‘1392’ population. Expression levels are shown in fragments per kilobase million (FPKM). Gene numbers, position and annotated function are referenced following the ‘Royal Royce’ reference genome.

| Gene | Locus (bp) | Predicted function | log2 fold change | padj-value | High esters (FPKM) | Low esters (FPKM) |
|---|---|---|---|---|---|---|
| Fxa6Ag102766 | Chr_6A:19173564-19178883 | Cinnamoyl CoA reductase (*FaCCR1(6A)*) | -4.013 | 1.41E-98 | 995.855 | 61.662 |
| Fxa6Ag102725 | Chr_6A: 18799173-18803905 | Organic cation/carnitine transporter (*FaOCT4(6A)*) | -3.91 | 6.37E-82 | 39.204 | 2.608 |
| Fxa6Bg102491 | Chr_6B:20325381-20330329 | Sugar transporter (organic cation – NCBI) | -3.568 | 6.43E-52 | 2.19 | 0.185 |
| Fxa6Cg102432 | Chr_6C:19521235-19532172 | Cinnamoyl CoA reductase | -3.363 | 7.37E-73 | 55.04 | 5.349 |
| Fxa6Bg102528 | Chr_6B:20665682-20670049 | Cinnamoyl CoA reductase | -3.148 | 2.37E-74 | 43.351 | 4.891 |
| Fxa3Cg100459 | Chr_3C:2844078-2845202 | - | -3.016 | 9.62E-06 | 1.148 | 0.113 |
| Fxa1Dg202056 | Chr_1D:14706312-14712105 | Major facilitator superfamily domain-containing protein 10 | -2.989 | 0.0453 | 0.337 | 0.028 |
| Fxa6Dg102347 | Chr_6D:18447826-18452684 | Sugar transporter (organic cation – NCBI) | -2.967 | 5.99E-36 | 1.878 | 0.24 |
| Fxa6Cg102410 | Chr_6C:19336981-19343714 | Organic/cation carnitine transporter | -2.825 | 9.55E-23 | 0.73 | 0.102 |
| Fxa7Ag201153 | Chr_7A:9309906-9311671 | Expansin B2 | -2.772 | 1.60E-05 | 0.708 | 0.103 |
| Fxa7Cg102670 | Chr_7C:20754467-20755019 | AP2/ERF domain superfamily | 2.583 | 4.83E-18 | 1.718 | 10.297 |
| Fxa1Ag100837 | Chr_1A:4643872-4645114 | AP2 domain | 2.619 | 2.43E-08 | 0.612 | 3.236 |
| Fxa7Dg102482 | Chr_7D:20310822-20311374 | Dehydration-responsive element-binding protein 1D | 2.67 | 1.09E-14 | 1.168 | 7.439 |
| Fxa5Cg201406 | Chr_5C:9283596-9284865 | ARM repeat superfamily protein | 2.798 | 2.43E-08 | 0.224 | 1.462 |
| Fxa3Ag101807 | Chr_3A:11481577-11483464 | Concanavalin A-like lectin protein kinase family protein | 2.934 | 3.10E-09 | 0.11 | 1.029 |
| Fxa2Dg202453 | Chr_2D:20978786-20979371 | AP2/ERF domain | 3.206 | 2.27E-07 | 0.227 | 1.804 |
| Fxa4Ag100300 | Chr_4A:2456154-2485680 | DNA binding protein | 3.262 | 5.92E-42 | 0.656 | 5.915 |
| Fxa2Bg200484 | Chr_2B:3869777-3870916 | AP2 domain class transcription factor | 3.442 | 9.70E-09 | 0.1 | 0.969 |
| Fxa5Cg200122 | Chr_5C:785857-786627 | Transcription factor ERF19 | 3.502 | 4.50E-04 | 0.492 | 4.659 |
| Fxa6Ag102901 | Chr_6A:20228619-20233236 | Putative disease resistance protein RGA3 isoform X2 | 3.936 | 1.34E-15 | 0.218 | 3.334 |

Kyoto Encyclopedia of Genes and Genomes (KEGG) pathway enrichment analysis identified a set of differentially enriched pathways (*p*-adjusted < 0.05) which might be related to ester biosynthesis (Fig. 3B; Supplementary Table S9). They were further examined through integrated visualization of expression data, revealing an upregulated metabolic flux towards fatty acid degradation (fve00071; Supplementary Fig. S6A), enclosing a group of genes annotated as long-chain acyl-CoA synthetases/ligases (e.g., *Fxa3Ag100873*, EC 6.2.1.3), as well as the majority of enzymes in the main cycle of β-oxidation: acyl-CoA oxidases (e.g., *Fxa6Cg103740*, EC 1.3.3.6), enoyl-CoA hydratases (e.g., *Fxa4Ag101989*, EC 4.2.1.17), and ketoacyl-CoA esterases/thiolases (e.g., *Fxa4Ag100029*, EC 2.3.1.16). Moreover, lines with high MCE content exhibited increased expression of aldehyde dehydrogenases (e.g., *Fxa5Dg200422*, *Fxa5Cg200421*, or *Fxa1Cg100763*, EC 1.2.1.3) and downregulation of a number of alcohol dehydrogenases (e.g. *Fxa2Ag102054* or *Fxa7Dg102898,* EC 1.1.1.1; Supplementary Fig. S6A; Supplementary Table S9). However, other alcohol dehydrogenases, such as *Fxa2Bg201327 or Fxa7Ag201111*, were up-regulated. A similar trend was found in the biosynthesis and mobilization of CoA (fve00770), driven by genes encoding dephospho-CoA kinases and CoA transferases (e.g., *Fxa2Bg202381*, EC 2.7.1.24 and *Fxa6Dg101588*, EC 2.7.8.-), or aldehyde dehydrogenases (e.g., *Fxa1Ag100799*, EC 1.2.1.3) as in the previous pathway, though the latter now related to β-alanine metabolism (Supplementary Fig. S6B). Within butanoate metabolism (fve00650) no differential expression was detected in medium-chain acyl-CoA synthetases/ligases (e.g., *Fxa4Ag101927*, EC 6.2.1.2), although differential upregulation of acyl-CoA dehydratase activity was found (e.g., *Fxa4Ag101989*, EC 4.2.1.17), which is consistent with the upregulation of the β-oxidation cycle (Supplementary Fig. S6C; Supplementary Table S9). In addition, we observed a general downregulation in the terpenoid biosynthesis (fve00900; Supplementary Fig. S6D) and an upregulation of the last part of the linoleic acid metabolism towards the formation of jasmonic acid (fve00592; Supplementary Fig. S6E). However, the first enzyme in the pathway, lipoxygenase (1.13.1112), was downregulated in F_1 l_ines with high MCE content.

A total of 15 DEGs were located in the common *qMCE-6A* confidence interval detected in both the ‘232’ × ‘1392’ population and the GWAS collection, with 9 of them being upregulated in the high MCE pool (Fig. 3C; Supplementary Tables S4 and S7). Given that esters accumulate in ripe fruit (Fait et al., 2008; Song et al., 2008), the underlying gene controlling MCE content is expected to be upregulated during fruit ripening, with greater expression in the ripe receptacle. Therefore, we analyzed the expression of these 15 candidate genes using previously published RNA-Seq data from different tissues of ‘Camarosa’ (Sánchez-Sevilla et al., 2017) here mapped to the ‘Royal Royce’ genome. Three out of the 15 genes were upregulated during receptacle ripening (Supplementary Fig. S7A). Among them, *Fxa6Ag102938* exhibited relatively low gene expression in ripe fruit, with FPKM values below 2 (Supplementary Table S8). In contrast, the other two genes (*Fxa6Ag102766* and *Fxa6Ag102725*) showed high expression levels in the ripe receptacle and were strongly upregulated during the final stages of ripening. *Fxa6Ag102766* was specifically upregulated in the receptacle during fruit ripening, while *Fxa6Ag102725* was upregulated both in the achene and the receptacle (Supplementary Fig. S7B and S7C). Remarkably, *Fxa6Ag102766*, encoding a cinnamoyl CoA reductase (EC 1.2.1.44), hereafter *FaCCR1(6A)*, and *Fxa6Ag102725*, with similarity to an organic cation/carnitine transporter 4, hereafter *FaOCT4(6A)*, showed the strongest upregulation in lines with high MCE content (Fig. 3A; Table 1; Supplementary Table S8). Notably, the expression level of *FaCCR1(6A)* in the high MCE pool was 25-fold higher than that of *FaOCT4(6A)*, with FPKM values of 995 and 39, respectively (Table 1). Among the four *FaCCR1* and *FaOCT4* homoeologs, *FaCCR1(6A)* and *FaOCT4(6A)*, from the *F. vesca*-derived subgenome A, where *qMCE-6A* is mapped, exhibited the highest expression. The coordinates or *FaOCT4(6A)* and *FaCCR1(6A)* on chromosome 6A of ‘Royal Royce’ are 18,799,173 - 18,803,905 bp and 19,173,564 - 19,178,883 bp, respectively (Fig. 2B).

RNAseq reads from both *FaCCR1(6A)* and *FaOCT4(6A)* were analyzed to identify coding-sequence polymorphisms occurring at different frequencies in the high- and low-ester pools that could potentially affect the enzymatic activity of the encoded proteins. In *FaCCR1(6A)*, a single SNP was identified at position 462 (c.462C/T). All reads from the high-esters pool carried the C allele at this position, whereas the low-ester pool showed an approximately equal proportion of C and T alleles (50% each). However, this SNP results in a synonymous substitution and does not alter the encoded amino acid sequence. The *FaOCT4(6A)* coding sequences from the high-and low-ester pools differed at two nucleotide positions: c.254G/A and c.264A/C, of which only the former resulted in an amino acid substitution (p.Gly85Asp).

While the function of FaOCT4 is unknown, it might facilitate the transport of small organic compounds (Küfner & Koch, 2008). In contrast, FaCCR1 has been broadly shown to be involved in monolignol biosynthesis, catalyzing the conversion of cinnamoyl-CoAs into their corresponding cinnamaldehydes (Lacombe et al., 1997; Yeh et al., 2014). Accordingly, the strong upregulation of *FaCCR1(6A)* in the contrasting lines was accompanied by an overall activation of the monolignol biosynthesis pathway, with genes from previous (e.g., the 4-coumarate-CoA ligase *Fxa5Cg201958*, EC 6.2.1.12) and later (e.g., cinnamyl alcohol dehydrogenases *Fxa2Cg200347*, *Fxa2Dg203268* and *Fxa2Ag103710*, EC 1.1.1.195) steps also upregulated (Supplementary Fig. S6F). However, besides this well-known activity of FaCCR1, its reductase activity could potentially target additional acyl-CoAs, providing aldehyde intermediates which would serve as precursors for ester biosynthesis. Altogether, these findings support the selection of *FaCCR1(6A)* and *FaOCT4(6A)* as candidate genes controlling fruit MCE content in strawberry.

### Transient overexpression of *FaCCR1(6A)* and *FaOCT4(6A)* led to MCE accumulation

To determine whether *FaCCR1(6A)* or *FaOCT4(6A)* influences MCE accumulation, we transiently overexpressed both genes in fruits of the cultivar ‘Rociera’. Fruits overexpressing *FaCCR1(6A)* displayed a significant increase in the concentration of 11 MCEs previously associated with *qMCE-6A* QTL, averaging 25-fold higher levels compared with fruits agroinfiltrated with the *35S:pBI-GUS* control construct. Notably, among the VOCs induced by *FaCCR1(6A)* overexpression, key VOCs associated with consumer liking such as butyl acetate, butyl butanoate and hexyl acetate were found (Fig. 4A-B; Supplementary Table S10). As expected, no significant differences were observed in the concentration of short-chain esters such as methyl acetate or methyl butanoate, which were not controlled by *qMCE-6A*.

**Figure 4.**
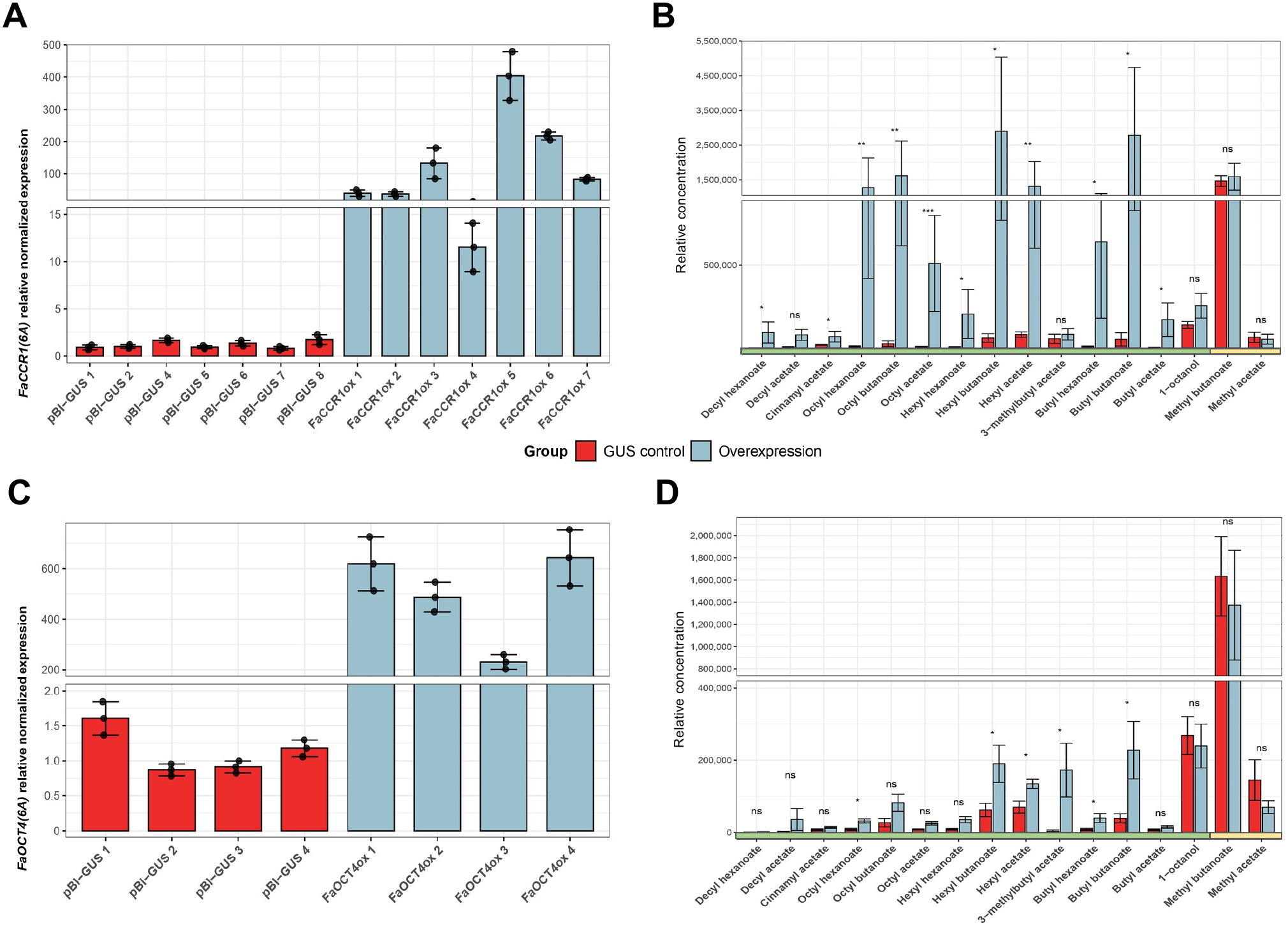
*FaCCR1(6A)* and *FaOCT4(6A)* transient overexpression induce MCE accumulation in strawberry fruit. **A)** Transient overexpression of *FaCCR1(6A)* and **C**) *FaOCT4(6A)* in ripe ‘Rociera’ fruits. Expression was analyzed by RT-qPCR in fruits from seven overexpressing (FaCCR1ox) or mock inoculated (pBI-GUS) lines for CCR1ox. For the OCT4ox experiment, four pools of two fruits per condition were analyzed. Bars represent mean values of three technical replicates and whiskers show SEM. **B)** Ester concentration in fruits overexpressing *FaCCR1(6A)* (blue bars) in comparison to controls (red bars) and **D)** in fruits overexpressing *FaOCT4(6A).* Average values ± SEM from the samples analyzed in each group were plotted. The green horizontal bar indicates compounds controlled by the *qMCE-6A* QTL, whereas the yellow bar encloses short-chain esters. Asterisks show statistically significant differences (* *p*-value ≤ 0.05; ** *p*-value ≤ 0.01; *** *p*-value ≤ 0.001; ns not significant) using Wilcox test for non parametric data or *t*-test for parametric data.

Transient overexpression of *FaOCT4(6A)* also led to a significant increase in six MCEs, whereas the concentration of other short-chain esters remained unchanged (Fig. 4C-D). However, the magnitude of this effect was substantially lower, with an average 2-fold increase in MCE content compared with the 25-fold increase observed following *FaCCR1(6A)* overexpression. Taken together, these results suggest that, although *FaOCT4(6A)* may contribute to MCE variation within the 6A QTL interval, *FaCCR1(6A)* is likely the primary gene controlling natural variation in these important strawberry aroma compounds. These results prompted us to further investigate whether natural variation in *FaCCR1(6A)* expression underlies the observed differences in MCE accumulation in strawberry.

### Identification of cis-regulatory haplotypes associated with *FaCCR1(6A)* expression and MCE content

To further investigate the relationship between *FaCCR1(6A)* expression and MCE accumulation, we quantified transcript levels by RT-qPCR across the entire ‘232’ × ‘1392’ population and observed a strong positive correlation with total MCE content (*r*= 0.70; Supplementary Fig. S8). We then performed an expression quantitative trait *locus* (eQTL) analysis across the same population, using *FaCCR1(6A)* transcript abundance as a quantitative trait, to determine whether the genetic variation underlying the *qMCE-6A* QTL contributes to the regulation of *FaCCR1(6A)* expression. In parallel, QTL analysis was conducted using the mean content across the three seasons for six selected MCEs showing the highest PVE by *qMCE-6A*. A major cis-eQTL controlling 65.2% of the variation in *FaCCR1(6A)* expression colocalized with *qMCE-6A* for the 3 seasons, which explained 43.7-50.6% of the phenotypic variance in the analyzed esters (Supplementary Fig. S9). These results indicate that cis-regulatory variation at the *qMCE-6A* locus regulates *FaCCR1(6A*) expression and MCE content in strawberry fruit.

Genetic variants within regulatory regions can alter transcription factor binding affinities or chromatin topology, thereby leading to allele-specific differences in gene expression. To investigate this possibility, we proceeded to identify the different *FaCCR1(6A)* haplotypes present in the ‘232’ × ‘1392’ population and assessed their association with variation in *FaCCR1(6A)* expression levels. Initially, we included in the analysis the parental lines ‘232’ and ‘1392’ together with eight F_1 l_ines displaying contrasting *FaCCR1(6A)* expression levels (Supplementary Fig. S10). A 1,147 bp fragment corresponding to the proximal upstream regulatory region of *FaCCR1(6A)* (−1086 to +61 relative to the ATG start codon) was amplified using subgenome-specific primers and subjected to direct Oxford Nanopore Technologies (ONT) sequencing.

Three *FaCCR1(6A)* haplotypes, designated *H1-H3*, were identified (Supplementary Fig. S10A). As predicted for the *qMCE-6A* locus, both parental lines were heterozygous at the promoter region. *FaCCR1(6A)-H1* was associated with increased *FaCCR1(6A)* promoter activity, whereas both *H2* and *H3* were associated with low expression levels. Six single-nucleotide variants were identified exclusively in the *FaCCR1(6A)-H1* haplotype (Supplementary Fig. S10B), whose potential contribution to enhanced expression and higher MCE accumulation remains to be established.

To validate the association between the *FaCCR1(6A)-H1* haplotype and enhanced *FaCCR1(6A)* expression across the entire ‘232’ × ‘1392’ population, we developed a KASP assay targeting the *H1*-specific A/G SNP located 543 bp upstream of the start codon (Supplementary Fig. S10). Instead of three expected genotype classes (A:A, A:G and G:G), four different clusters were resolved, corresponding to the *H1*:*H1*, *H1*:*H2*, *H1*:*H3* and *H2*:*H3* haplotype combinations. Marker genotypes were significantly associated with both *FaCCR1(6A)* transcript abundance and MCE accumulation (Supplementary Fig. S10C; Supplementary Table S11). Individuals homozygous for *H1* exhibited the highest expression and MCE levels, whereas *H2:H3* genotypes showed the lowest values, with reductions of up to 80% for both traits. Heterozygous combinations carrying *H1* (*H1:H2* and *H1:H3*) displayed intermediate phenotypes, consistent with a partially dominant effect of the favorable haplotype. Thus, the −543 SNP represents a useful marker for the *H1* haplotype, but further studies are needed to establish whether this SNP or other polymorphisms account for the enhanced *FaCCR1(6A)* expression associated with this haplotype.

### *FaCCR1(6A)* encodes a highly conserved protein localized to the cytosol

Genes encoding CCRs have been identified and characterized in several species and belong to a highly diverse small gene family, whose members are classified as genuine *CCRs* or *CCR-like* genes (Barakat et al., 2011). We compared the amino acid sequence of *FaCCR1(6A)* with reported CCR proteins from other species, selected based on the CCR motifs described previously (Lacombe et al., 1997), along with other CCR-like proteins. A phylogenetic analysis placed *FaCCR1(6A)* within the *bona fide* CCR clade, exhibiting high homology among the Rosaceae family members (>80%), while the CCR-like proteins formed clearly distinct clusters (Supplementary Fig. S11; Supplementary Table S12). A high degree of similarity was observed between FaCCR1 and the two genuine CCRs from *Arabidopsis thaliana*. In this sense, FaCCR1 shared a 77.7% identity with AtCCR1, compared to a 72.6% with AtCCR2. While AtCCR1 and AtCCR2 shared 81.1% identity, FaCCR1(6A) and FaCCR2(5A) (*Fxa5Ag200102*) shared only 63.2%. A multiple alignment of genuine CCR proteins revealed that FaCCR2(5A) presented a 65-aa deletion that included the catalytic domain, explaining the reduced similarity, and suggesting it might not be functional (Supplementary Fig. S12). Furthermore, RNA-seq analysis from different tissues of ‘Camarosa’ (Sánchez-Sevilla et al., 2017) revealed that this gene was not expressed in any tissue (data not shown). No other *FaCCR2* genes were detected on the homoeologous chromosomes 5B, 5C or 5D. The multiple alignment revealed that FaCCR1 shared the CCR signature motif (KNWYCYGK), as well as the S-Y-K triad in the catalytic center, the domains associated with NADPH binding and specificity, and the substrate recognition (Barakat et al., 2011; Chao et al., 2017; Pichon et al., 1998). Moreover, *FaCCR1(6A*) conserves the SSIGAVY motif driving broader substrate specificity, in contrast to the Y to T substitution associated with higher specificity for feruloyl-CoA binding in the *Sorghum bicolor* gene *SbCCR1* (Sattler et al., 2017). These findings indicate that *FaCCR1(6A)* encodes a CCR1 protein, consistent with the previously characterized *FaCCR1* allele from the cultivar ‘Elsanta’ (GenBank accession JX290510; Yeh et al., 2014), with which it shares 99.3% sequence identity.

CCR1 has been reported as a globular, soluble protein localized at the cytosol in species such as *Sorghum bicolor* (Li et al., 2016) and rice (Kawasaki et al., 2005). *In silico* topology prediction using the “DeepTMHMM” tool (https://dtu.biolib.com/DeepTMHMM) also predicted a cytosolic localization of FaCCR1. We determined the subcellular localization of FaCCR1 in *N. benthamiana* leaves using confocal microscopy by coexpressing FaCCR1 fused to the green fluorescent protein (GFP) together with the cytosolic mCherry marker. As expected, FaCCR1-GFP fully colocalized with free *mCherry* at the cortical plane, confirming its predicted cytosolic localization (Fig. 5A). The integrity of the FaCCR1-GFP protein was verified by Western blot (Fig. 5B) before checking its subcellular localization.

**Figure 5.**
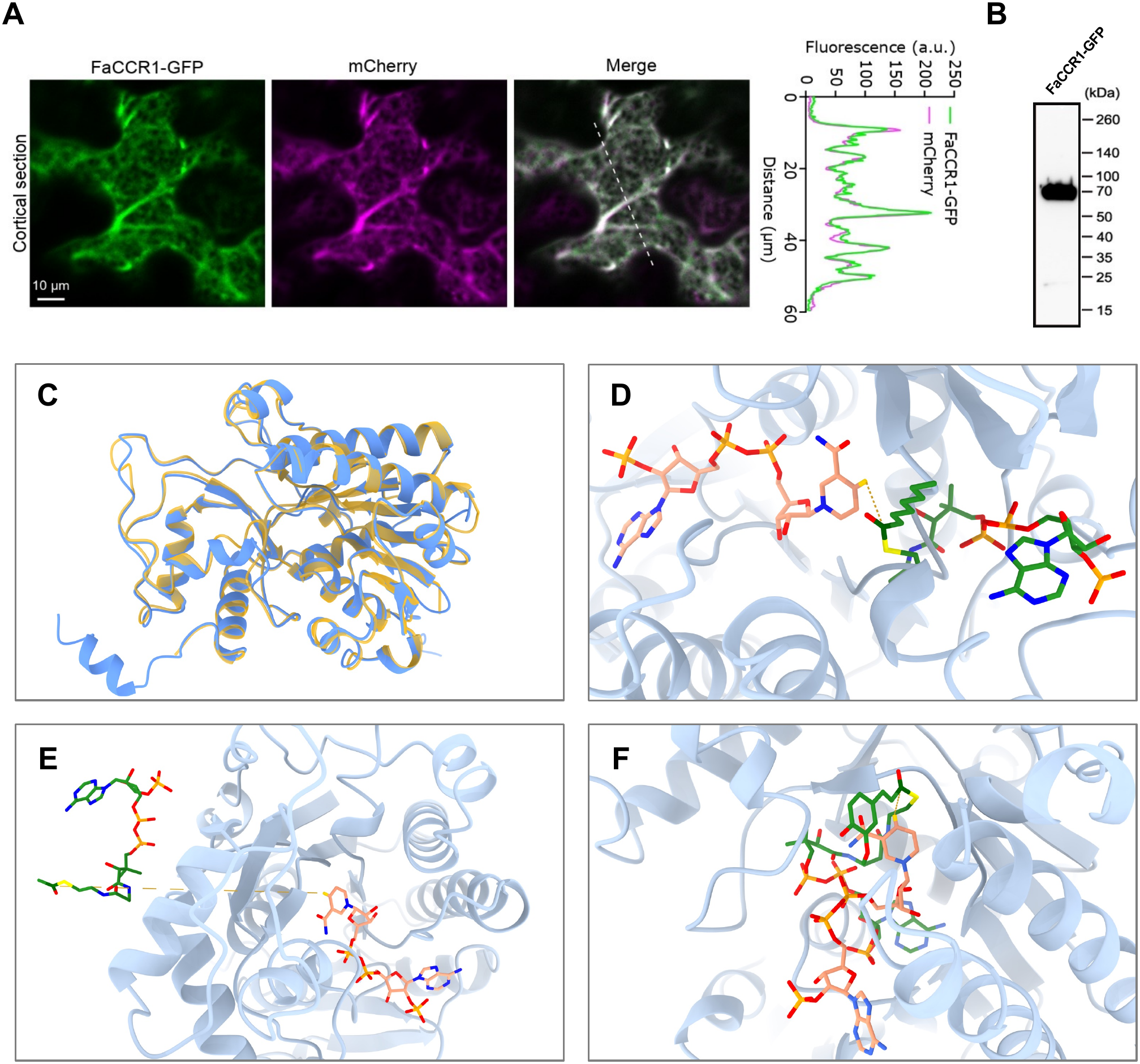
Cytosolic localization and representative poses of FaCCR1(6A) with selected acyl-CoA substrates and structural comparison with SbCCR1. **A)** Coexpression of *a* C-terminal *FaCCR1(6A)-*GFP translational fusion with a cytosolic marker (free mCherry) in *Nicotiana benthamiana* leaves. The cortical region (maximum Z-projection of six slices with 1 μΜ separation between each slice) of abaxial epidermal cells from 2 days post infiltration leaves is shown in individual channels for each protein (two left panels) and merged (right panel). The intensity plot along the white dashed line is shown. Scale bar = 10 μm. Images are representative of all cells analysed (n ≥ 20 cells from 4 randomly chosen regions of interest of infiltrated leaves per experiment). Three independent experiments were performed with consistent results. **B)** Immunoblot analysis of FaCCR1-GFP protein expressed in *N. benthamiana* leaves. Full scan of the immunoblot demonstrate the integrity of FaCCR1-GFP protein. **C)** Global structural superposition of FaCCR1 and SbCCR1 docking models, illustrating the conserved CCR fold used for comparative docking analysis. **D)** Representative FaCCR1 docking pose with octanoyl-CoA, showing a near-attack arrangement between the NADPH nicotinamide H4N atom and the substrate thioester carbonyl carbon. **E)** FaCCR1 docking pose with acetyl-CoA, used as a negative-control substrate, showing lack of productive NADPH–substrate positioning. **F)** Representative FaCCR1 docking pose with feruloyl-CoA, showing NADPH proximity but incomplete catalytic orientation relative to the strict full-geometry criteria. Proteins are shown as cartoons; FaCCR1 is colored light blue and SbCCR1 gold. NADPH is shown in light salmon and acyl-CoA substrates in green, with heteroatoms colored by element. Dashed yellow lines indicate the H4N– *C_carbonyl_* geometric readout used to evaluate NADPH–substrate proximity. Representative poses were selected from the final 500-decoy docking validation panel. These structural docking views illustrate geometric compatibility and should not be interpreted as kinetic measurements.

### Molecular docking analyses reveal substrate-dependent catalytic geometries in FaCCR1

It is well established that CCR1 catalyzes the NADPH-dependent reduction of hydroxycinnamoyl-CoA thioesters to the corresponding cinnamaldehydes in the monolignol biosynthesis pathway (Muhlemann et al., 2014). In strawberry, FaCCR1 has been reported to catalyze the reduction of caffeoyl-CoA, p-coumaroyl-CoA and, with higher preference, feruloyl-CoA (Yeh et al., 2014). Given the association between *FaCCR1(6A)* expression and MCE content, we asked whether FaCCR1 could also accommodate aliphatic medium-chain acyl-CoAs in a NADPH-compatible catalytic geometry. These substrates could generate medium-chain aldehydes that could be further reduced to alcohols and incorporated into MCE biosynthesis by alcohol acyltransferases (Aharoni et al., 2000; Beekwilder et al., 2004; Dudareva et al., 2013; Schwab et al., 2008).

To examine this possibility, we performed a Rosetta-based docking validation panel using FaCCR1 and the structurally characterized sorghum SbCCR1 as a comparator (Sattler et al., 2017). The FaCCR1 AlphaFold3 model and SbCCR1 structure displayed a conserved global fold after superposition, supporting comparison of their NADPH-containing catalytic pockets (Fig. 5C). Rather than relying only on docking score, we evaluated 500 independent decoys per enzyme-ligand condition using NADPH-substrate catalytic-geometry metrics. This analysis distinguished poses with simple cofactor proximity from those satisfying the complete distance and angular criteria expected for hydride transfer (Table 2; Supplementary Tables S13 and S14).

**Table 2.**
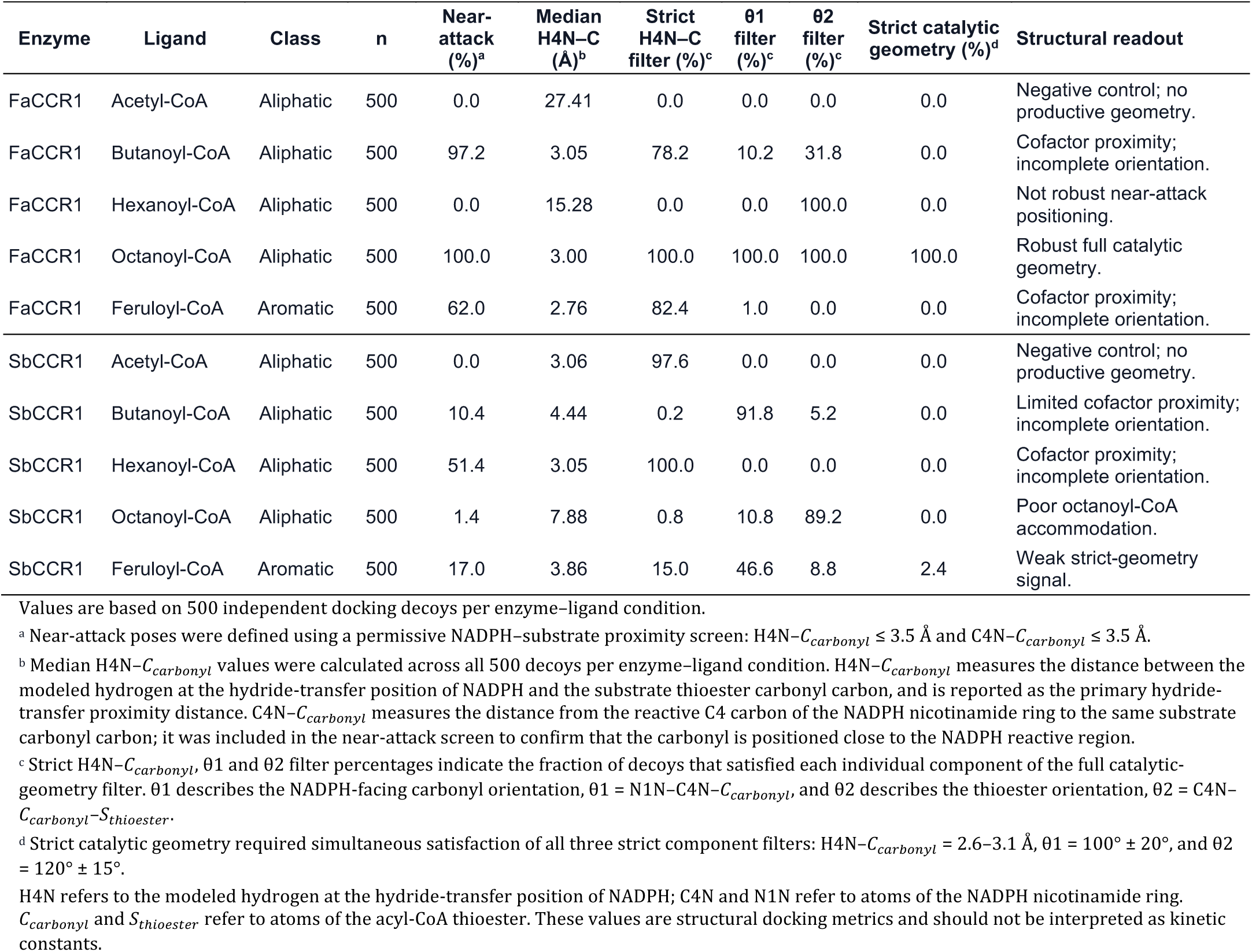
Catalytic-geometry analysis of FaCCR1 and SbCCR1 docking ensembles.

| **Enzyme** | **Ligand** | **Class** | **n** | **Near-attack (%)<sup>a</sup>** | **Median H4N–C (Å)<sup>b</sup>** | **Strict H4N–C filter (%)<sup>c</sup>** | **θ1 filter (%)<sup>c</sup>** | **θ2 filter (%)<sup>c</sup>** | **Strict catalytic geometry (%)<sup>d</sup>** | **Structural readout** |
|---|---|---|---|---|---|---|---|---|---|---|
| FaCCR1 | Acetyl-CoA | Aliphatic | 500 | 0.0 | 27.41 | 0.0 | 0.0 | 0.0 | 0.0 | Negative control; no productive geometry. |
| FaCCR1 | Butanoyl-CoA | Aliphatic | 500 | 97.2 | 3.05 | 78.2 | 10.2 | 31.8 | 0.0 | Cofactor proximity; incomplete orientation. |
| FaCCR1 | Hexanoyl-CoA | Aliphatic | 500 | 0.0 | 15.28 | 0.0 | 0.0 | 100.0 | 0.0 | Not robust near-attack positioning. |
| FaCCR1 | Octanoyl-CoA | Aliphatic | 500 | 100.0 | 3.00 | 100.0 | 100.0 | 100.0 | 100.0 | Robust full catalytic geometry. |
| FaCCR1 | Feruloyl-CoA | Aromatic | 500 | 62.0 | 2.76 | 82.4 | 1.0 | 0.0 | 0.0 | Cofactor proximity; incomplete orientation. |
| SbCCR1 | Acetyl-CoA | Aliphatic | 500 | 0.0 | 3.06 | 97.6 | 0.0 | 0.0 | 0.0 | Negative control; no productive geometry. |
| SbCCR1 | Butanoyl-CoA | Aliphatic | 500 | 10.4 | 4.44 | 0.2 | 91.8 | 5.2 | 0.0 | Limited cofactor proximity; incomplete orientation. |
| SbCCR1 | Hexanoyl-CoA | Aliphatic | 500 | 51.4 | 3.05 | 100.0 | 0.0 | 0.0 | 0.0 | Cofactor proximity; incomplete orientation. |
| SbCCR1 | Octanoyl-CoA | Aliphatic | 500 | 1.4 | 7.88 | 0.8 | 10.8 | 89.2 | 0.0 | Poor octanoyl-CoA accommodation. |
| SbCCR1 | Feruloyl-CoA | Aromatic | 500 | 17.0 | 3.86 | 15.0 | 46.6 | 8.8 | 2.4 | Weak strict-geometry signal. |

Values are based on 500 independent docking decoys per enzyme–ligand condition.
<sup>a</sup> Near-attack poses were defined using a permissive NADPH–substrate proximity screen: H4N–$C_{carbonyl}$ ≤ 3.5 Å and C4N–$C_{carbonyl}$ ≤ 3.5 Å.
<sup>b</sup> Median H4N–$C_{carbonyl}$ values were calculated across all 500 decoys per enzyme–ligand condition. H4N–$C_{carbonyl}$ measures the distance between the modeled hydrogen at the hydride-transfer position of NADPH and the substrate thioester carbonyl carbon, and is reported as the primary hydride-transfer proximity distance. C4N–$C_{carbonyl}$ measures the distance from the reactive C4 carbon of the NADPH nicotinamide ring to the same substrate carbonyl carbon; it was included in the near-attack screen to confirm that the carbonyl is positioned close to the NADPH reactive region.
<sup>c</sup> Strict H4N–$C_{carbonyl}$, θ1 and θ2 filter percentages indicate the fraction of decoys that satisfied each individual component of the full catalytic-geometry filter. θ1 describes the NADPH-facing carbonyl orientation, θ1 = N1N–C4N–$C_{carbonyl}$, and θ2 describes the thioester orientation, θ2 = C4N–$C_{carbonyl}$–$S_{thioester}$.
<sup>d</sup> Strict catalytic geometry required simultaneous satisfaction of all three strict component filters: H4N–$C_{carbonyl}$ = 2.6–3.1 Å, θ1 = 100° ± 20°, and θ2 = 120° ± 15°.
H4N refers to the modeled hydrogen at the hydride-transfer position of NADPH; C4N and N1N refer to atoms of the NADPH nicotinamide ring. $C_{carbonyl}$ and $S_{thioester}$ refer to atoms of the acyl-CoA thioester. These values are structural docking metrics and should not be interpreted as kinetic constants.

In FaCCR1, acetyl-CoA behaved as a negative control, with no near-attack positioning or strict catalytic geometry. Butanoyl-CoA and feruloyl-CoA frequently reached NADPH proximity (97.2% and 62.0% near-attack poses, respectively), but failed to satisfy the full catalytic-geometry filter because of incomplete angular orientation. Hexanoyl-CoA did not show robust near-attack positioning in the primary docking panel. In contrast, octanoyl-CoA showed the most coherent NADPH-compatible catalytic geometry in FaCCR1, with 100% near-attack poses, a median H4N-C distance of 3.00 Å and 100% strict catalytic geometry (Table 2; Fig. 5D). The comparison with SbCCR1 further supported enzyme-dependent differences in substrate accommodation. SbCCR1 did not reproduce the robust octanoyl-CoA geometry observed in FaCCR1, showing only 1.4% near-attack poses and cero strict catalytic geometry for this ligand. Conversely, SbCCR1 showed 17% near-attack poses and a 2.4% strict-geometry signal with feruloyl-CoA, consistent with its known CCR activity toward aromatic acyl-CoAs (Sattler et al., 2017). Overall, the docking analysis supports the structural plausibility that FaCCR1 can accommodate selected aliphatic, medium-chain acyl-CoAs in the NADPH-containing catalytic pocket, while also indicating that cofactor proximity alone is not sufficient to infer a productive catalytic orientation. These structural docking metrics were therefore interpreted as complementary structural support for further biochemical assays, rather than as direct estimates of catalytic efficiency.

### FaCCR1 exhibits activity towards medium-chain ester precursors

To functionally validate the ability of FaCCR1 to produce medium-chain aldehydes, we performed *in vitro* enzymatic assays using purified recombinant protein and aliphatic medium-chain acyl-CoAs as substrates. To obtain a highly pure and functional enzyme, FaCCR1–GFP was affinity-purified from *N. benthamiana* leaves agroinfiltrated with *35S:FaCCR1(6A)-GFP*. Free GFP, purified from leaves transformed with 35S:GFP, served as negative control. Feruloyl-CoA and acetyl-CoA were included as positive and negative substrate controls, respectively, as acetyl-CoA has previously been shown not to be a substrate for CCR enzymes (Wengenmayer et al., 1976). In addition, butanoyl-CoA, hexanoyl-CoA, and octanoyl-CoA were tested as potential novel CCR1 substrates, given their plausible role as MCE intermediates and the structural compatibility suggested by the docking analysis. Initially, the canonical and novel substrates were tested at 100 μM, which are near-saturating conditions (Yeh et al., 2014). As expected, feruloyl-CoA was converted to coniferaldehyde (4.15 ± 0.31 µg/mL; Fig. 6B), whereas no product formation was detected with acetyl-CoA (Fig. 6C). Interestingly, FaCCR1-GFP showed activity for all three novel substrates producing butanal, hexanal and octanal (Fig. 6C; Supplementary Fig. S13). Free GFP, used as a blank control in place of FaCCR1–GFP, did not yield any aldehyde products with any of the substrates tested. These results together with the transient overexpression of *FaCCR1(6A)* in strawberry fruits, underscore a novel substrate activity for FaCCR1 and uncovered a route for medium-chain aldehyde biosynthesis.

**Figure 6.**
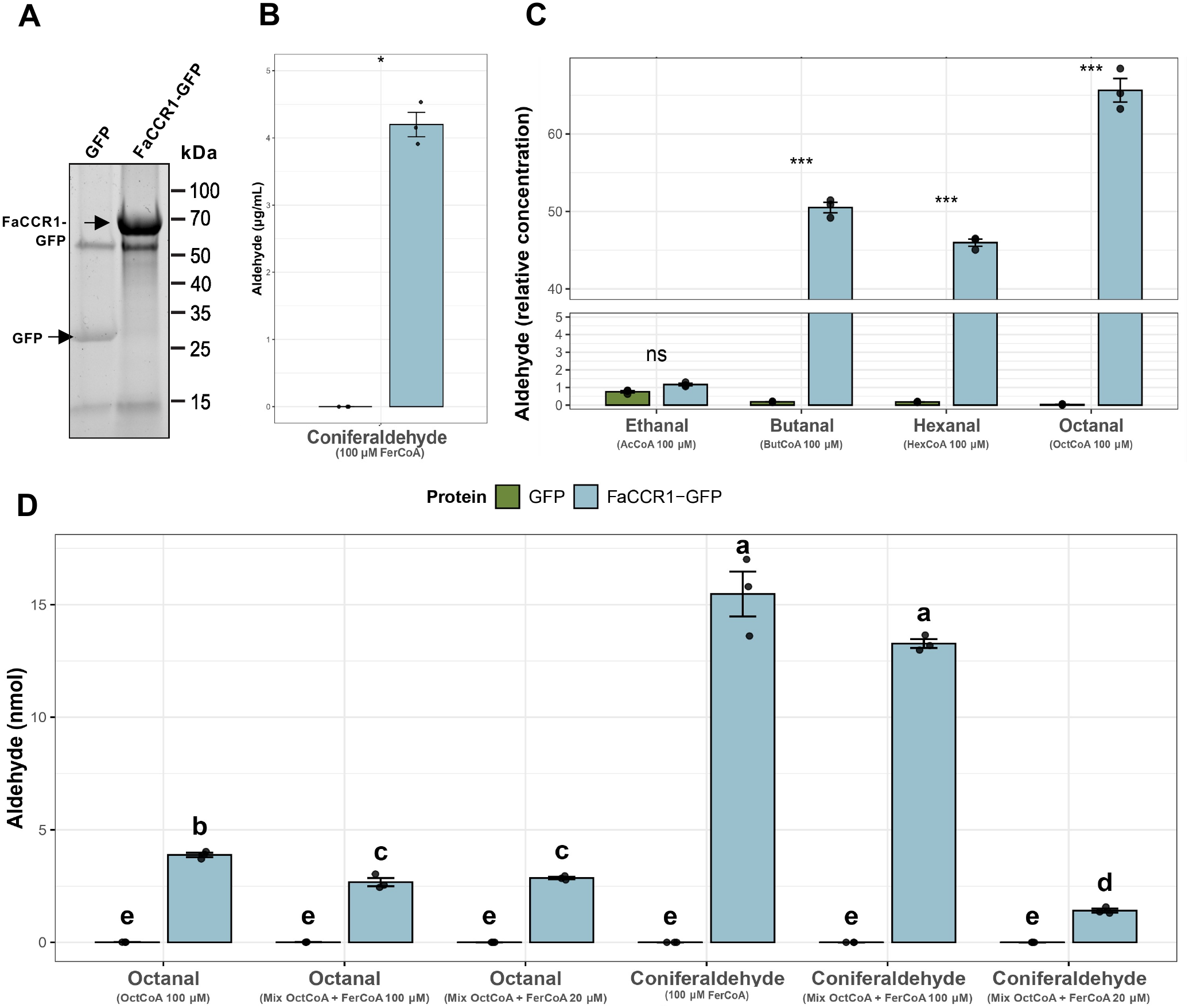
FaCCR1(6A) exhibits reductase activity towards aliphatic medium-chain acyl-CoA esters. **A)** SDS–PAGE for protein quantification and integrity assessment of immunoprecipitated free GFP and FaCCR1–GFP. **B)** Quantification of aldehyde produced in the reactions performed with FaCCR1-GFP protein (blue color) or free GFP as a negative control (green) using feruloyl-CoA as substrate (positive control; Yeh *et al.,* 2014). **C)** Relative concentration of aldehydes produced in enzymatic reactions using acetyl-CoA, negative control (Wengenmayer et al. 1979) or the test substrates (butanoyl-CoA, hexanoyl-CoA and octanoyl-CoA). Asterisks denote statistically significant differences (* *p*-value ≤ 0.01; ** *p*-value ≤ 0.001; *** *p*-value ≤ 0.0001; ns not significant) determined by t-test. **D)** Absolute quantification of aldehyde production in independent or substrate competition assays. Letters denote statistically significant differences *(p*-value < 0.01) between groups, determined by Welch’s one-way ANOVA with *post-hoc* Games-Howell test for multiple comparisons. Bars indicate the aldehyde mean values of three technical replicates and whiskers show SEM.

To directly compare the utilization of medium-chain acyl-CoAs with that of the canonical CCR substrate, a second set of enzymatic assays was performed using commercial aldehyde standards for absolute product quantification. When FaCCR1 was incubated with an equimolar mixture (100 μM each) of butanoyl-CoA, hexanoyl-CoA, and octanoyl-CoA, all three substrates were converted to their corresponding aldehydes (Supplementary Table S15). These results demonstrate that FaCCR1 can simultaneously utilize multiple medium-chain acyl-CoA substrates, although product formation decreased with increasing acyl-chain length. We next evaluated whether medium-chain acyl-CoAs could be utilized in the presence of feruloyl-CoA, the canonical CCR substrate (Fig. 6D). When supplied individually at 100 μM, octanoyl-CoA and feruloyl-CoA were reduced by FaCCR1-GFP to octanal and coniferaldehyde, respectively. Notably, simultaneous incubation with equimolar concentrations of both substrates (100 μM each) resulted in the production of both aldehydes, with octanal and coniferaldehyde levels decreasing by 31.13% and 14.25%, respectively, compared with the corresponding reactions in which each substrate was supplied individually. These results indicate that octanoyl-CoA can be efficiently utilized by FaCCR1-GFP even in the presence of feruloyl-CoA. To further challenge substrate competition, the assay was repeated using a 5-fold lower concentration of both octanoyl-CoA and feruloyl-CoA (20 μM each). Interestingly, under these conditions, octanal production remained unchanged relative to the corresponding mixed-substrate reaction at 100 μM, whereas coniferaldehyde production decreased by 90% (Fig. 6D). Collectively, these results demonstrate that FaCCR1(6A) can catalyze the reduction of both aromatic and aliphatic acyl-CoA substrates, expanding its recognized substrate range beyond aromatic substrates associated with monolignol biosynthesis and supporting a role in medium-chain ester biosynthesis in strawberry fruit.

### Development of KASP assays for the selection of genotypes with high MCE content

The *FaCCR1(6A)-H1* KASP marker developed to genotype the ‘232’ × ‘1392’ population, revealed poor clustering performance in the diverse GWAS panel (*n* = 124), with seven to eight partially overlapping clusters that precluded unambiguous genotype calling. Subsequently, a query using the CENSOR database of repetitive DNA elements showed that *H1*-specific SNPs are located within a region sharing 88% similarity with a hAT DNA transposon, likely compromising primer specificity and preventing reliable amplification. Therefore, we tried to design another assay targeting the c.462C/T SNP identified in the *FaCCR1(6A)* CDS but it didn’t render a suitable KASP/HRM marker either.

As an alternative approach, we developed two KASP assays targeting other polymorphisms within the *qMCE-6A locus*. The first assay was directed against the GWAS lead SNP AX-184056927 (Chr_6A:19785859; Supplementary Table S6), located 0.6 Mb downstream of *FaCCR1(6A)*. However, when tested in the ‘232’ × ‘1392’ population, this marker failed to discriminate between the favorable *H1* and the unfavorable *H3* haplotypes, both of which carried the A allele at this position, whereas the also unfavorable *H2* carried the G allele (Supplementary Fig. S10B). Although G:G genotypes showed a significant decrease in MCE concentration in both populations (Supplementary Fig. S14; Supplementary Table S16), this marker could not accurately track the functional haplotypes underlying the *qMCE-6A* effect.

The second KASP assay targeted the c.254G/A SNP in *FaOCT4(6A)* (chr_6A: 18,799,453), approximately 0.37 Mb upstream of *FaCCR1(6A)* (Fig. 7A). In the ‘232’ × ‘1392’ population, both parental lines were heterozygous for this SNP and the marker segregated in the progeny according to the expected 1:2:1 Mendelian ratio. Consistent with *FaCCR1(6A)* −543 A/G marker, the *FaOCT4(6A)* assay efficiently discriminated the favorable *H1* haplotype (G allele) from the *H2* and *H3* haplotypes (A allele) (Supplementary Fig. S10B) and was significantly associated with MCE accumulation (Fig. 7). Variance components extracted from mixed linear models estimated that this marker accounted for 44% of the phenotypic variance, consistent with the PVE observed for *qMCE-6A*. This effect was reflected in a five-fold reduction in the mean total MCE content in F_1 l_ines homozygous for the unfavorable A allele compared with G:G homozygotes (Fig. 7C; Supplementary Table S17). Given the strong linkage between *FaOCT4(6A)* and *FaCCR1(6A), FaOCT4(6A)* genotype classes also showed significant differences in *FaCCR1* expression levels (Fig. 7D).

**Figure 7.**
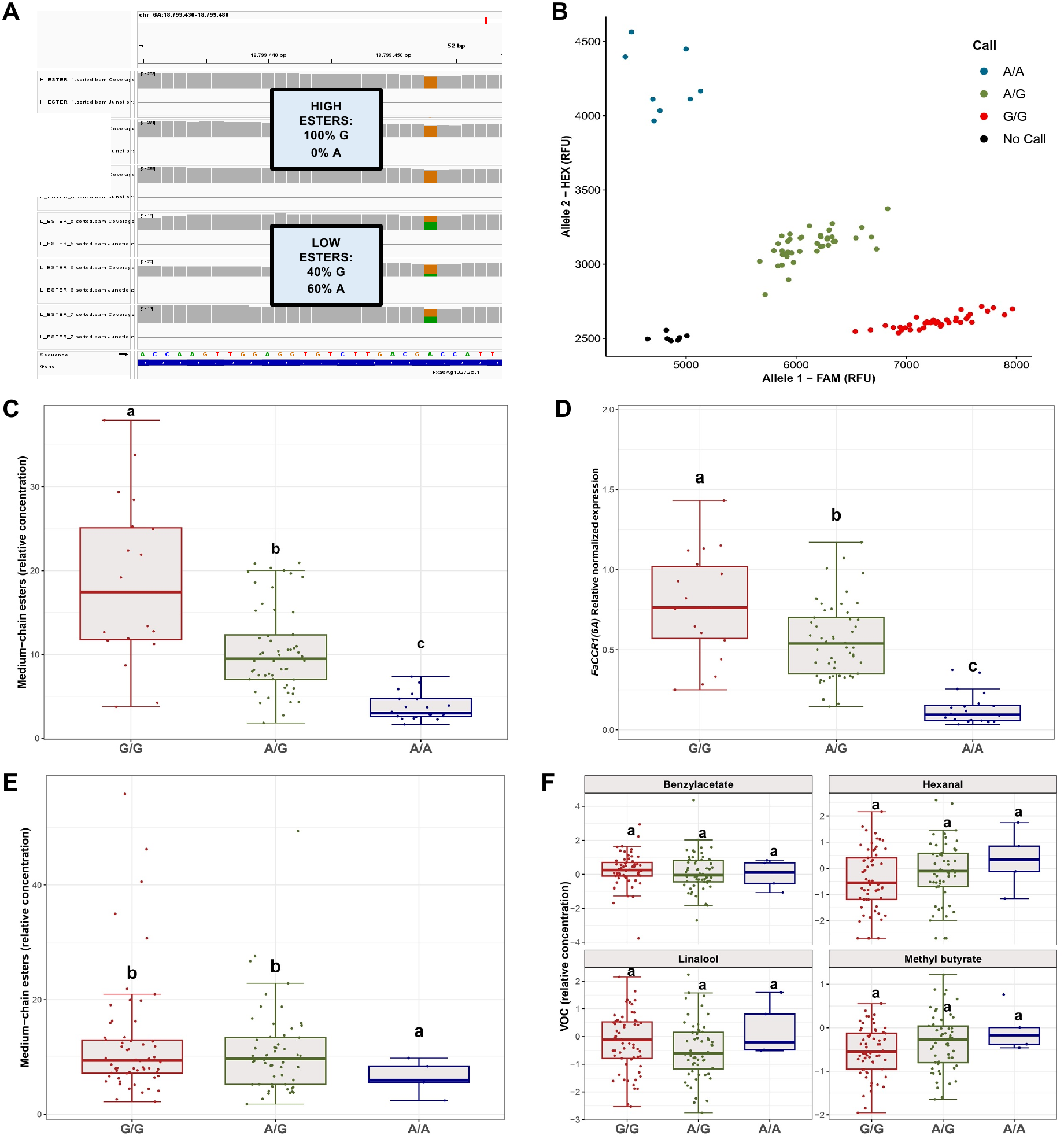
Development and evaluation of a Kompetitive Allele Specific PCR (KASP) assay in *FaOCT4(6A)* for fruit MCE prediction. **A)** IGV visualization of RNA-Seq reads from F_1_ fruit pools of high and low MCE content. The SNP at chr_6A: 18,799,453 position within *FaOCT4(6A)* CDS shows contrasting allelic frequencies. **B)** Example of genotype clusters for KASP *FaOCT4(6A)* marker in the 124 diverse accessions of the IFAPA GWAS population. The reference A allele is labelled with HEX (blue color) and the alternative G allele is labelled with FAM (red color). Heterozygous samples are represented in green and non-template controls or no calls in black. RFU, relative fluorescence units for HEX and FAM fluorescent dyes. **C)** Box-plot of allelic effects of *FaOCT4(6A)* KASP marker on total MCE content in the ‘232’ × ‘1392’ population (n = 96). **D)** Allelic effects of *FaOCT4(6A)* KASP marker on *FaCCR1(6A)* expression. **E)** Box-plots of allelic effects on total MCE content in the 124 diverse accessions from the GWAS collection evaluated in 2020-21, and **F)** in other VOCs not controlled by *qMCE-6A*. Boxes span the 25th and 75th percentiles and the middle line represents the median. Whiskers extend to the minimum and maximum data points within 1.5 times the interquartile range (IQR). Letters denote statistically significant differences *(p*-value < 0.01) between groups, determined by Welch’s one-way ANOVA with *post-hoc* Games-Howell test for multiple comparisons.

Beyond the biparental population, in the GWAS panel (*n* = 124), accessions homozygous for the unfavorable A allele had approximately half the fruit MCE content. (*p*-value < 0.01), while no significant effects were observed for other VOC classes, including short-chain esters, aldehydes, terpenoids, and alcohols (Fig. 7E-F). In this panel, there was a skewed distribution toward the favorable G allele. The unfavorable A allele had a minor allele frequency (MAF) of 0.27, with only 5% of accessions being A homozygotes and 45% A heterozygotes. A Bayesian linear mixed model accounting for the low frequency of A genotypes estimated that the marker was associated with a 24% of the phenotypic variation (Gelman & Pardoe, 2006). Together, these results support *FaOCT4(6A)* KASP marker as a promising tool for selecting high-MCE, highly aromatic strawberry cultivars.

## Discussion

Strawberry aroma is determined by a complex mixture of esters, aldehydes, alcohols, terpenoids, furans, and lactones which have been extensively studied (Jetti et al., 2007; Schieberle & Hofmann, 1997; Ulrich et al., 1997). Esters are qualitatively and quantitatively the most important class of VOCs in strawberry ripe fruit, playing a key role as a source of fruity and floral notes (Jetti et al., 2007; Song et al., 2008). Consequently, there is a great interest in identifying their genetic basis with the aim of improving fruit aroma to meet consumer preferences. Although the optimal ester concentration in fruit remains uncertain, increasing the total content, particularly butanoic and hexanoic derivatives, is generally considered a promising strategy to improve fruit aroma (Porter et al., 2023).

In this work, we narrowed the QTL interval for MCE content on chromosome 6A to a region of about 2 Mb (Fig. 2), previously mapped with lower resolution (Fan et al., 2022; Rey-Serra et al., 2022; Zorrilla-Fontanesi et al., 2012). The GWAS collection exhibits substantial genetic diversity and separation in distinct subpopulations, characterized by low stratification (Muñoz et al., 2024), thereby reducing the detection of false positives in GWAS (Burghardt et al., 2017). LD decay analyses further underscored the high-resolution of the association study enabling fine QTL definition (Ibrahim et al., 2020; Supplementary Fig. S3). Considering this genetic diversity, no phenotypic differentiation was observed based on the geographic or temporal origin, nor to genetic background (Supplementary Fig. S2), which suggests that fruit MCE content has not been affected by a genetic bottleneck resulting from past breeding efforts (Feldmann et al., 2024; Hardigan, et al., 2021b). In agreement, the same QTL was detected in the modern and advanced Floridian collection or in independent biparental populations (Fan et al., 2022; Rey-Serra et al., 2022). Phenotypic analysis in our populations revealed a strong correlation among MCEs, and further association studies identified a region in chromosome 6A accounting for about 35% of the phenotypic variation in both the biparental and the GWAS populations (Fig. 1-2; Supplementary Fig. S4, S9), indicative of a major QTL governing MCE accumulation, regardless of the population or the environment.

High MCE lines exhibited a significant enrichment of the oxidative degradation of fatty acids in the peroxisomes, which lead to the production of key intermediates in ester biosynthesis such as acyl-CoAs and in theory to volatile aldehydes and alcohols (Fig. 3B; Supplementary Fig. S6A; Beekwilder et al., 2004; Schwab et al., 2008). Aldehydes are initially reduced to their corresponding alcohols by alcohol dehydrogenases and subsequently esterified with acyl-CoAs by alcohol acyltransferases (Aharoni et al., 2000; Beekwilder et al., 2004). Alternatively, ‘fresh green aroma’ aldehydes and esters are also produced from fatty acids through the lipoxygenase pathway, which leads to C_6 a_nd C_9 d_erived VOCs as hexanal, hexanol or hexenyl and hexyl acetates (Beekwilder et al., 2004; Dudareva et al., 2013; Schwab et al., 2008). Despite the enrichment of the linoleic acid metabolism in high MCE lines, genes producing green aroma VOCs, such as lipoxygenases, were not upregulated and not likely involved in providing precursors for the MCEs controlled by the *qMCE-6A* QTL (Supplementary Fig. S6E). No differential expression was detected in genes encoding medium-chain acyl-CoA synthetases/ligases (Supplementary Fig. S6C), suggesting that the main source of acyl-CoA is the β-oxidation of fatty acids. In each cycle of β-oxidation, one acetyl-CoA is produced, and the fatty acid is shortened by two carbons (Goepfert & Poirier, 2007), with the resulting –2C acyl-CoA either returning to the β-oxidation cycle or potentially serving as substrate for ester biosynthesis. As one molecule of CoA is consumed in each oxidation cycle, CoA mobilization could contribute to fatty acid metabolism (Supplementary Fig. S6A-B). A notable downregulation of genes from the terpenoid biosynthesis pathway was observed (Supplementary Fig. S6D), which is particularly relevant since cytosolic acetyl-CoA serves as the primary precursor for terpenoid anabolism (Schwab et al., 2008). Thus, the cytosolic acetyl-CoA, and other acyl-CoAs derived from the β-oxidation, such as butanoyl-, hexanoyl- and octanoyl-CoA, may instead be redirected to ester biosynthesis in high MCE lines (Aharoni et al., 2000).

Amino acid degradation has been linked to ester biosynthesis through the generation of aldehydes, alcohols, and acyl-CoAs (Dudareva et al., 2013; Schwab et al., 2008). Accordingly, high MCE lines may exhibit greater precursor availability through the enrichment of amino acid metabolism pathways, including those of cysteine, methionine, serine or phenylalanine (Fig. 3B). In banana, apple and strawberry, branched chain esters, such as methyl-butyl acetate and butanoate, arise from the branched chain amino acids leucine, isoleucine and valine (Dudareva et al., 2013; Schwab et al., 2008). Additionally, alanine has been suggested to be a precursor for ethyl esters in strawberry (Pérez et al., 1992). However, none of these esters were controlled by *qMCE-6A* and, in agreement, no obvious differences in the expression of genes related to these pathways were observed in the contrasting lines. Overall, although differential expressions were often modest, the collective up- or downregulation of gene sets within these pathways may explain the MCE differences in strawberry fruit. Therefore, part of the unexplained phenotypic variance could be attributed to additional QTL of smaller effect controlled by some of these genes.

Multiple lines of evidence support *FaCCR1(6A)* as the primary gene underlying natural variation in MCE accumulation in strawberry. 1) *FaCCR1(6A)* displayed the highest upregulation in the high MCE pool and colocalized within the *qMCE-6A* QTL; 2) its expression increases during receptacle ripening, coinciding with active ester biosynthesis (Beekwilder et al., 2004; Fait et al., 2008; Song et al., 2008); 3) *FaCCR1(6A)* was the most highly expressed *FaCCR1* homoeolog in receptacle tissue, whereas the corresponding homoeologs from the other subgenomes exhibited minimal to negligible expression, and 4) *FaCCR1(6A)* expression showed a strong positive correlation with MCE content in the ‘232’ × ‘1392’ population, consistent with a similar correlation previously reported in a Floridian germplasm collection (Fan et al., 2022). Furthermore, the identification of a cis-eQTL colocalizing with *qMCE-6A* strongly suggests that regulatory variation affecting *FaCCR1(6A)* expression underlies the observed differences in MCE accumulation. In line with this, promoter haplotype analysis revealed allelic variation associated with contrasting expression levels, providing a potential genetic basis for this regulatory effect.

Although a positive correlation between *FaCCR1* expression and lignin content has been reported (Yeh et al., 2014), its weak expression in leaf, stem, runner, and root tissues (Supplementary Fig. S7) suggests a specialized role of the gene in fruits. Given that lignin is present at much higher levels in these vegetative tissues than in the receptacle, the expression pattern of *FaCCR1(6A)* is not consistent with a primary role in lignin biosynthesis and instead points to a fruit-specific function. Most importantly, transient overexpression of *FaCCR1(6A)* resulted in a substantial increase in MCE content (Fig. 4A-B), including several key aroma esters such as butyl acetate, butanoate and hexanoate, hexyl acetate, and octyl butanoate and hexanoate, which are known contributors to strawberry flavor quality (Fan et al., 2021a; Fan et al., 2021b; Porter et al., 2023; Schwieterman et al., 2014). In contrast, although *FaOCT4(6A)* was also differentially expressed between pools with contrasting MCE content, its overexpression caused only a modest increase in a subset of MCEs (Fig. 4C-D). This comparatively weak effect suggests that FaOCT4(6A) may play a secondary or accessory role in MCE accumulation, which needs further research.

The high homology between FaCCR1 and the *bona fide* CCR compared to CCR-like proteins, and the conservation of the known CCR motifs in FaCCR1(6A) strongly suggests that *FaCCR1(6A)* encodes a functional CCR protein (Barakat et al., 2011; Chao et al., 2017; Supplementary Fig. S11-12). The phylogenetic distinction between *bona fide* CCR and CCR-like clades is driven by lack of conservation in the CCR motifs, suggesting no CCR activity in *CCR-like* genes, as shown in switchgrass (Escamilla-Treviño et al., 2010). Two *bona fide* CCR genes exhibiting high homology have been identified in many plant species, leading to a lack of criteria to differentiate the two variants (Zhou et al., 2010), although the higher similarity between *FaCCR1(6A)* and *AtCCR1* supported the classification of this gene as *CCR1* (Supplementary Table S12). We identified an additional *CCR, FaCCR2(5A)*, in strawberry (*Fxa5Ag200102*). However, it is not expressed in any of the analyzed ‘Camarosa’ tissues (data not shown) and a large deletion in its catalytic domain indicates that it does not encode a functional protein (Supplementary Fig. S12).

To date, it is established that CCR1, including the strawberry ortholog, converts hydroxycinnamoyl-CoA esters to their corresponding cinnamyl aldehydes in the monolignol biosynthesis pathway (Muhlemann et al., 2014; Yeh et al., 2014), which is consistent with the enhanced metabolic flux towards lignin biosynthesis observed in the RNA-seq data (Supplementary Fig. S6F). However, the identification of *FaCCR1(6A)* as the most likely gene underlying the *qMCE-6A* QTL raised the question of how a canonical cinnamoyl-CoA reductase influences MCE accumulation. Therefore, we investigated whether FaCCR1 could generate aldehyde intermediates from medium-chain aliphatic acyl-CoA substrates, thereby contributing to strawberry aroma biosynthesis. Although the specific biosynthesis of aldehydes in fruit is not well understood, enzymatic reduction of acyl-CoAs formed by β-oxidation has been previously suggested (Schwab et al., 2008). CCR is a multifunctional enzyme able to use different acyl-CoA substrates (Chao et al., 2017), as illustrated by CcCCR1 from cinnamon, which has recently been shown to convert cinnamoyl-CoA into *trans-*cinnamaldehyde with a 14.7-fold higher efficiency compared to the Arabidopsis AtCCR1 (Ye et al., 2024). In this study, we have evaluated the ability of FaCCR1 to reduce aliphatic acyl-CoAs and thereby generate the corresponding aldehydes intermediates for MCE biosynthesis. To this end, we performed immunoprecipitation to specifically purify FaCCR1 using *N. benthamiana* as an efficient system for recombinant protein production, while preserving post-translational modifications (Botella & Botella, 2016). *In vitro* enzymatic assays demonstrated the ability of FaCCR1 to reduce aliphatic medium-chain acyl-CoAs, including butanoyl-CoA, hexanoyl-CoA and octanoyl-CoA, into their corresponding volatile medium-chain aldehydes (Fig. 6). Absolute quantification in competition assays using equimolar mixtures of octanoyl-CoA and feruloyl-CoA demonstrated that FaCCR1 exhibits appreciable activity toward aliphatic substrates even in the presence of an aromatic substrate. Consistent with these biochemical data, the expanded docking analysis provided complementary structural support for the ability of FaCCR1 to accommodate selected medium-chain acyl-CoAs in an NADPH-compatible catalytic arrangement. Among the aliphatic substrates tested, octanoyl-CoA showed the most robust catalytic-geometry signal in FaCCR1. These docking results should be interpreted as evidence of substrate accommodation and reactive-site geometry rather than as direct estimates of catalytic efficiency, since *in vitro* product formation is also influenced by substrate availability, competition, assay conditions and analytical detection. Together with the enzymatic assays, these structural results support an expanded substrate range for FaCCR1 beyond the canonical hydroxycinnamoyl-CoA substrates. Distinct kinetic parameters have been detailed for CCR1 enzymes across various species using the known substrates to date (Chabannes et al., 2001; Escamilla-Treviño et al., 2010; Lauvergeat et al., 2001; Sattler et al., 2017; Yeh et al., 2014), highlighting the convenience of more-detailed comprehension of the enzymatic kinetics of FaCCR1 with the novel substrates. Notably, another strawberry gene, the O-methyl transferase *FaOMT*, has also been shown to play a dual role in both lignin and flavor volatile biosynthesis (Wein et al., 2002). These findings provide a foundation for the in-depth characterization of the contribution of CCR1 proteins to aroma across Rosaceae species, as well as in other plant families.

Altogether, our findings fill a critical gap in the current understanding of MCE biosynthesis and highlight the key role of *FaCCR1(6A)* upregulation in enhancing aroma in strawberry fruit. Based on these results and previous evidence, we propose a model for MCE biosynthesis in which both alcohols and acyl-CoA precursors for MCEs are primarily derived from fatty acid β-oxidation (Fig. 8), with aldehydes such as butanal, hexanal, and octanal being produced through a FaCCR1-catalyzed reduction of acyl-CoAs. *In vitro* enzymatic assays demonstrate that FaCCR1(6A) possesses an unexpected substrate promiscuity that expands its metabolic role beyond monolignol biosynthesis. Identifying the molecular determinants underlying this activity, as well as exploring the potential to engineer CCR enzymes for tailored volatile production in strawberry and other fruit crops, represents an exciting avenue for future research.

**Figure 8.**
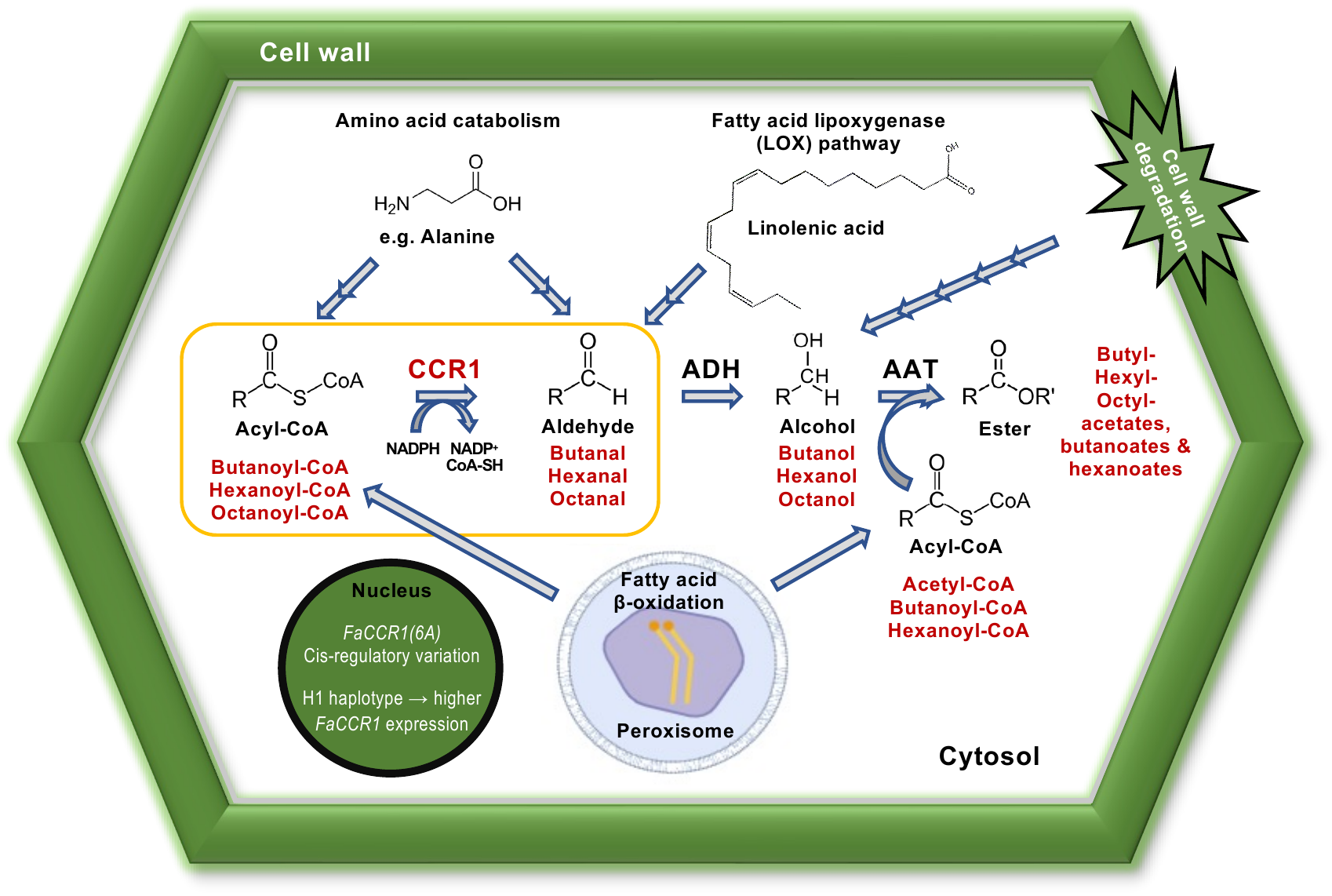
Model for medium-chain ester (MCE) biosynthesis in strawberry. Medium-chain acyl-CoAs, produced from fatty acid β-oxidation (e.g. butanoyl-CoA, hexanoyl-CoA and octanoyl-CoA), are reduced by the cytosolic cinnamoyl-CoA reductase FaCCR1 to their corresponding medium-chain aldehydes (e.g. butanal, hexanal and octanal). Aldehydes can also be generated through the fatty acid lipoxygenase pathway (e.g. C_6_ and C_9_ aldehydes) or amino acid catabolism (e.g. methyl-butanal). Subsequently, alcohol dehydrogenase (ADH) catalyzes their reduction into their corresponding alcohols (e.g. butanol, hexanol and octanol), whose levels could be further reinforced by amino acid catabolism and cell wall degradation. The combination of alcohols with acyl-CoAs, mediated by alcohol acyltransferase (AAT), ultimately leads to the production of MCEs (e.g. butyl acetate, hexyl acetate, octyl acetate). The enzymatic reduction of acyl-CoAs by CCR1, which was inferred in this study, is highlighted in yellow. AAT = alcohol acyltransferase; ADH = alcohol dehydrogenase; CCR1 = cinnamoyl-CoA reductase 1

Despite an estimated LD decay of 800 kb (Supplementary Fig. S3), attempts to narrow down the 6A QTL resulted in a 2-Mb interval, leaving open the possibility that additional genes in the region other than *FaCCR1(6A)* contribute to fruit aroma variation, potentially as part of a gene cluster involved in specialized metabolism, as reported for other plant pathways (Bharadwaj et al., 2021; Fan et al., 2020; Frey et al., 2009; Zhan et al., 2022; Nützmann et al., 2020). In this sense, *FaOCT4(6A)*, whose transient overexpression moderately increased MCE content (Fig. 4C–D), represents one such candidate although its specific role remains unclear. Its Arabidopsis ortholog, the tonoplast-localized AtOCT4, has been implicated in solute homeostasis under drought stress (Küfner & Koch, 2008), whereas plasma membrane-localized AtOCT1 influences root development and carnitine-related processes (Lelandais-Brière et al., 2007). Given that carnitine conjugates with acyl-CoAs during lipid metabolism, as in octanoyl-carnitine formation (Jacques et al., 2018; Nguyen et al., 2016), a role for FaOCT4(6A) in acyl-CoA mobilization is conceivable. Additionally, 13 genes other than *FaCCR1(6A)* and *FaOCT4(6A)* located in the *qMCE-6A* QTL interval were differentially expressed between contrasting MCE fruit pools (Fig. 3C), opening avenues for deeper molecular dissection of the evolutionary dynamics shaping this region, as well as for further clarification of the MCE biosynthetic pathway in strawberry fruit.

To translate our genetic insights into tools for MAS, we developed and validated KASP markers able to predict MCE content across different strawberry populations. We have shown that *FaOCT4(6A)* KASP marker serve as a reliable proxy for the favorable *FaCCR1-H1* haplotype. This marker explained 44% of total phenotypic variance in the biparental ‘232’ × ‘1392’ population, and 24% in the diverse collection, likely due to the influence of additional *qMCE-6A* alleles not targeted by this biallelic marker, the action of linked *loci* within the *qMCE-6A* QTL, and putative additional unidentified QTLs at distant genomic regions. Dominance and potential epistatic interactions may also contribute to the trait, as evidenced by marker behavior: dominant in the GWAS panel but closer to additive in the mapping population, underscoring population-specific genetic architecture (Acquaah, 2012; Hill et al., 2008). Finally, the absence of pleiotropic effects on other VOCs confirms the specificity and utility of this marker for MCE-targeted selection.

## Methods

### Plant material and growth conditions

The strawberry (*F.* × *ananassa*) ‘232’ × ‘1392’ mapping population consists of a full-sib family of 95 F_1 l_ines derived from the cross of two contrasting IFAPA selections: ‘232’, a highly productive line, and ‘1392’, which is characterized by its firmer and tastier fruits (Zorrilla-Fontanesi et al., 2011). This population was previously phenotyped for VOCs during three seasons (2007, 2008 and 2009) and QTLs were identified using a linkage map based on AFLPs and SSRs (Zorrilla-Fontanesi et al., 2012). For this study, six plants of each line were vegetatively propagated and grown under commercial conditions in Huelva, Spain, during the 2012 season. For phenotypic and molecular analysis, 10–15 ripe fruits were collected the same day from the six plants of each line, divided into three biological replicates and independently ground in liquid nitrogen. Samples were stored at −80°C until analyses.

The GWAS panel comprised a diverse set of 124 accessions (Supplementary Table S18) from the *Fragaria* germplasm collection at IFAPA (ESP138). Plants were arranged following a randomized complete block design with 6 replicates per accession and grown in a shaded greenhouse at the IFAPA Centre in Churriana, Málaga (Spain) during season 2020. Further details about growing conditions and the genetic diversity of this population were reported in Muñoz et al., 2024. About 25 ripe fruits from each genotype were collected weekly at midday throughout the peak of the season, immediately frozen in liquid nitrogen, and stored at 80°C. Fruits from each genotype were divided into three replicates, powdered in liquid nitrogen using a coffee grinder and stored at −80°C until analysis. *F.* × *ananassa* ‘Rociera’ plants used for transient gene overexpression in fruit were cultivated in 6.5 L pots in a GMO greenhouse under natural photoperiod and ambient temperatures partially regulated by cooling at the IFAPA Centre in Churriana. *Nicotiana benthamiana* plants used for agroinfiltration were grown in a growth chamber at 25°C with 60% relative humidity under long-day photoperiod (16-hour-light/8-hour-dark cycle).

### QTL and eQTL analysis for medium-chain ester content and *FaCCR1* **expression**

QTL analysis was performed using the ‘232’ξ ‘1392’ integrated map previously developed (Sánchez-Sevilla et al., 2015), relative ester content from Zorrilla-Fontanesi et al., 2012 and the MapQTL 6 software (Van Ooijen, 2009). QTL analysis was also performed using mean values across the three seasons for six selected MCEs and for expression levels of *FaCCR1(6A)* quantified by RT-qPCR. The raw relative data was analyzed first by the nonparametric Kruskal-Wallis rank-sum test, and the significance level of *p* = 0.005 was used as threshold. Second, the integrated genetic linkage map and transformed data sets for most traits were used to identify and locate QTLs using Interval Mapping as previously described in Pott et al., 2020. Significance LOD thresholds were estimated with a 1,000-permutation test for each trait and QTL and LOD scores greater than the genome-wide threshold at 95% were declared significant. Linkage group and QTL confidence intervals in the LG were drawn using MapChart 2.2 (Voorrips, 2002).

### Genome-wide association study (GWAS) for medium-chain esters

Ester content was quantified from the 124 genotypes of the IFAPA GWAS panel using 1 g of homogenized fruit from each of two biological replicates of each accession, each comprising a pool of 7–8 fruits, weighed into separate 5-mL vials. Esters were analyzed by headspace solid-phase microextraction coupled to gas chromatography–mass spectrometry (HS-SPME/GC-MS) following the protocol described by Pott et al., 2021. Compounds were identified by comparison of both mass spectrum and retention time to those of pure standards (SIGMA-Aldrich). Peak areas of selected ions were integrated and normalized against the corresponding peak area in a regularly injected control sample to account for variations in detector sensitivity and fiber aging. Ester relative concentration was expressed on a dry weight basis (DW) of the corresponding fruit sample. DW was calculated gravimetrically by drying fruit samples at 104°C for 3 hours. The average data of the biological replicates for each genotype and ester were Box-Cox transformed when necessary to correct deviations from normality (Box & Cox, 1964) and used for the association study (Supplementary Table S5).

Genomic DNA and genotypic data were previously generated by Muñoz et al., 2024. Briefly, genotypic files from the 50K FanaSNP Axiom^TM^ array (Hardigan et al., 2020) were processed using PLINK v.1.9 software (Chang et al., 2015) to exclude indel and single nucleotide polymorphism (SNP) markers with no position in the ‘Royal Royce’ reference annotation (Hardigan et al., 2021a). After filtering markers with a minor allele frequency (MAF) below 0.05, a total of 40,808 SNPs were preserved. Missing calls were imputed using Beagle v.5.2 with the default settings (Browning et al., 2018, 2021). In order to maximize the resolution of the association analysis, we did not perform LD-pruning of adjacent SNPs prior to GWAS.

GWAS was conducted using both genotypic and phenotypic data of medium-chain ester content. The analysis was performed with the “GAPIT” v.3 package (Wang & Zhang, 2020) in *R* v.4.2.2 (R Core Team, 2020), with the Bayesian-information and Linkage-disequilibrium Iteratively Nested Keyway (BLINK) model (Huang et al., 2019). Manhattan plots were generated using the “fastman” package in *R*. Associations with a −log_10(_*p*-value) exceeding the Bonferroni correction threshold [-log_10(_0.05/40,808)] were accepted as significant. To account for population stratification, the genomic relationship matrix was generated using the VanRaden algorithm (VanRaden, 2008) and included as a random effect, while the top three principal components, explaining the 26.1% of the variance, were used as covariates in the GWAS model. Haplotype blocks were computed using the “SnpStats” package in *R*, where pairwise linkage disequilibrium (LD), measured by *D’*, was calculated with the *ld* function and visualized with the *ld.plot* function from the “gaston” package. LD decay was computed with “PopLDdecay” v3.4.0 package in Linux (Zhang et al., 2019). Population genetics were examined with the *popgen* function from the “snpReady” *R* package.

### RNA-seq, differential expression and enrichment analysis

RNA-seq was performed on high- and low-ester fruit pools from the ‘232’ × ‘1392’ F₁ mapping population, each comprising equal amounts of ripe fruit from 10 F₁ lines selected for contrasting concentrations of six medium-chain esters with the highest PVE by the 6A QTL: butyl acetate, butyl hexanoate, octyl acetate, octyl butanoate, decyl acetate, and octyl hexanoate (Supplementary Table S1). Three biological replicates were analyzed per pool. Total RNA was extracted from 300 mg fruit tissue following the CTAB protocol described in Gambino et al., 2008. Residual genomic DNA was removed using TURBO DNA-free™ (Invitrogen). DNase was subsequently removed by chloroform:isoamyl alcohol extraction, followed by sodium acetate/ethanol precipitation. The RNA pellet was washed with 75% ethanol and resuspended in 25 µL of ultrapure water. RNA integrity was assessed by agarose gel electrophoresis and further verified using a 2100 Bioanalyzer (Agilent, Folsom, CA), which resulted in RIN values ranging between 8.0 and 8.9 for the six samples. RNA sequencing was performed by Illumina’s HiSeq technology to produce 100 pair-end sequences (40 M reads per sample). RNA-seq analysis was carried out essentially as described Vallarino et al., 2019, but using the *F.* × *ananassa* ‘Royal Royce’ genome v1.0 as reference (Hardigan et al., 2021a). Raw RNA-seq reads were filtered to remove low-quality nucleotides and subsequently mapped to the reference genome using the HISAT2 aligner with default settings (Kim et al., 2019) at the Picasso cluster facilities at Servicio de Supercomputación y Bioinformática from Málaga, Spain (http://scbi.uma.es).

Raw count matrix was generated from the resulting .bam files through *FeatureCounts* script from the *R* package “Rsubread” (Liao et al., 2019), including the .gff annotation file from ‘Royal Royce’ genome as reference. Downstream transcriptomic analyses were performed using the “DESeq2” *R* package (Love et al., 2014). The count matrix was transformed into an appropriate *R* object with the *DESeqDataSetFromMatrix* function to establish the comparisons between high and low medium-chain ester groups. Differential expression analysis was then computed through the *DESeq* function and Fragments Per Kilobase Million (FPKM) quantification was performed using the *fpkm* function. Significant differential gene expression threshold was filtered for |log_2(_fold change)| ≥ 1 and adjusted *p*-value < 0.05. Transcripts with a total count sum lower than 10 were removed, as they were considered no expressed genes. Both volcano plot showing RNA-seq differential expression and Principal Component Analysis (PCA) for sample clustering were generated using the “ggplot2” package of *R*.

Gene enrichment analysis was performed with the Kyoto Encyclopedia of Genes and Genomes (KEGG) using the *gseKEGG R* function from the “ClusterProfiler” package (Wu et al., 2021). Gene IDs of ‘Royal Royce’ were transformed into KEGG *Fragaria vesca* database (fve) IDs using nucleotide BLAST against RefSeq sequences from the NCBI annotation release. Bonferroni-Holm correction was applied, with a *p-*adjusted value cutoff > 0.05, a minimum gene size of 10 and 100 of maximum. Enriched pathways were represented using the *dotplot* function and sorted by their normalized enrichment score (Subramanian et al., 2005). Data integration and visualization of enriched pathways of interest were performed using the “pathview” *R* package with the log_2(_fold change) values (Luo & Brouwer, 2013).

### Transient overexpression of *FaCCR1(6A)* and *FaOCT4(6A)* in strawberry fruit

The coding sequences (CDS) of *FaCCR1(6A)* and *FaOCT4(6A)* were amplified from cDNA synthesized using RNA extracted from ripe fruits of the F₁ line 93-22, which accumulates high levels of medium-chain esters, and the High Capacity cDNA Reverse Transcription Kit (Applied Biosystems by ThermoFisher Scientific^TM^). PCR amplification was performed using iProof™ High-Fidelity DNA Polymerase (Bio-Rad), following the manufacturer’s instructions. Primers for cloning (Supplementary Table S19) were designed using Primer-BLAST (https://www.ncbi.nlm.nih.gov/tools/primer-blast/) and included the attB1 and attB2 sequences to enable Gateway™ cloning. The resulting PCR products were cloned into pDONR221 using the Gateway™ BP Clonase™ reaction (Invitrogen) and verified by Sanger sequencing. Overexpression constructs were generated by LR recombination between the entry clones pDONR221-*FaCCR1(6A)* and pDONR221-*FaOCT4(6A)* and the binary vector pK7WG2 (Karimi et al., 2002), which drives transgene expression under the control of the cauliflower mosaic virus 35S promoter (CaMV35S). The resulting constructs were verified by PCR and Sanger sequencing and then transformed into competent *Agrobacterium tumefaciens* AGL0 cells as described by Hofgen & Willmitzer, 1988.

Transient overexpression was performed in *F.* × *ananassa* ‘Rociera’ fruits following the protocol of Hoffmann et al., 2006 with minor modifications. Briefly, a 1:1 suspension mix of *A. tumefaciens* AGL0 culture harboring the *FaCCR1(6A)* or *FaOCT4(6A)* construct in combination with *A. tumefaciens* LBA4404 harboring the p19 suppressor of gene silencing (Voinnet et al., 2003) was coinfiltrated into fruits at the late green/early white stage. As a control, fruits were infiltrated with a 1:1 mixture of Agrobacterium suspensions carrying the pBI-35S:GUS (pBI-GUS) and the p19 constructs. Fruits were harvested after about one week when they were fully ripe, immediately frozen in liquid N_2 a_nd stored at −80°C for further analysis. For *FaCCR1(6A)* overexpression, seven fruits per construct (CCR1ox or pBI-GUS) were analyzed independently as biological replicates. For *FaOCT4(6A)* overexpression, eight fruits from OCT4ox or pBI-GUS were agroinfiltrated and pooled in four biological replicates of two fruits each. Fruit samples were immediately frozen in liquid nitrogen after harvesting, ground to a fine powder and stored at −80°C for further analyses. Transient overexpression was validated by RT-qPCR using gene-specific primers (Supplementary Table S19) as detailed in the next section. Esters were extracted from 0.5 g of cryopreserved ground fruit and quantified by HS-SPME/GC-MS following the protocol described by Rambla & Granell in Rodríguez-Concepción & Welsch, 2020. For identification, a characteristic ion (Q Ion, *m/z*) for each compound was selected. Mean values of ester relative concentrations were compared to those in control samples to explore differences resulting from transient gene overexpression.

### Total RNA extraction and RT-qPCR

Total RNA was extracted from 150 mg of fruit tissue using the Plant/Fungi Total RNA Purification Kit (Norgen Biotek^TM^), followed by removal of residual DNA with the TURBO DNA-free™ Kit (Invitrogen) according to the manufacturer’s instructions. 1 µg of RNA was employed for retrotranscription using the High-Capacity cDNA Reverse Transcription Kit (Applied Biosystems by ThermoFisher Scientific^TM^). qPCR was carried out in a 25 µl reaction volume using a CFX96 thermocycler (Bio-Rad) as previously described (Muñoz-Avila et al., 2022), using the primers listed in Supplementary Table S19. Three technical replicates were performed for each sample. Gene expression levels were calculated according to the ^ΔΔ^Cq method with the CFX maestro software and using *FaDBP* and *FaGAPDH* as reference genes (Galli et al., 2015; Pimentel et al., 2010).

### *FaCCR1(6A)* promoter sequencing

*FaCCR1(6A)* upstream regulatory region was amplified using the subgenome-specific primers ccm211 and ccm210 (Supplementary Table S19) and the Thermo Scientific Phusion high fidelity polymerase (Thermo Scientific™ F530). PCR products were column purified using the FavorPrep™ PCR Clean Up Kit and subjected to ONT sequencing using the premium full-length amplicon pipeline at Plasmidsaurus (Germany).

### Phylogenetic Analysis

BLAST searches were performed against the Genome Database for Rosaceae (GDR; https://www.rosaceae.org/) and Phytozome (https://phytozome-next.jgi.doe.gov/) using the coding sequences of *F. × ananassa FaCCR1(6A)* (Fxa6Ag102766) and Arabidopsis *AtCCR1* (At1g15950) and *AtCCR2* (At1g80820) as queries to retrieve homologous *CCR* sequences from representative *Rosaceae* species and other plant taxa (Supplementary Table S12). The translated sequences were aligned using the Geneious alignment algorithm in Geneious v2021.2.2 (https://www.geneious.com), with a gap-open penalty of 5, a gap-extension penalty of 3, and two refinement iterations. The phylogenetic tree was constructed in the same software using the neighbor-joining method and the Jukes–Cantor genetic distance model. Branch lengths represent the amount of evolutionary change, and the tree was left unrooted.

### Analysis of subcellular localization of *FaCCR1(6A)*

The *35S:FaCCR1(6A)–GFP* vector was generated by LR recombination (Invitrogen™) between the entry clone pDONR221-*FaCCR1(6A)* and the destination vector pGWB5, which enables the expression of FaCCR1(6A) fused to GFP at its C-terminus under the control of the CaMV35S promoter (Nakagawa et al., 2007). The final construct was verified by PCR and Sanger sequencing before being introduced into *A. tumefaciens* strain *GV3101::pMP90* as described by Hofgen & Willmitzer, 1988.

Subcellular localization studies were performed in *N. benthamiana* leaves agroinfiltrated with the *35S*:*FaCCR1(6A)-GFP* and p19 constructs, together with the *UBQ10:mCherry* construct, which expresses mCherry as a red fluorescent cytoplasmic marker following the procedure described in Ruiz-Lopez et al. 2021. Briefly, the abaxial surfaces of the second and third youngest leaves of four-weeks-old plants were infiltrated, and the plants kept in a growth chamber at 23°C for two days after agroinfiltration. Expression of FaCCR1(6A)–GFP was assessed by detecting GFP fluorescence in the infiltrated leaves using a Chemidoc Imaging System (BioRad^TM^) equipped with a StarBright™ Blue 520 fluorescence detector, and by immunoblot analysis with an anti-GFP antibody, which also enabled verification of fusion protein integrity.

Proteins were extracted from 100 mg of sample using 300 μL of 2× Laemmli buffer [125 mM Tris-HCl, pH 6.8, 4% (w/v) SDS, 20% (v/v) glycerol, 2% (v/v) β-mercaptoethanol, and 0.01% (w/v) bromophenol blue] by incubation at 75 °C for 45 min, followed by centrifugation at 20,000 × g for 5 min. The supernatants were subjected to SDS-PAGE, and proteins were transferred to PVDF membranes (Immobilon-P, Millipore) using the Trans-Blot Turbo Transfer System (Bio-Rad), according to the manufacturer’s instructions. GFP-tagged proteins were detected by immunoblotting with a mouse monoclonal anti-GFP antibody (clone B-2, 1:1,000; sc-9996, Santa Cruz Biotechnology) and a peroxidase-conjugated anti-mouse IgG secondary antibody (1:50,000; A9044, Sigma-Aldrich). Chemiluminescent signals were detected using a ChemiDoc XRS+ imaging system (Bio-Rad) with either Clarity Western ECL reagent (Bio-Rad) or SuperSignal West Femto Substrate (Thermo Scientific).

Leaf disks were excised for confocal microscopy. The subcellular localization of the FaCCR1(6A)–GFP fusion protein was examined in optical sections acquired in the cortical plane. Maximum-intensity projections were generated from six optical sections acquired at 1 μm intervals, starting at the cell surface. Images were processed using Fiji software (Schindelin et al., 2012), with contrast adjusted by setting 0.35% of pixels as saturated. GFP signal intensity profile was compared with that of the cytoplasmic mCherry marker to confirm the cytosolic localization of FaCCR1(6A)–GFP.

#### Molecular Docking Analysis

For molecular docking analyses, the FaCCR1(6A) amino-acid sequence was modelled with AlphaFold3 (Abramson et al., 2024) (Abramson et al., 2024). The highest-confidence model was structurally aligned to the *Sorghum bicolor* SbCCR1 crystal structure in complex with NADP+ (PDB 5TQM; Sattler et al., 2017)) using UCSF ChimeraX (Pettersen et al., 2021). The cofactor coordinates from SbCCR1 were used to position NADPH in the FaCCR1 active site, and a hydrogen atom at the C4 position of the nicotinamide ring was defined as H4N to represent the hydride-transfer position. The resulting FaCCR1–NADPH and SbCCR1–NADPH complexes were inspected in ChimeraX, non-relevant molecules were removed, and NADPH was retained as part of the receptor in all docking simulations.

Eight acyl-CoA thioester ligands were evaluated: acetyl-CoA, butanoyl-CoA, hexanoyl-CoA, octanoyl-CoA, decanoyl-CoA, feruloyl-CoA, caffeoyl-CoA and p-coumaroyl-CoA.

Acetyl-CoA was included as a negative-control substrate. Ligand structures were generated from SMILES strings, converted to three-dimensional conformations, energy-minimized and parameterized for Rosetta. Protonation states were assigned for pH 7.4, maintaining deprotonated phosphate groups and the intact thioester group.

Docking simulations were performed with Rosetta3 in enzyme-design mode using RosettaScripts (Fleishman et al., 2011; Richter et al., 2011). FaCCR1–NADPH and SbCCR1–NADPH were used as independent receptor models. For each enzyme–ligand pair, the docking protocol sampled ligand position, orientation and torsional flexibility, followed by repacking and local minimization of active-site side chains. The protein backbone was kept fixed. Soft catalytic-geometry constraints were used to guide sampling toward NADPH-compatible poses, penalizing but not rigidly enforcing deviations from the expected hydride-transfer geometry.

Docking poses were evaluated using structural criteria focused on the relative geometry between NADPH and the substrate thioester carbonyl. These criteria were defined as operational docking filters guided by the NADPH-dependent hydride-transfer mechanism described for CCR enzymes and by general Bürgi–Dunitz-type carbonyl attack geometry (Bürgi et al., 1974; Sattler et al., 2017). Near-attack poses were defined by a permissive proximity screen requiring both H4N–*C_carbonyl_* ≤ 3.5 Å and C4N–*C_carbonyl_* ≤ 3.5 Å. Strict catalytic geometry required simultaneous satisfaction of H4N–*C_carbonyl_* = 2.6–3.1 Å, θ1 = N1N–C4N–*C_carbonyl_* = 100° ± 20°, and θ2 = C4N–*C_carbonyl_*-*S_thioester_* = 120° ± 15°.

The final comparative docking panel comprised 16 enzyme–ligand conditions, corresponding to FaCCR1–NADPH and SbCCR1–NADPH docked independently with the eight acyl-CoA ligands. For each condition, 500 independent docking decoys were generated, yielding 8,000 models overall. The percentage of decoys satisfying each geometric criterion and the median values for H4N–*C_carbonyl_*, θ1 and θ2 were calculated for each enzyme–ligand condition. The main docking readouts are summarized in Table 2, the expanded docking panel is shown in Supplementary Table S15, and the definitions of the geometric criteria and atom selections are provided in Supplementary Table S16. These docking metrics were used as comparative structural readouts and were not interpreted as kinetic constants.

### Immunoprecipitation of FaCCR1-GFP and free GFP for *in vitro* enzymatic assays

FaCCR1-GFP and free GFP immunoprecipitation (IP) was performed based on a reported Co-IP protocol with some modifications (Amorim-Silva et al., 2019). Agroinfiltrated *N. benthamiana* leaves transformed with 35S:FaCCR1(6A)–GFP or the empty pGWB5 (35S:GFP) vector, each co-infiltrated with the p19 construct, were ground to a fine powder in liquid nitrogen. 0.5 g of the powdered tissue was homogenized in 0.5 mL of extraction buffer [50 mM Tris-HCl, pH 7.5; 50 mM NaCl; 10% glycerol; 10 mM EDTA, pH 8; 1 mM NaF; 1 mM Na_2M_oO_4·_2H_2O_; 10 mM DTT; 0.5 mM PefaBlock®; 1% (v/v) P9599 protease inhibitor cocktail (Sigma); 0.2% (v/v) Nonidet P-40, CAS: 9036-19-5 (USB Amersham life science)]. Samples were incubated on an end-over-end shaker for one hour at 4°C to enable protein extraction and then centrifuged at 20,000 x g for 20 min at 4°C. Supernatants were filtered by gravity through Poly-Prep Chromatography Columns (#731-1550 Bio-Rad) in a cold room. An aliquot of each filtrate (input) was retained for subsequent immunoblot analysis.

GFP-Trap® Magnetic Agarose beads (ChromoTek^TM^) were employed to immunoprecipitate FaCCR1-GFP and free GFP through specific binding to the GFP tag. Beads were prewashed four times with extraction buffer lacking Nonidet P-40 detergent, before being added to each sample in equal volume. Then, protein-beads samples were incubated on an end-over-end rocker for 2 h at 4°C, followed by four washes with extraction buffer containing 0.05% Nonidet P-40. Finally, protein-beads were resuspended in 250 µL of 0.1 M sodium phosphate buffer pH 6 [0.013 M Na_2H_PO_4·_7H_2O_; 0.087 M NaH_2P_O_4·_H_2O_ and used in FaCCR1 in vitro enzyme activity assay. An aliquot of 15 μL of protein-beads was used to elute proteins by adding 30 μL of Laemmli extraction buffer 2X and incubating at 75°C for 45 minutes. Eluted proteins were separated by SDS-PAGE in precast Bio-Rad TGX gels (4-20% MP TGX Stain-Free Gel, catalog no. #4568093). Protein quantity and integrity were assessed using the stain-free method (BioRad) following the manufacturer instructions.

### *In vitro* enzymatic activity assay of FaCCR1

FaCCR1 activity was assayed using the immunoprecipitated (IP)FaCCR1-GFP protein following the protocol described in Yeh et al., 2014. Feruloyl-CoA and acetyl-CoA were used as positive control and negative control respectively, while the test substrates included butanoyl-CoA, hexanoyl-CoA and octanoyl-CoA. The reaction mixture contained 1 µg of protein, either IP FaCCR1-GFP or IP free GFP in the blank control, 100 µM of NADPH and 100 µM of acyl-CoA substrate in a total volume of 1 mL, prepared in 0.1 M sodium phosphate buffer pH 6. Samples were gently mixed during 30 min at 25°C and immediately stored at −80°C. Reactions were performed in triplicate. To compare the utilization of aliphatic and aromatic acyl-CoA substrates by FaCCR1(6A), enzyme assays were also performed using combinations of feruloyl-CoA, butanoyl-CoA, hexanoyl-CoA and octanoyl-CoA within the same reaction mixture. Reactions contained equimolar concentrations (either 20 or 100 µM) of the indicated substrates. Product accumulation in mixed-substrate reactions was used to assess the relative ability of FaCCR1(6A) to utilize aromatic and aliphatic acyl-CoA substrates under competitive conditions.

Following incubation, coniferaldehyde formed when feruloyl-CoA was used as the substrate was quantified by UHPLC-MS at the IBMCP Metabolomics Platform (UPV-CSIC, Valencia, Spain) as described by Vazquez-Vilar et al., 2023. 1 µL of the aqueous solution resulting from enzymatic reaction with 1ppm Genistein added as internal standard was directly injected in an Orbitrap Exploris 120 mass spectrometer coupled with a Vanquish UHPLC System (Thermo Fisher Scientific, Waltham, MA, USA). For absolute quantification a standard curve was performed using coniferaldehyde standards.

To detect the products formed when butanoyl-CoA, hexanoyl-CoA, octanoyl-CoA or acyl-CoA were used as substrates, volatile aldehydes butanal, hexanal, octanal or ethanal were quantified by HS-SPME/GC-MS based on the protocol by Rambla & Granell in Rodríguez-Concepción & Welsch, 2020, with a slightly modified sampling procedure: a 500 µL aliquot of the enzymatic reaction was transferred to a 10 mL headspace vial with a screw cap and a silicone/PTFE septum. Then, 500 µL of a 5 M CaCl₂·2H₂O solution was added, followed by gentle mixing. For identification, a characteristic ion (Q Ion, *m/z*) for each compound was selected. The following volatile compounds were unequivocally identified and quantified: acetaldehyde, with a retention time of 4.24 min and a quantifier ion of 29 m/z; butanal, with a retention time of 8.02 min and a quantifier ion of 41 m/z; hexanal, with a retention time of 15.69 min and a quantifier ion of 72 m/z; and octanal, with a retention time of 23.50 min and a quantifier ion of 84 m/z. Peak areas of selected ions were integrated and normalized against the corresponding peak area in a regularly injected control sample-consisting of an equimolar mixture of each compound- to account for variations in detector sensitivity and fiber aging. To compare the capacity of FaCCR1 to reduce each substrate, absolute quantification was conducted using four-pointed calibration curves (correlation coefficients ≥ 0.99) across a working range of 100-10,000 ppb for butanal and hexanal, and 1,000-1,000,000 ppb for octanal.

### KASP assays

Three Kompetitive-allele specific PCR (KASP) markers were developed. The first one, *FaCCR1(6A)-H1*, targeted the A/G SNP located 543 bp upstream of *FaCCR1(6A)* start codon. An additional marker KASP-184056927, was designed targeting the AX-184056927 A/G SNP (chr_6A: 19,785,859 bp in ‘Royal Royce’) from the 50K FanaSNP Axiom^TM^ array (Hardigan et al., 2020), which was significantly associated to medium-chain ester content by GWAS. The third marker, KASP-FaOCT4(6A), was designed targeting a G/A SNP at chr_6A: 18,799,453 bp in the ‘Royal Royce’ genome, located in the CDS of *FaOCT4(6A)* (ATG +254 bp). PolyOligo1.0 in Linux (https://github.com/MirkoLedda/; Ledda et al., 2019 was employed to design KASP primers (Supplementary Table S19) using default settings and the ‘Royal Royce’ reference genome. PCRs were carried out on a CFX96 real-time thermal cycler (Bio-Rad) using the KASP-TF Master Mix (LGC Genomics) in a 10 µl reaction containing 0.2 µM of HEX and FAM primers, 0.5 µM of common primer and 20 ng of gDNA. KASP thermal cycling conditions were as follows: 1) 94°C for 15 min; 2) 94°C for 20 s; 3) 61°C for 1 min (drop −0.5°C per cycle); 4) Repeat steps 2-3 nine times (a total of 10 cycles), achieving a final annealing temperature of 56°C; 5) 94°C for 20 s; 6) 57°C for 1 min; 7) Repeat steps 5-6 27 times (a total of 28 cycles); 8) Read step at 37°C for 1 min. Following completion of the KASP thermal cycling program, when genotype clusters were not clearly resolved, up to three additional recycle steps were performed until satisfactory cluster separation was achieved. Each recycle step comprised three PCR cycles (94°C for 20 s and 57°C for 1 min), followed by plate reading. The *FaCCR1(6A)-H1* KASP marker required modified thermal cycling conditions, consisting of 1) 94°C for 15 min; 2) 94°C for 20 s; 3) 67°C for 1 min (drop −0.6°C per cycle); 4) Repeat steps 2-3 nine times (a total of 10 cycles), achieving a final annealing temperature of 61.6°C; 5) 94°C for 20 s; 6) 61.5°C for 1 min; 7) Repeat steps 5-6 14 times (a total of 15 cycles); 8) 94°C for 20 s; 9) 61°C for 1 min; 10) Repeat steps 8-9 24 times (a total of 25 cycles); 11) Read step at 37°C for 1 min. After completion of this program, up to 4-5 recycling steps were required to achieve satisfactory cluster separation. Each recycling step consisted of three PCR cycles (94 °C for 20 s and 61 °C for 1 min), followed by fluorescence reading. The proportion of phenotypic variation attributed to the *FaOCT4(6A)* KASP marker was calculated by 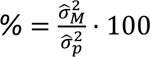, where 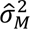 represents the genetic additive variance and 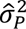 defines the total phenotypic variance, which includes both genetic and residual effects 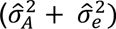 (Falconer & Mackay, 1996; Mathew et al., 2018). Variance components were extracted from the random effects of a linear mixed model, implemented with the “lme4” *R* package. If necessary, a Bayesian equivalent model, constructed with the “blme” *R* package (Chung et al., 2013) was used to deal with model overfitting caused by small sample size (Gelman & Pardoe, 2006). To account for potential bias in variance estimates due to unbalanced genotypic data, a correction factor was applied following the method proposed by (Feldmann et al., 2021).

### Statistical analyses

All statistics were conducted using the *R* programming language. Data normality and homogeneity of variances were examined by the *shapiro.test* and *leveneTest* functions respectively. When data adjusted to a normal distribution, pairwise comparisons were computed using *t*-test. If the data deviated from normality, the Wilcoxon test was used for pairwise comparisons, or data was transformed. Multiple comparison tests were performed using Welch’s one-way ANOVA followed by a *post-hoc* Games-Howell test (*oneway.test* and *games_howell_test* functions, respectively) when the homogeneity of variances assumption was not met. Otherwise, standard ANOVA with *post-hoc* Tukey HSD test were applied (*aov* and *TukeyHSD* functions). Box-Cox transformation was performed following the formula 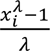 (Box & Cox, 1964), where *x* is the phenotypic data for *i* genotype and *λ* is the scalar calculated with the *boxcox* function from the “MASS” package. Log_10-_transformed data were used to compute Pearson correlations with the “corrplot” package, while PCA was performed and plotted through the “FactoMineR” and “Factoextra” packages. Data and statistics visualization were carried out using “ggplot2” and its extended version, “ggpubr”, R packages. ANOVA and/or *t*-test results for the data presented in each figure are reported in Supplementary data set 1.

### Accession Numbers

The RNA-seq data generated in this study have been deposited in ArrayExpress at EMBL-EBI under accession number **E-MTAB-15704**.

## Acknowledgments

This work was supported by grants PID2022-138290OR-I00 and PID2025-174691OB-I00 funded by MICIU/AEI/ 10.13039/501100011033 and by ERDF/EU. The *Fragaria* collection at IFAPA is financed by IFAPA Project PR.CRF.CRF202200.002 with funds from the European Agricultural Fund for Rural Development. Francisco Javier Roldán-Guerra acknowledges the PhD grant PRE2020-094454 funded by MICIU/AEI/10.13039/501100011033 and by ESF Investing in your future. Vítor Amorim-Silva was funded by a grant (“Programa Emergia 2023”, DGP_EMEC_2023_00375) from the “Consejería de Universidad, Investigación e Innovación de la Junta de Andalucía” (Regional Ministry of Universities, Research, and Innovation of the Regional Government of Andalusia). We are grateful to Francisco J. Durán for his excellent care of strawberry plants and to the IBMCP Plant Metabolomics lab for support in volatile analysis. We also acknowledge José Mora and Sonia Osorio for detection and quantification of fruit esters in the GWAS collection by HS-SPME/GC-MS at the “Unidad de Espectrometría de Masas” at SCAI-Universidad de Málaga. We thank Noemí Ruiz-López for providing the UBQ10:mCherry construct.

## Author Contributions

FJR-G performed most experiments. JRdR participated in transient overexpression in strawberry. VA-S and MAB supervised transient overexpression of proteins in *N. benthamiana*, protein extractions, immunoblots, immunolocalizations, immunoprecipitations and FaCCR1 enzymatic assay. JJ and AG detected and quantified targeted volatiles in agro-infiltrated strawberry fruits and in enzymatic reactions by HS-SPME/GC-MS or UHPLC-MS. AM-A and AG performed molecular docking studies. RT contributed to KASP development and validation. JFS-S maintains the germplasm collection, participated in GWAS and supervised RNA-seq analysis. CC analyzed data and supervised the study. IA conceptualized and supervised the study and secured funding. FJR-G wrote the manuscript with the help of CC and IA. All authors contributed to revising the manuscript.

## Supplementary data

The following materials are available in the online version of this article.

**Supplementary Figure S1**. PCA biplot in the GWAS population.

**Supplementary Figure S2**. Total MCE content in the six subpopulations from the diverse collection.

**Supplementary Figure S3**. LD decay in the diverse collection.

**Supplementary Figure S4**. Manhattan plots showing significant marker-trait associations from GWAS for MCEs.

**Supplementary Figure S5**. MCE content in the fruit pools used for RNA-Seq.

**Supplementary Figure S6**. KEGG pathview graphs enriched in the high MCE lines.

**Supplementary Figure S7**. DEGs within the qMCE-6A QTL.

**Supplementary Figure S8**. Expression of *FaCCR1* in the ‘232’ × ‘1392’ population.

**Supplementary Figure S9**. eQTL analysis.

**Supplementary Figure S10**. Alleles of the 5’ upstream regulatory region of *FaCCR1(6A)*

**Supplementary Figure S11**. Neighbour-joining phylogenetic tree of CCR and CCR-like amino acid sequences.

**Supplementary Figure S12**. Multiple alignment of cinnamoyl-CoA reductase amino acid sequences.

**Supplementary Figure S13**. Chromatograms of aldehydes resulting from enzymatic reactions with GFP or FaCCR1-GFP.

**Supplementary Figure S14.** KASP marker assay in SNP AX-184056927 for fruit MCE prediction

**Supplementary Table S1**. Relative concentrations of medium-chain esters and alcohols controlled by QTL on 6A in the ‘232’ × ‘1392’ population.

**Supplementary Table S2.** QTLs detected in the ‘232’ × ‘1392’ population controlling the content of selected esters based on Kruskal-Wallis (K-W) and interval mapping (IM).

**Supplementary Table S3.** Marker positions in LG 6A of the ‘232’ × ‘1392’ population.

**Supplementary Table S4.** Candidate genes in the 6A QTL interval detected in the ‘232’ × ‘1392’ population.

**Supplementary Table S5.** Relative content of selected volatile organic compounds (VOCs) in the GWAS population.

**Supplementary Table S6.** GWAS results for medium-chain esters in strawberry fruit.

**Supplementary Table S7.** Candidate genes in the 6A QTL interval detected by GWAS.

**Supplementary Table S8.** Expression levels by RNA-seq in fruits of F1 lines (HIGH and LOW) contrasting in MEC content.

**Supplementary Table S9.** KEGG pathway gene set enrichment analysis with Benjamini-Hochberg (BH) adjustment method.

**Supplementary Table S10.** Mean ester values in fruit control samples and samples overexpressing *FaCCR1* and *FaOCT4* candidate genes.

**Supplementary Table S11.** *FaCCR1(6A)-H1* KASP genotypes in the ‘232’ × ‘1392’ population.

**Supplementary Table S12.** Similarity between different cinnamoyl-CoA reductases (CCRs) amino acid sequences.

**Supplementary Table S13.** catalytic-geometry summary of the docking validation panel.

**Supplementary Table S14.** Catalytic-geometry criteria and atom definitions used for docking evaluation.

**Supplementary Table S15.**Enzymatic reactions using either FaCCR1-GFP or GFP as negative control.

**Supplementary Table S16.** AX-184056927 KASP genotypes in the diverse collection and the ‘232’ × ‘1392’ population.

**Supplementary Table S17.** *FaOCT4(6A)* KASP genotypes in the diverse collection and the ‘232’ × ‘1392’ population.

**Supplementary Table S18.** Accessions of the IFAPA GWAS collection used in this study.

**Supplementary Table S19.** Primers used in this work.

**Supplementary Data Set 1.** ANOVA and/or T-test results providing variables, parameters, degrees of freedom, and test statistics.

## Data availability

The data underlying this article are available in the article and in its online supplementary material.

