## Supplementary Figures for "Differential expression and novel activity of FaCCR1 regulate medium-chain ester content and strawberry fruit aroma"

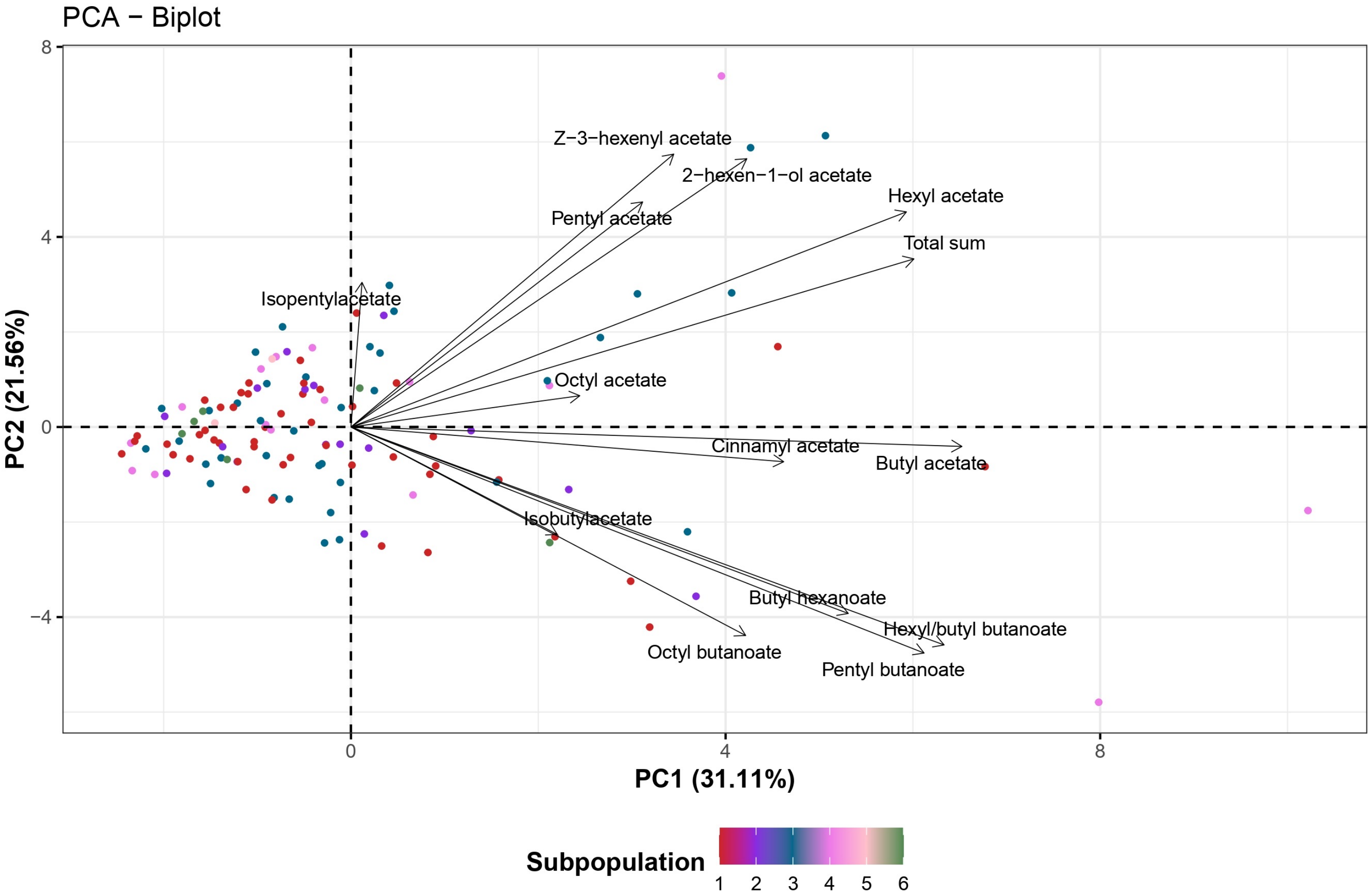


**Supplementary Figure S1.** Principal Component Analysis (PCA) biplot revealing a large diversity in medium-chain ester content in the GWAS population. Genotypes are shown by dots, while arrows represent the trait effect. The contribution of each PC is shown in the axis labels. Accessions are color-coded according to their membership to the six subpopulations described by Muñoz et al., 2024. Supports Figure 2.

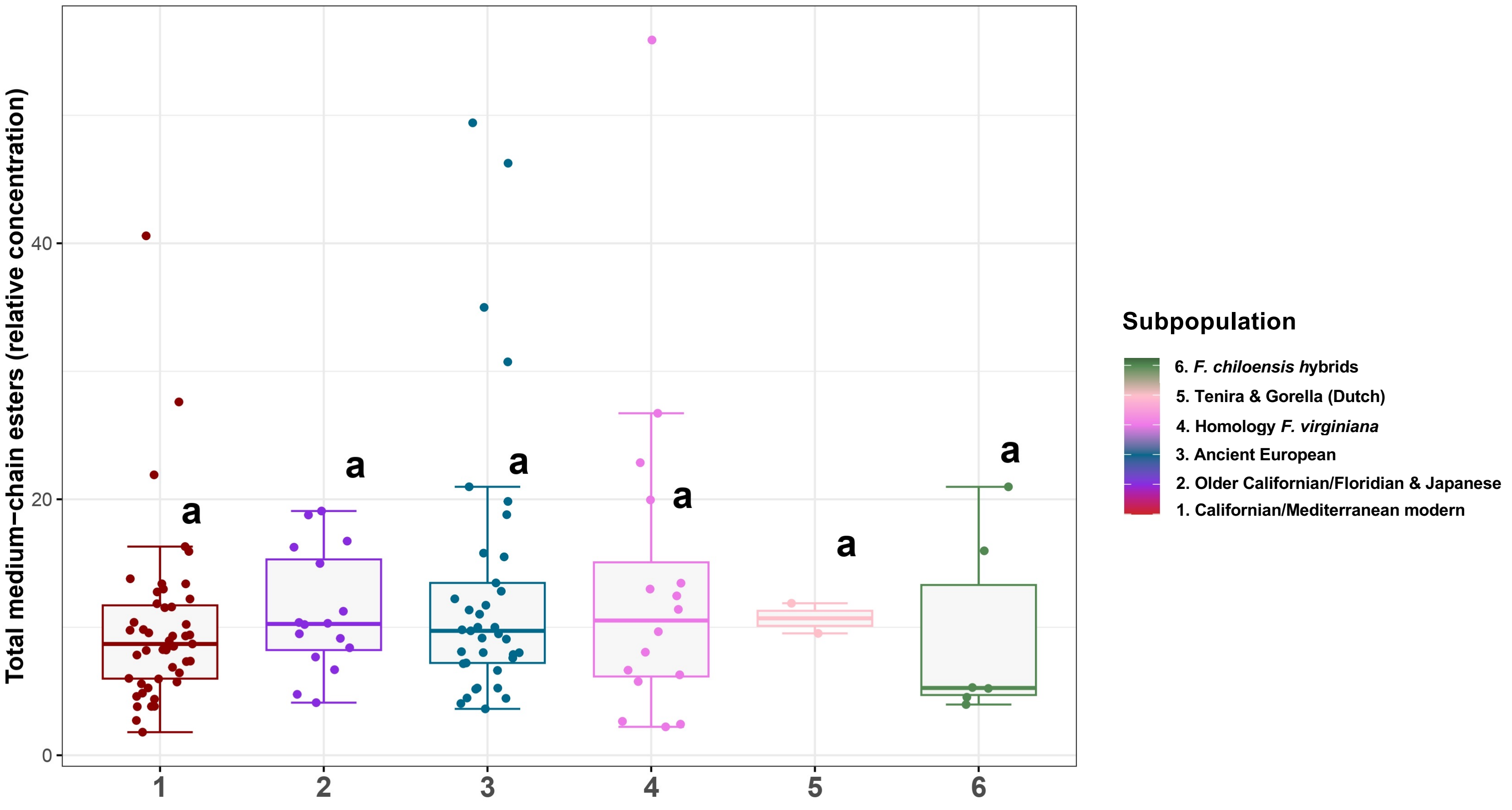


**Supplementary Figure S2.** Distribution of total medium-chain ester content in the six subpopulations from the IFAPA GWAS population, described by Muñoz et al., 2024. X-axis represents the subpopulations detailed in the legend, while dots indicate the total medium-chain phenotype (y-axis) of each accession. Boxes span the 25th and 75th percentiles and the middle line represents the median Whiskers extend to the minimum and maximum data points within 1.5 times the interquartile range (IQR). Letters denote statistically significant differences (*p*-value < 0.01) between groups, determined by one-way ANOVA with *post-hoc* Tukey HSD test for multiple comparisons. Supports Figure 2.

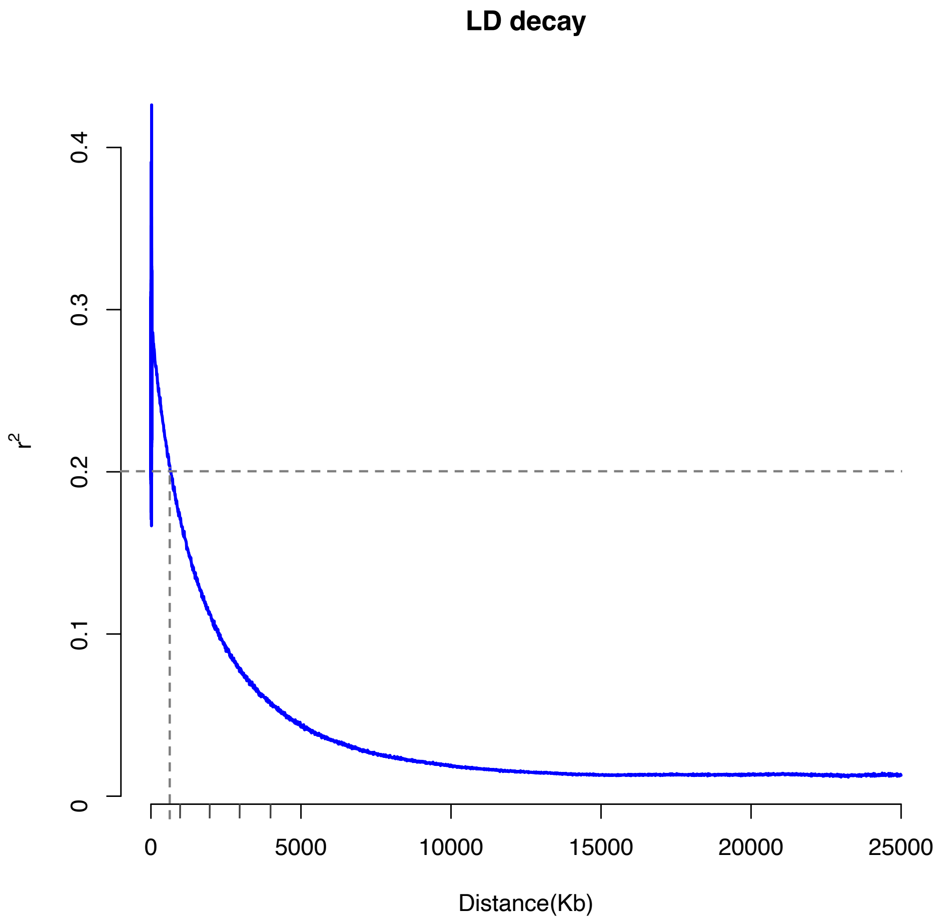


**Supplementary Figure S3.** Average linkage disequilibrium (LD) decay ($r^2$) in the strawberry genome, computed using genotypic data of 40,808 SNPs from the FanaSNP 50K array (Hardigan et al., 2020) and 124 accessions from the IFAPA GWAS population. Decay of LD across the genome is represented by the blue line, reflecting the genetic recombination and diversity within the strawberry population. Supports Figure 2.

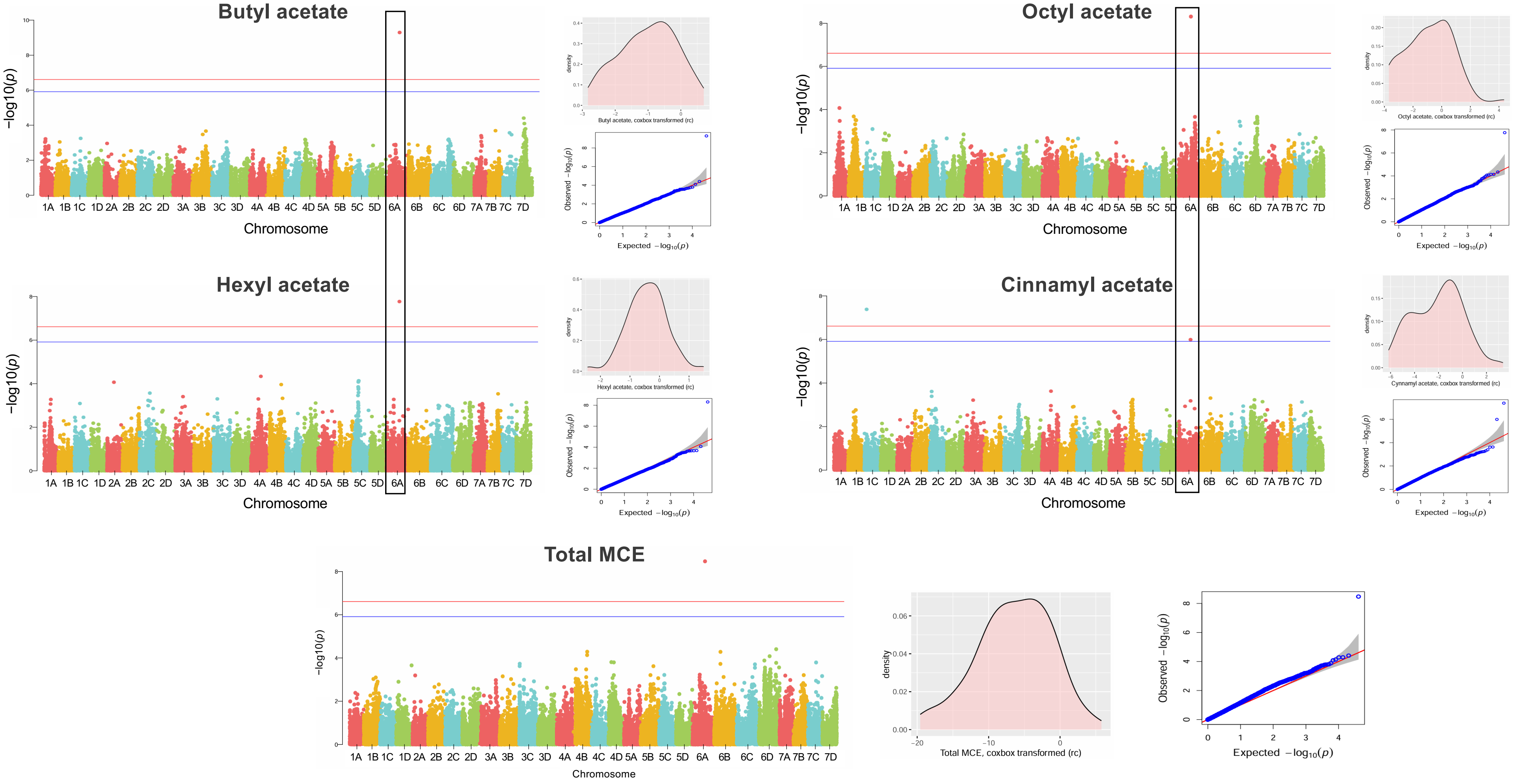


**Supplementary Figure S4.** Significant marker-trait associations for MCEs detected by GWAS using the BLINK multi-*locus* model. SNPs are represented by a dot and chromosomes are color coded according to their subgenomic origin based on the 'Royal Royce' reference genome (Hardigan et al., 2021a). The red and blue horizontal bars indicate the significance threshold at -log10 (0.01/total SNPs) and -log10 (0.05/total SNPs), respectively. Significant associations detected for the same SNP across different compounds are highlighted with black rectangles. Density plots of phenotypic distribution and QQ-plots evaluating the GWAS models are shown at right. Supports Figure 2.

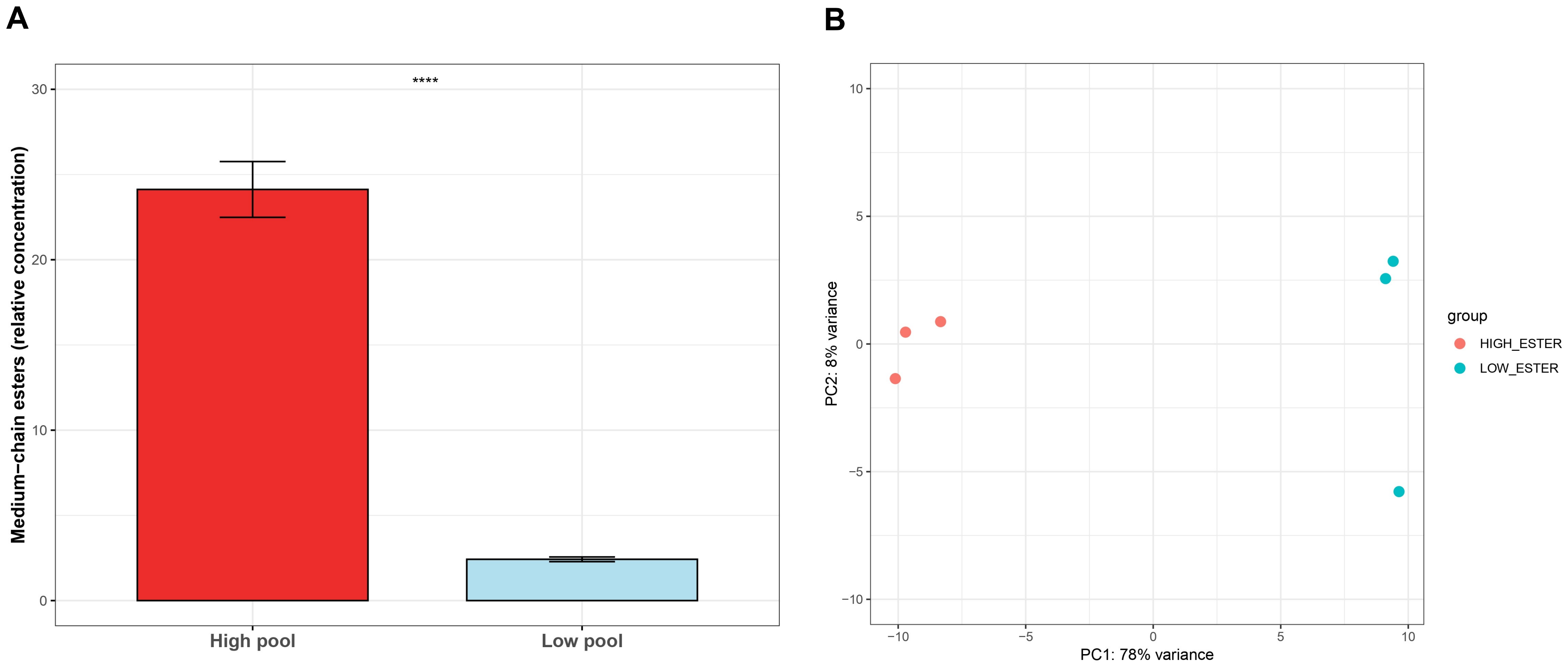


**Supplementary Figure S5. A)** Total medium-chain ester (MCE) content in the bulked pools contrasting in ester concentration used for RNA-Seq. Bars indicate the mean values of 10 $F_1$ lines over 3 seasons and whiskers show SEM. Asterisks denote statistically significant differences ≤ 0.0001 determined Wilcox test. **B)** Principal component analysis (PCA) of expressed genes in the three biological replicates of pools with high (red dots) and low (blue) MCE content. The axis labels indicate the variance explained by principal component 1 (x axis) and principal component 2 (y axis). Supports Figure 3.

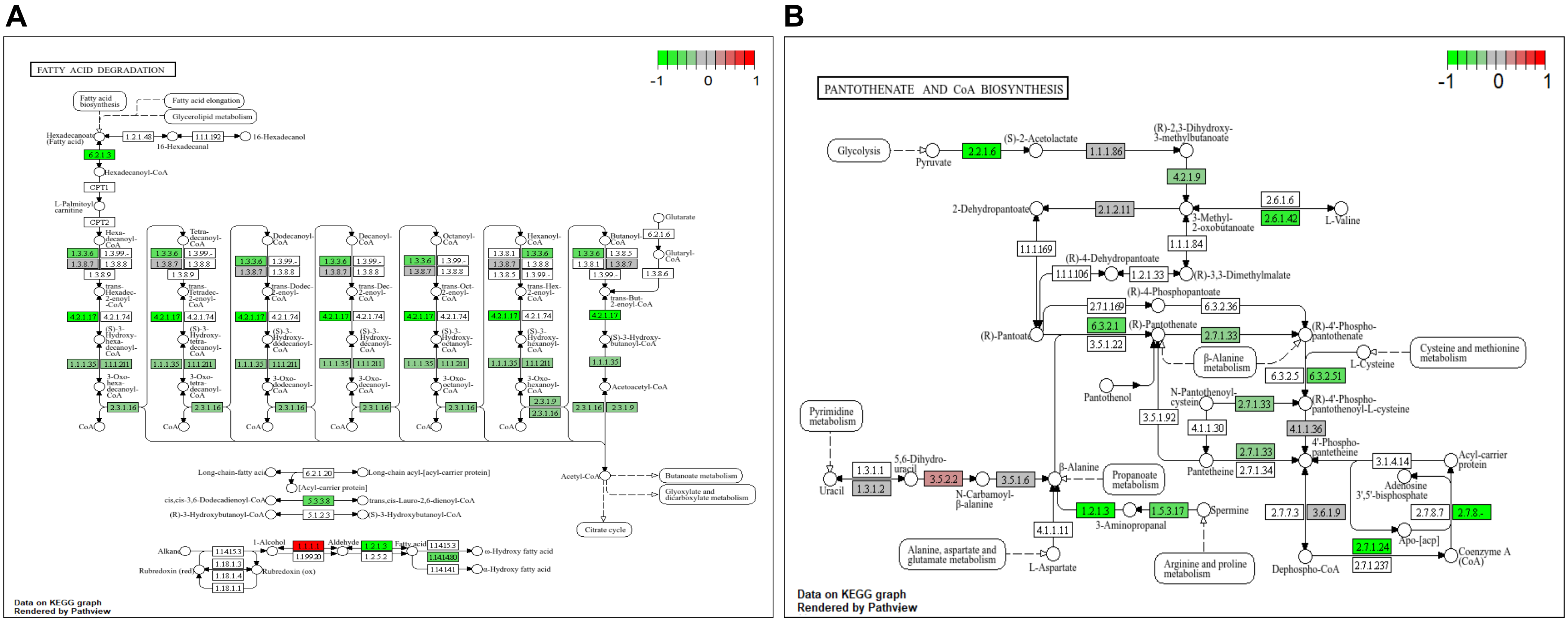


**Supplementary Figure S6.** KEGG pathway visualizations generated using Pathview for **A)** fatty acid degradation (fve00071), **B)** panthoneate and CoA biosynthesis (fve00070), **C)** butanoate metabolism (fve00650)**, D)** terpenoid biosynthesis (fve00900), **E)** linoleic acid metabolism (fve00592) and **F)** phenylpropanoid biosynthesis (fve00940). These diagrams are based on standardized KEGG pathway templates, with differentially expressed genes (DEGs) from this study overlaid onto the pathways using a color scale from green (upregulation in the high MCE pool) to red (downregulation). Arrows indicate reactions catalyzed by KEGG-annotated enzymes. Supports Figure 3.

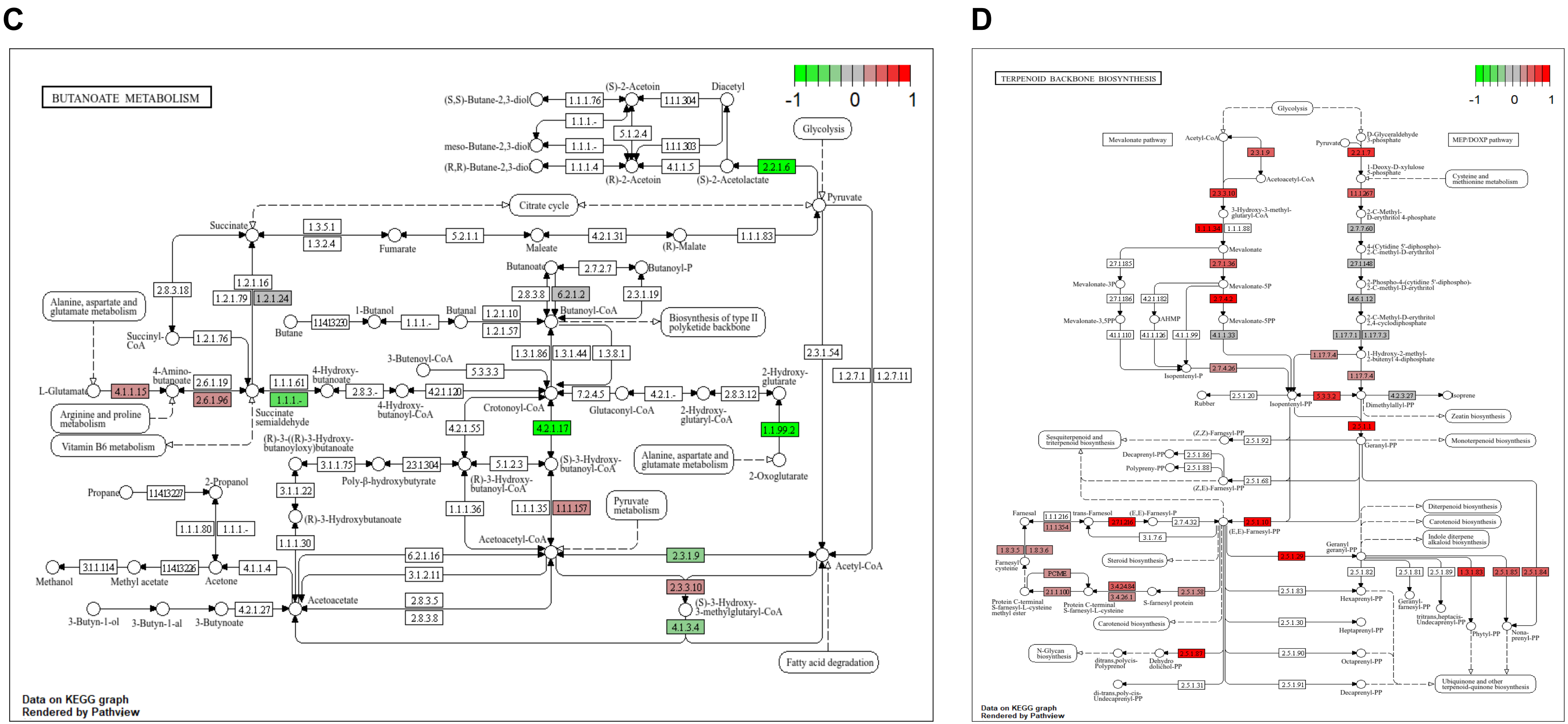


**Supplementary Figure S6.** KEGG pathway visualizations generated using Pathview for **A)** fatty acid degradation (fve00071), **B)** panthoneate and CoA biosynthesis (fve00070), **C)** butanoate metabolism (fve00650)**, D)** terpenoid biosynthesis (fve00900), **E)** linoleic acid metabolism (fve00592) and **F)** phenylpropanoid biosynthesis (fve00940). These diagrams are based on standardized KEGG pathway templates, with differentially expressed genes (DEGs) from this study overlaid onto the pathways using a color scale from green (upregulation in the high MCE pool) to red (downregulation). Arrows indicate reactions catalyzed by KEGG-annotated enzymes. Supports Figure 3.

E

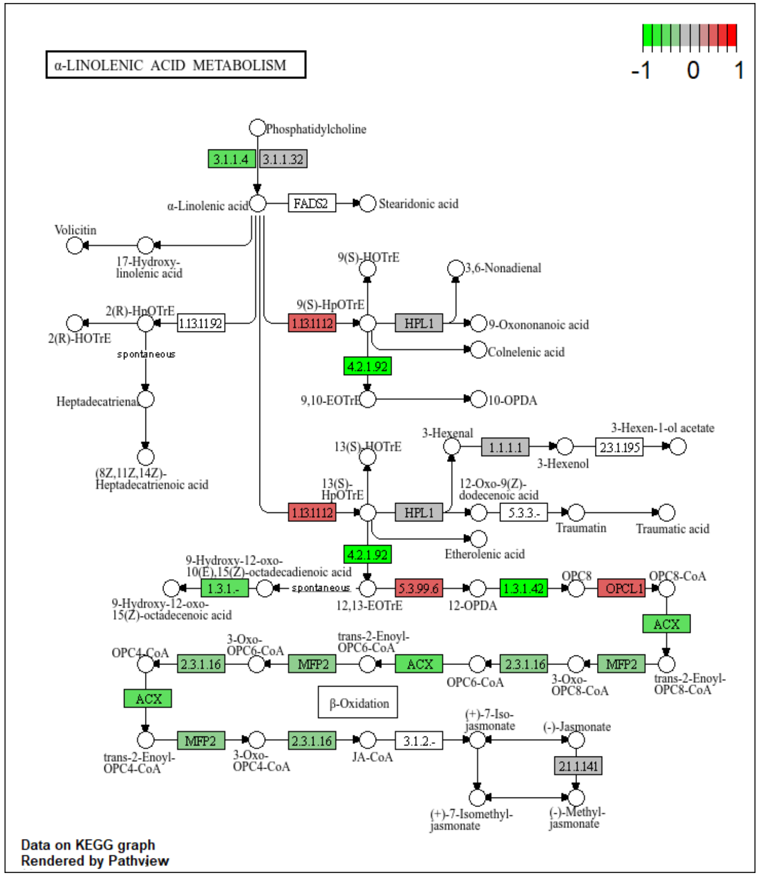


F

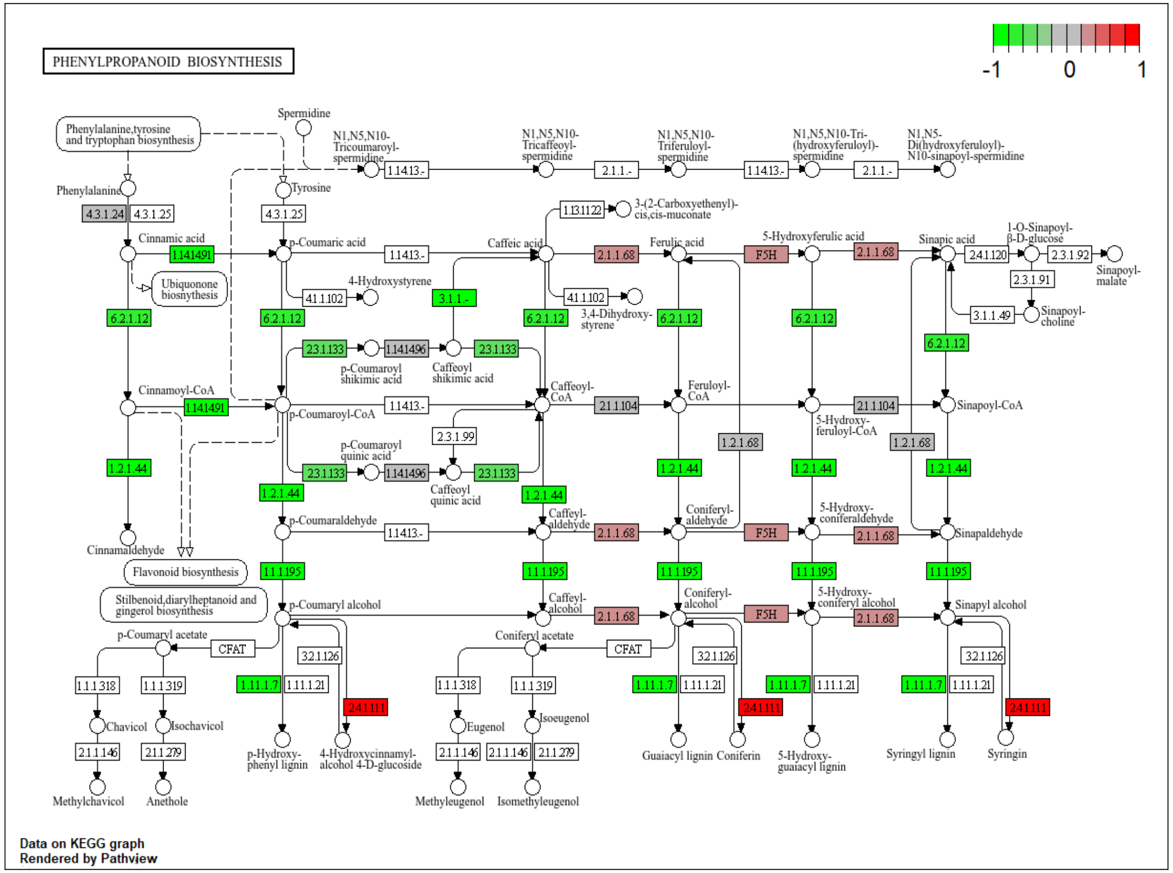


**Supplementary Figure S6.** KEGG pathway visualizations generated using Pathview for **A)** fatty acid degradation (fve00071), **B)** panthoneate and CoA biosynthesis (fve00070), **C)** butanoate metabolism (fve00650)**, D)** terpenoid biosynthesis (fve00900), **E)** linoleic acid metabolism (fve00592) and **F)** phenylpropanoid biosynthesis (fve00940). These diagrams are based on standardized KEGG pathway templates, with differentially expressed genes (DEGs) from this study overlaid onto the pathways using a color scale from green (upregulation in the high MCE pool) to red (downregulation). Arrows indicate reactions catalyzed by KEGG-annotated enzymes. Supports Figure 3.

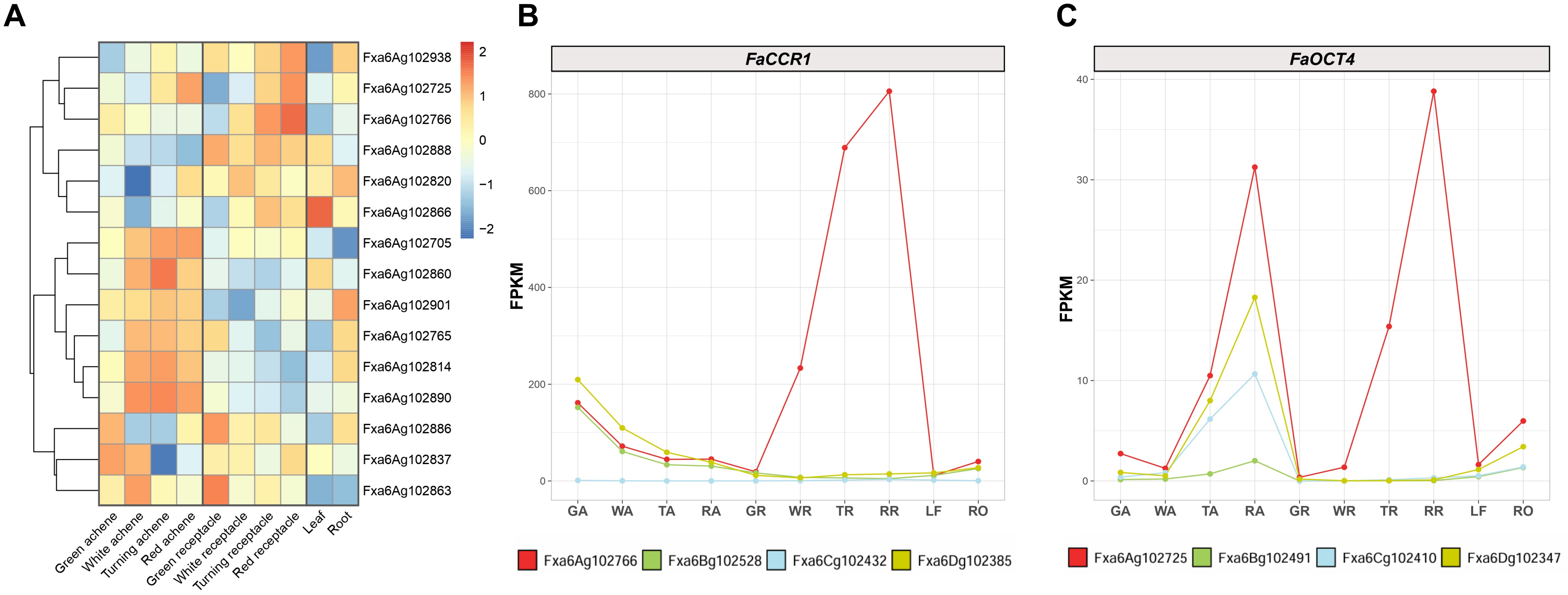


**Supplementary Figure S7. A)** Heatmap of gene expression in Z-Score transformed Fragments Per Kilobase Million (FPKM) values of the 15 DEGs within the *qMCE-6A* QTL across tissues and ripening stages of 'Camarosa'. Changes in gene expression are represented in a scale from blue (down-regulation) to red (up-regulation). **B)** Mean FPKM values of *FaCCR1(6A)* and **(C)** *FaOCT4(6A)* compared with their homoeologs in different tissues and ripening stages of 'Camarosa'. Green receptacle (GR), white receptacle (WR), turning receptacle (TR), red receptacle (RR), green achene (GA), white achene (WA), turning achene (TA), red achene (RA), leaf (L) and root (R). Gene expression estimates were extracted from publicly available data from Sánchez-Sevilla et al., (2015). Supports Figure 3.

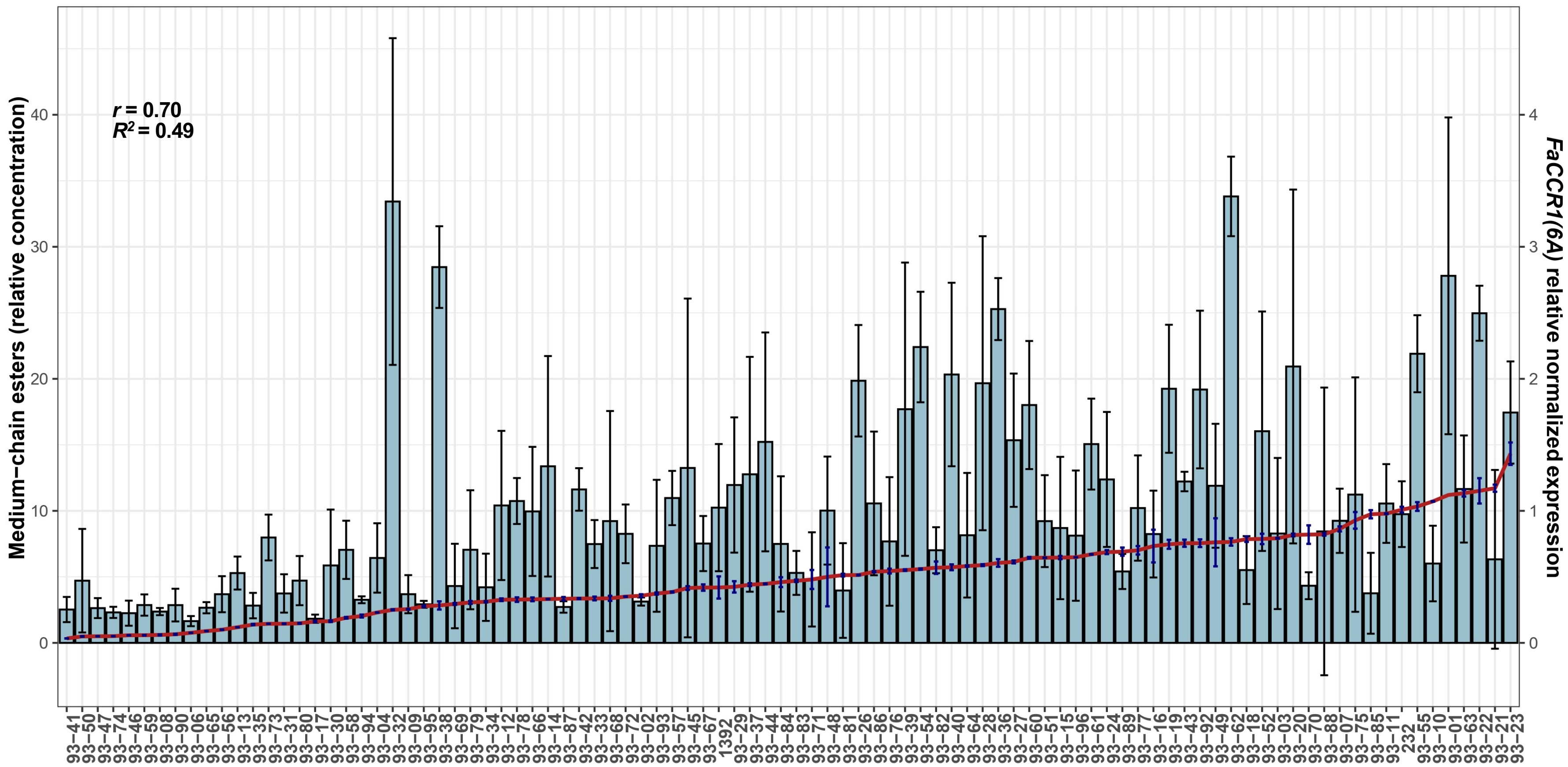


**Supplementary Figure S8.** Positive correlation between *FaCCR1(6A)* expression by RT-qPCR and total MCE concentration in the '232' × '1392' population. Bars indicate the mean MCE content from two biological replicates across two seasons, while black whiskers show the standard error. The red line represents the mean expression of *FaCCR1(6A)* measured in three technical replicates per line, with standard errors indicated by blue whiskers. Pearson correlation coefficient (*r)* and r-squared value from $\log_{10}$-transformed data are displayed in the upper left. Supports Figure 3.

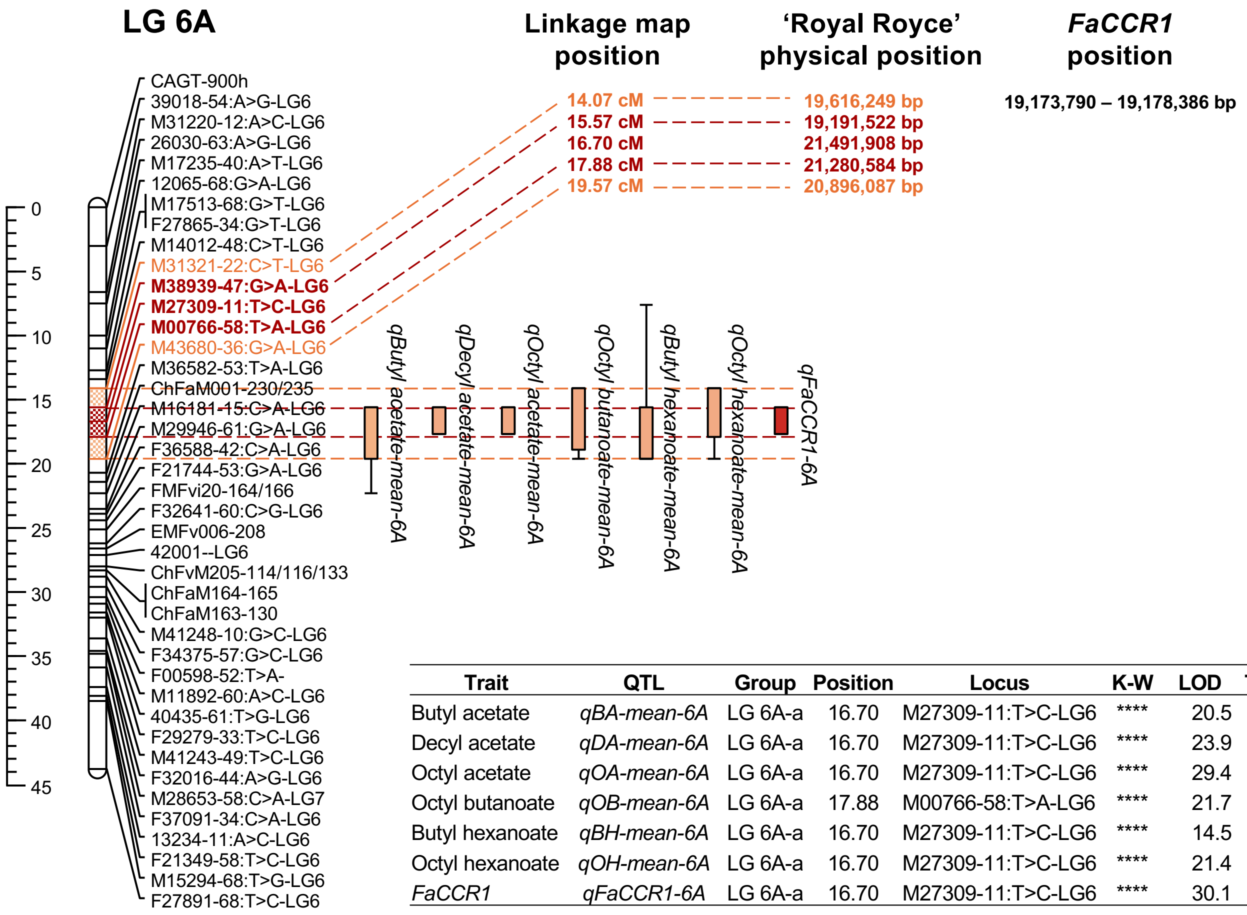


| Trait | QTL | Group | Position | Locus | K-W | LOD | Thrs. | mu_ac{00} | mu_ad{00} | mu_bc{00} | mu_bd{00} | Variance | PVE |
|---|---|---|---|---|---|---|---|---|---|---|---|---|---|
| Butyl acetate | *qBA-mean-6A* | LG 6A-a | 16.70 | M27309-11:T>C-LG6 | **** | 20.5 | 5.2 | 0.426 | 1.937 | 2.686 | 3.789 | 0.270 | 43.7 |
| Decyl acetate | *qDA-mean-6A* | LG 6A-a | 16.70 | M27309-11:T>C-LG6 | **** | 23.9 | 5.2 | 0.160 | 0.635 | 0.766 | 1.953 | 0.277 | 48.7 |
| Octyl acetate | *qOA-mean-6A* | LG 6A-a | 16.70 | M27309-11:T>C-LG6 | **** | 29.4 | 5.3 | 0.098 | 0.570 | 0.703 | 1.312 | 0.209 | 50.6 |
| Octyl butanoate | *qOB-mean-6A* | LG 6A-a | 17.88 | M00766-58:T>A-LG6 | **** | 21.7 | 5.1 | 0.039 | 0.236 | 0.363 | 0.597 | 0.392 | 45.1 |
| Butyl hexanoate | *qBH-mean-6A* | LG 6A-a | 16.70 | M27309-11:T>C-LG6 | **** | 14.5 | 5.2 | 0.151 | 0.763 | 0.904 | 1.732 | 0.625 | 46.3 |
| Octyl hexanoate | *qOH-mean-6A* | LG 6A-a | 16.70 | M27309-11:T>C-LG6 | **** | 21.4 | 5.1 | 0.041 | 0.294 | 0.507 | 0.920 | 0.554 | 47 |
| *FaCCR1* | *qFaCCR1-6A* | LG 6A-a | 16.70 | M27309-11:T>C-LG6 | **** | 30.1 | 5.2 | 0.070 | 0.405 | 0.370 | 0.587 | 0.141 | 65.2 |

**Supplementary Figure S9.** QTL analysis using the mean values of six selected MCEs in the three consecutive seasons (2007-2009) and the expression levels of *FaCCR1(6A)* in the '232' × '1392' population. A cis eQTL controlling the expression of *FaCCR1(6A)* (in red) collocated at linkage group (LG) 6A with the *qMCE-6A* locus indicating that the differential expression of this gene likely underlies the variation observed in MCEs. QTL are shown as colored boxes (1-LOD interval) to the right of the LG, with error bars showing the 2-LOD interval. Red and orange dashed horizontal lines define the 1-LOD intervals for the eQTL and MCEs, respectively. Marker positions in genetic (cM) and physical map ('Royal Royce') are indicated, as well as the physical position of *FaCCR1(6A)*. The table summarizes detected QTL based on Kruskal-Wallis (K-W) and interval mapping (IM). The position (in cM) of the most closely associated marker is indicated. The estimated mean effect (for raw data) of the QTL (mu) associated with each genotype (ac, ad, bc, bd; for the ab x cd configuration) with phase type {00} is shown as well as the variance and percentage of variance explained (PVE). Supports Figure 3.

**A**

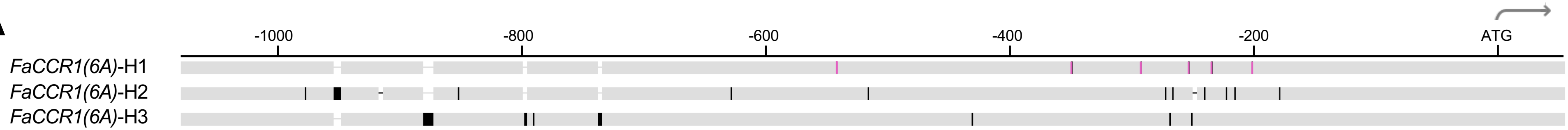


| Position from ATG | -950 | -928 | -896 | -862 | -841 | -789 | -783 | -732 | -628 | -543 | -518 | -435 | -355 | -299 | -279 | -276 | -274 | -261 | -259 | -256 | -248 | -242 | -231 | -224 | -210 | -188 |
|---|---|---|---|---|---|---|---|---|---|---|---|---|---|---|---|---|---|---|---|---|---|---|---|---|---|---|
| *FaCCR1(6A)*-H1 | A | C------ | T | A-------- | T | A-- | C | A---- | C | A | T | C | T | C | C | A | G | A | C | T | C | A | A | C | G | A |
| *FaCCR1(6A)*-H2 | C | CTATTTC | - | A-------- | A | A-- | C | A---- | G | G | C | C | C | T | T | A | A | G | C | - | T | G | C | A | A | T |
| *FaCCR1(6A)*-H3 | A | C------ | T | ATATTGGAT | T | AAA | T | AGTAC | C | G | T | T | C | T | C | G | G | G | T | T | C | G | A | C | A | A |

**B**

| Line | Genotype | *FaCCR1(6A)* expression | KASP AX-184056927 | KASP *FaOCT4(6A)* |
|---|---|---|---|---|
| 232 | *H1:H3* | Intermediate | A:A | G:A |
| 1392 | *H1:H2* | Intermediate | A:G | G:A |
| 93-22 | *H1:H1* | High | A:A | G:G |
| 93-36 | *H1:H1* | High | A:A | G:G |
| 93-62 | *H1:H1* | High | A:A | G:G |
| 93-42 | *H1:H3* | Intermediate | A:A | G:A |
| 93-89 | *H1:H2* | Intermediate | A:G | G:A |
| 93-46 | *H2:H3* | Low | A:G | A:A |
| 93-74 | *H2:H3* | Low | A:G | A:A |
| 93-90 | *H2:H3* | Low | A:G | A:A |

**C**

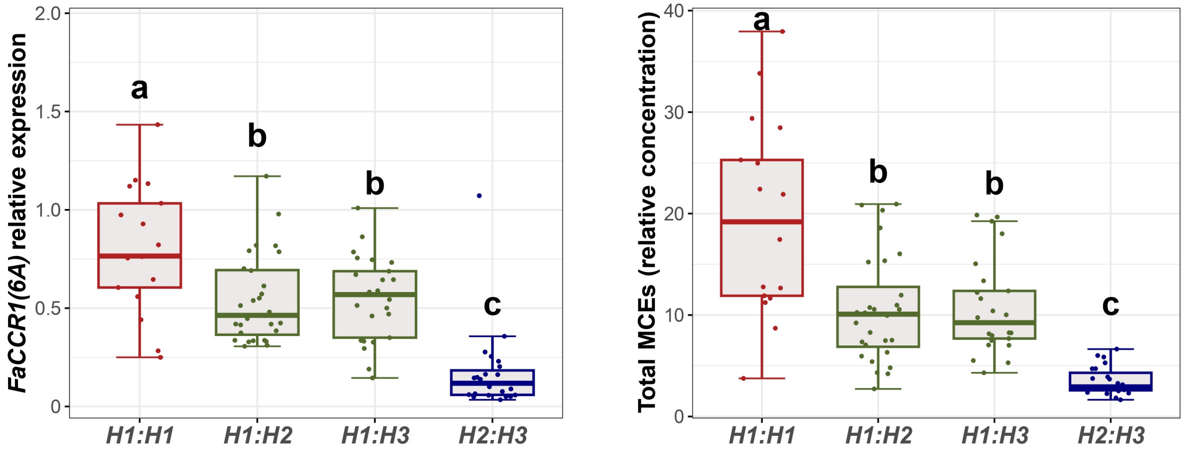


**Supplementary Figure S10. A)** Schematic alignment of the 1,147 bp fragment corresponding to the 5' upstream regulatory region of *FaCCR1(6A)* from the three haplotypes (*H1-H3*) identified in the '232' × '1392' population. Numbering indicates the position relative to the *FaCCR1(6A)* ATG start codon according to the 'Royal Royce' genome. Vertical lines indicate polymorphic positions among haplotypes, and the *H1*-specific polymorphisms are highlighted in pink. The table below summarizes all variants detected at each polymorphic position, with the six H1-specific SNPs highlighted in pink. **B)** Table showing the *FaCCR1(6A)* genotypes determined by full-length amplicon sequencing using the ONT platform, together with *FaCCR1(6A)* expression levels and the allele calls obtained with the KASP assays targeting AX-184056927 and *FaOCT4(6A)*. *H1* haplotype was associated with high *FaCCR1(6A)* expression while *H2* and *H3* haplotypes conferred low expression. **C)** Allelic effect of FaCCR1-H1 KASP on *FaCCR1(6A)* expression and total MCE content. Boxes span the 25th and 75th percentiles and the middle line represents the median. Whiskers extend to the minimum and maximum data points within 1.5 times the interquartile range (IQR). Letters denote statistically significant differences (*p*-value < 0.05) between groups, determined by ANOVA with *post-hoc* Tukey-HSD test for multiple comparisons. Supports Figure 3.

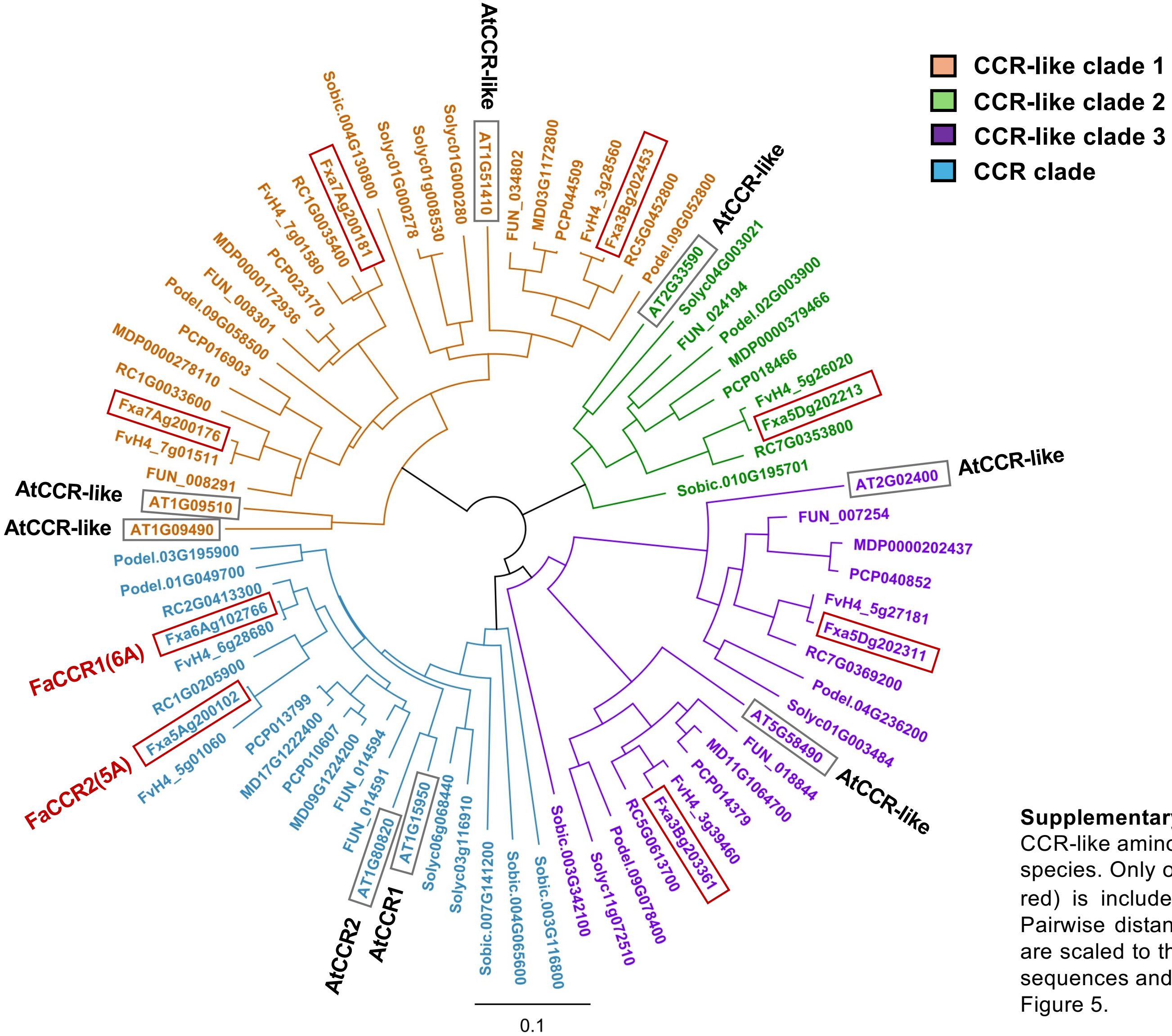


**Supplementary Figure S11.** Neighbour-joining phylogenetic tree of CCR and CCR-like amino acid sequences from strawberry (*Fragaria* × *ananassa)* and other species. Only one homoeologous copy from each strawberry gene (highlighted in red) is included. The orthologous gene in *F. vesca* is also included (FvH4). Pairwise distances were computed using Jukes-Cantor method. Branch lengths are scaled to the amount of evolutionary change. Percentage of identity between sequences and species names are shown in Supplementary Table S12. Supports Figure 5.

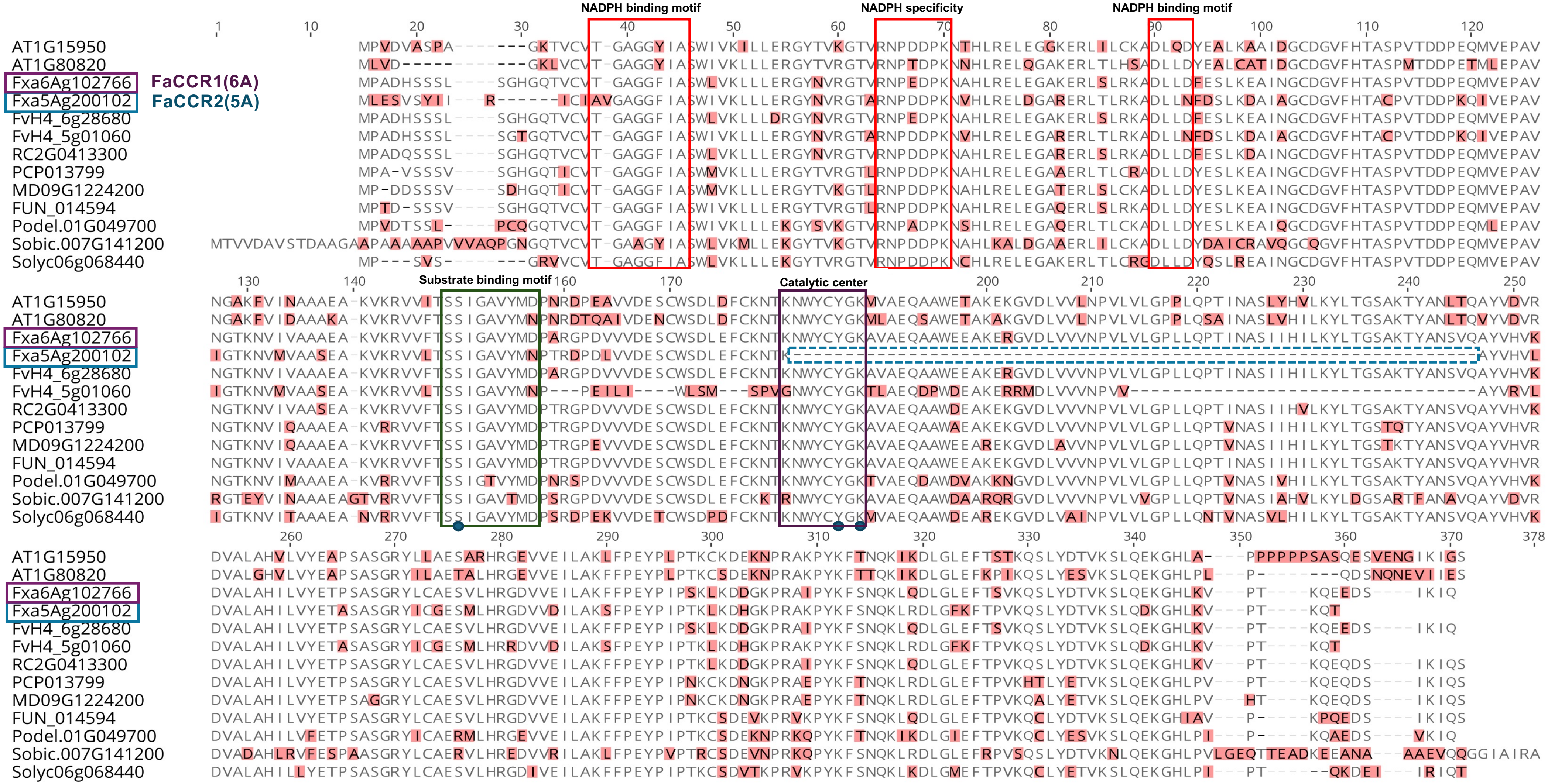


**Supplementary Figure S12.** Multiple alignment of cinnamoyl-CoA reductase amino acid sequences from strawberry (*Fragaria x ananassa)* and other species generated by Geneious v2021.2.2. Disagreements are highlighted in red. Conserved motifs and the catalytic triad are indicated with a box or circle, respectively, according to Barakat et al. (2011) and Chao et al. (2017). Strawberry FaCCR1(6A) and FaCCR2(5A) are underlined, along with the deletion in the catalytic center. Supports Figure 5.

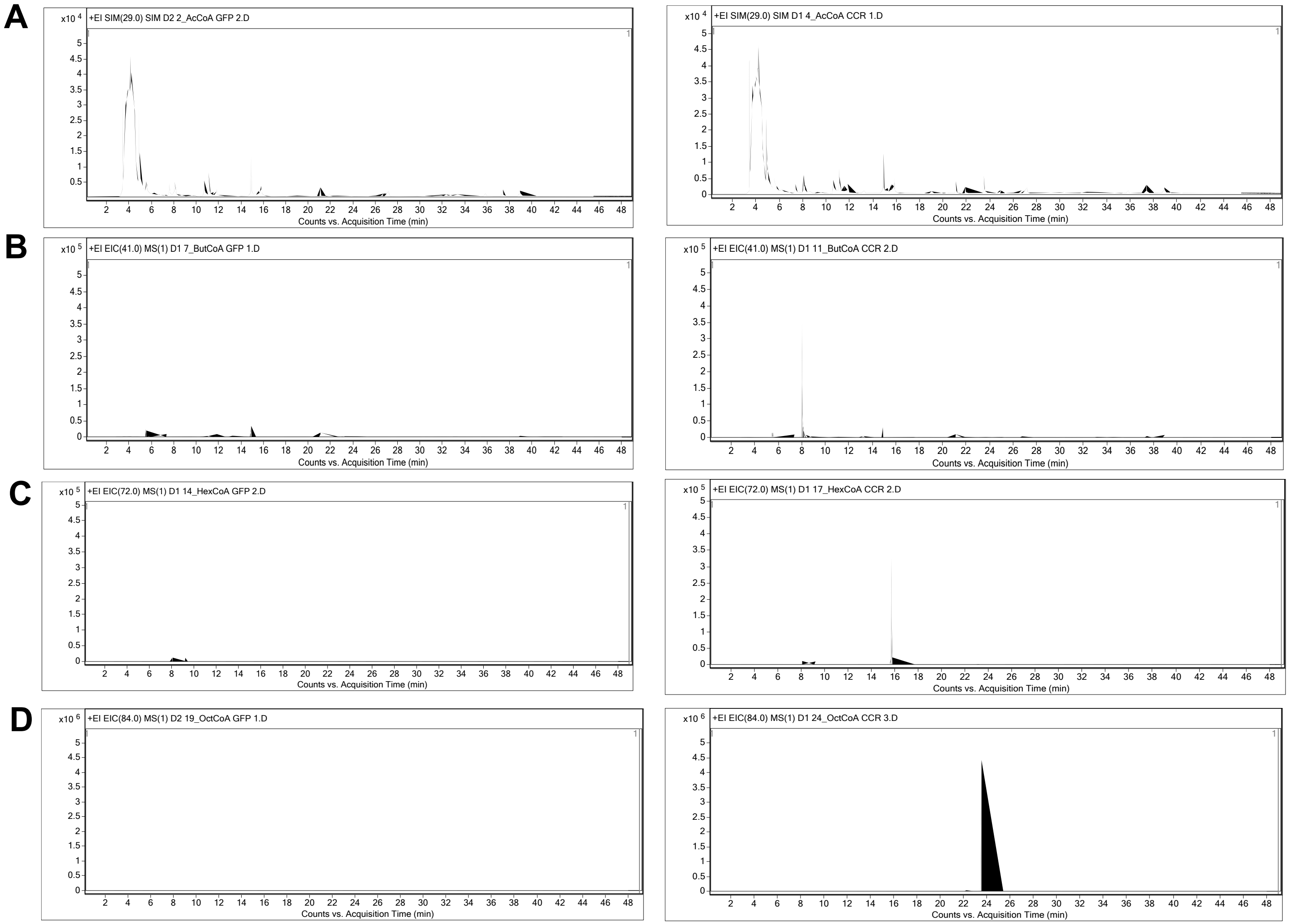


**Supplementary Figure S13**: Extracted ion chromatograms of a characteristic ion (Q Ion, *m/z*) of aldehydes resulting from enzymatic reactions with GFP as blank control (left) or FaCCR1-GFP (right). Reactions were performed using acetyl-CoA **(A)** as negative control or the test substrates butanoyl-CoA **(B)**, hexanoyl-CoA **(C)** and octanoyl-CoA **(D)**. Peaks at the corresponding retention time are observed in reactions with FaCCR1-GFP. Supports Figure 6.

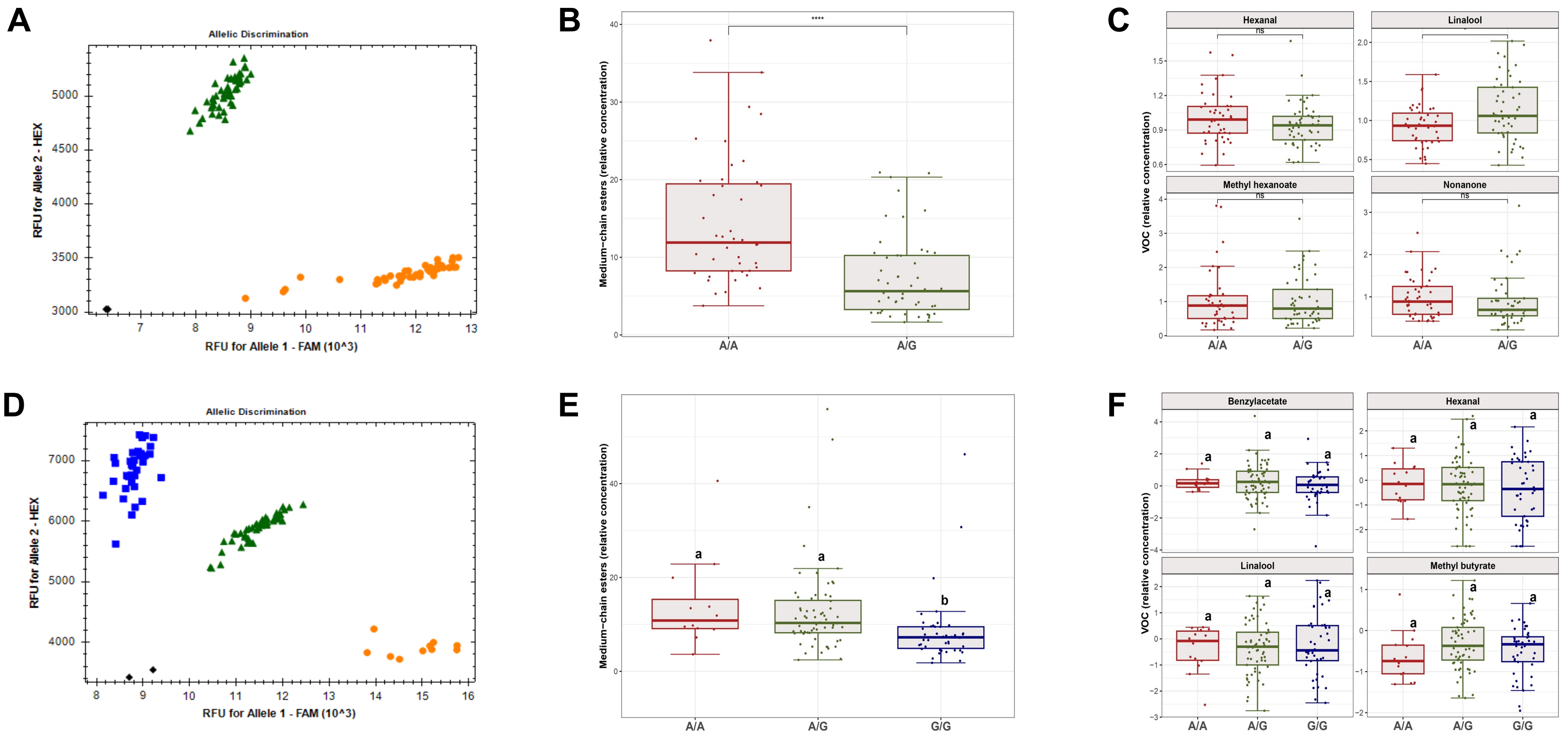


**Supplementary Figure S14.** Evaluation of a KASP assay in SNP AX-184056927 for fruit MCE prediction. **A)** Example of genotype clusters for KASP AX-184056927 in the '232' × '1392' population. The reference G allele is labelled with HEX (blue color) and the alternative A allele is labelled with FAM (yellow color). Heterozygous samples are represented in green and non-template controls or no calls in black. RFU, relative fluorescence units for HEX and FAM fluorescent dyes. **B)** Box-plot of KASP AX-184056927 allelic marker effects on total MCE content in the '232' × '1392' mapping population ($n$ = 96) and **C)** on other VOCs not controlled by *qMCE-6A*. **D**) Genotype clusters in the 124 diverse accessions from the IFAPA GWAS collection. **E)** Box-plot of allelic marker effects on total MCE content in the GWAS collection evaluated in 2020-21, and **F)** in other VOCs. Boxes span the 25th and 75th percentiles and the middle line represents the median. Whiskers extend to the minimum and maximum data points within 1.5 times the interquartile range (IQR). Letters denote statistically significant differences ($p$-value < 0.05) between groups, determined by one-way Welch's ANOVA with *post-hoc* Games-Howell test for multiple comparisons. Asterisks show statistically significant differences (**** $p$-value ≤ 0.0001; ns, not significant) using Wilcox test for non-parametric data or t-test for parametric data). Supports Figure 7.
